# Elucidation of gene clusters underlying withanolide biosynthesis in ashwagandha through yeast metabolic engineering

**DOI:** 10.1101/2024.12.24.630284

**Authors:** Erin Reynolds, Marena Trauger, Fu-Shuang Li, Jonathan Huang, Trevor Moss, Bastien Christ, Menglong Xu, Eva Knoch, Jing-Ke Weng

**Affiliations:** Institute for Plant-Human Interface, Northeastern University, Boston, MA 02115, USA; Department of Chemical Engineering, Massachusetts Institute of Technology, Cambridge, MA 02139, USA; Department of Chemistry and Chemical Biology, Department of Bioengineering, and Department of Chemical Engineering, Northeastern University, Boston, MA 02115, USA; Whitehead Institute for Biomedical Research, Cambridge, MA 02142, USA; Department of Electrical Engineering and Computer Science, Massachusetts Institute of Technology, Cambridge, MA 02139, USA; Department of Biomedical Engineering, Boston University, Boston, MA 02215, USA; RIKEN Center for Sustainable Resource Science, Yokohama, Kanagawa 230-0045, Japan

## Abstract

Withanolides are medicinally relevant steroidal lactones produced by *Withania somnifera* (ashwagandha) amongst other Solanaceae family plants. However, the biosynthetic pathway to withanolides is largely unknown, preventing scale-up and hindering pharmaceutical applications. We sequenced the genome of *W. somnifera* and identified two biosynthetic gene clusters exhibiting a segmented tissue-specific expression pattern. We characterized the cluster enzymes through stepwise pathway reconstitution in yeast and transient expression in *Nicotiana benthamiana*, leading to the identification of three cytochrome P450s (CYP87G1, CYP88C7, and CYP749B2) and a short-chain dehydrogenase that produce a lactone ring-containing intermediate when co-expressed. A fourth cytochrome P450 (CYP88C10) and a sulfotransferase convert this into an intermediate with the characteristic withanolide A-ring structure, featuring a C_1_ ketone and C_2_-C_3_ unsaturation. The discovery of the sulfotransferase as a core pathway enzyme challenges the conventional paradigm of sulfotransferases as tailoring enzymes. These insights pave the way for an efficient biomanufacturing process for withanolides and future development of withanolide-derived drugs.

## Introduction

*Withania somnifera*, commonly known as ashwagandha, has been used for over 3,000 years in Indian Ayurvedic traditional medicine to treat ailments including tuberculosis and inflammation^1^. In recent years, ashwagandha has gained popularity as a dietary supplement used for stress reduction and sleep improvement^2^. Clinical trials have shown *W. somnifera* extracts to be effective for treating depression and anxiety^3,4^, as well as improving cardiorespiratory endurance, memory and cognition^5,6^. The main bioactive compounds in *W. somnifera* are steroidal lactones known as withanolides^7^. Withanolides possess a range of pharmacological properties including anti-cancer, anti-inflammatory, neurological, and immunomodulatory activities^7,8^. The core withanolide structure is a C_28_ ergostane skeleton with C_26_ and either C_22_ or C_23_ oxidized to form a δ- or γ-lactone ring^7^ (Fig 1d, Supplementary Fig. 1). The two other characteristic chemical features are a C_1_ ketone, possessed by over 90% of withanolides^9^, and A-ring C_2_- C_3_ unsaturation^9,10^. There are over 550 reported withanolides^10^ and over 130 withanolides produced by *W. somnifera* alone^11^ (Supplemental Fig. 1). The most abundant withanolide in *W. somnifera* leaf tissue is withaferin A and the most abundant withanolide in root tissue is withanolide A (Fig. 1c, Supplementary Fig. 2). Withaferin A exhibits anti-carcinogenic effects on various cancers^12^, while withanolide A can promote regeneration of damaged neurons^13^.

**Fig. 1.**
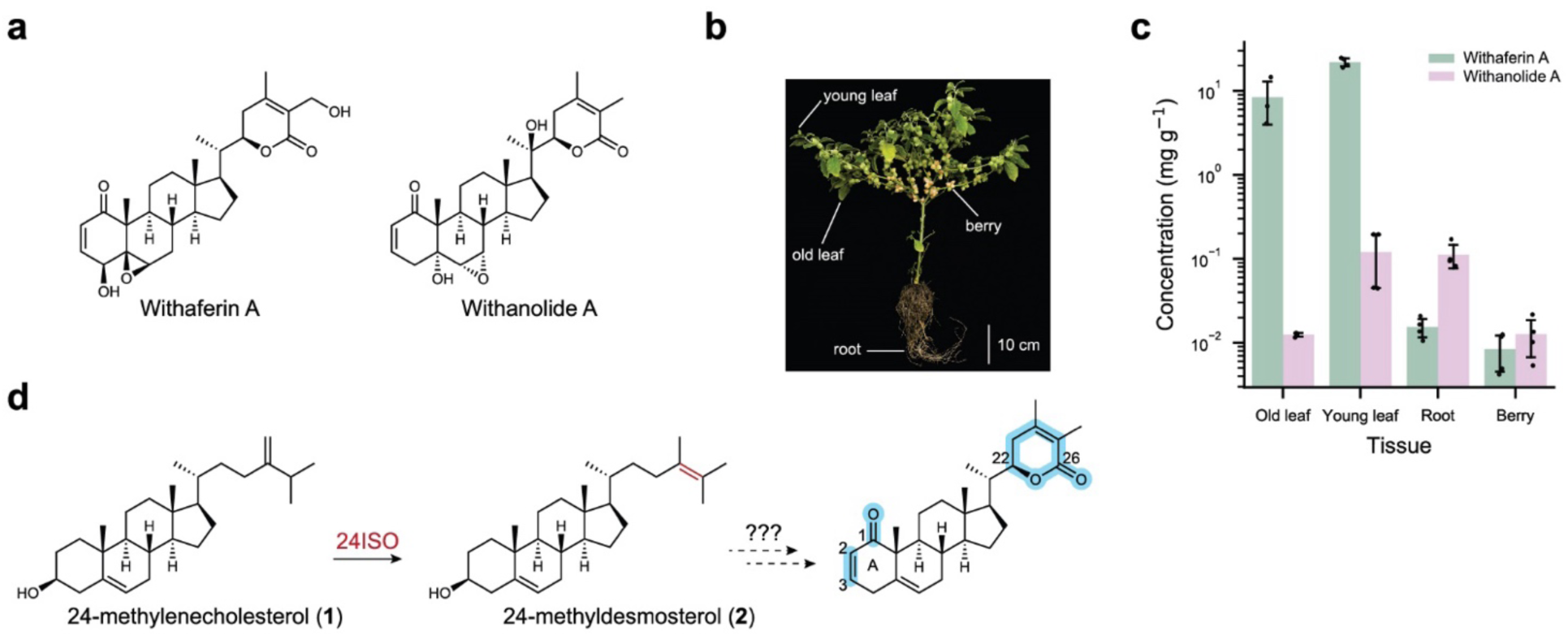
Major withanolides found in *W. somnifera*. **a**, Chemical structures of the major withanolides found in *W. somnifera*, withaferin A and withanolide A. **b**, Tissues (old leaf, young leaf, root, and berry) from *W. somnifera* plants were harvested for metabolic profiling and RNA sequencing. **c**, Withaferin A and withanolide A content of *W. somnifera* tissues. Mean values are plotted of n=4 biological replicates and error bars indicate the standard deviation. Data plotted on a log scale. **d**, 24ISO isomerizes 24- methylenecholesterol (**1**) to 24-methyldesmosterol (**2**) in the first committed step of withanolide biosynthesis. The subsequent steps required to generate the key chemical features of withanolides (lactone ring, C_1_ ketone, C_2_-C_3_ unsaturation; highlighted blue) are unknown.

Despite their promise, pharmaceutical applications are hindered by the lack of a scalable method for withanolide production. Withanolides are difficult to chemically synthesize due to their structural complexity. The first divergent chemical synthesis for withanolides was only recently reported and requires more than 14 steps^14^. Currently, isolation from plant material is the only feasible production method for withanolides. Engineering withanolide production in a heterologous host such as *Saccharomyces cerevisiae* (yeast) is a more scalable and higher-yield alternative; however, elucidation of the withanolide biosynthetic pathway is necessary for this approach.

Little is known about withanolide biosynthesis beyond the first committed step: the conversion of phytosterol pathway intermediate 24-methylene cholesterol (**1**) to 24-methyldesmosterol (**2**) by sterol Δ^24^- isomerase (*24ISO*) (Fig 1d)^15^. Previous research identified putative biosynthetic gene clusters in the genomes of several Solanaceae family plants wherein numerous 2-oxoglutarate-dependent dioxygenases (2OGDs) and cytochrome P450s (P450s) were co-located with *24ISO*^15^. We hypothesized that the genome of *W. somnifera* may also contain a biosynthetic gene cluster around *24ISO*, and that the cluster genes may be involved in withanolide biosynthesis. In this study, we sequenced the genome of *W. somnifera* and discovered two withanolide biosynthetic gene clusters. We employed a learning-by-building engineering approach to elucidate withanolide biosynthesis through stepwise pathway reconstitution in yeast, followed by validation experiments in planta. This approach identified six enzymes encoded by these gene clusters leading to the production of withanolides possessing key chemical characteristics such as the lactone ring, C_1_ ketone, and C_2_-C_3_ unsaturation. We generated yeast strains capable of milligram-scale production of these intermediates, serving as prototypes that can be further optimized and tailored for industrial production of withanolides for future pharmaceutical applications.

## Results

### Discovery of two withanolide biosynthetic gene clusters in *W. somnifera* genome

To enable biosynthetic gene cluster discovery, we sequenced the genome of *W. somnifera* using PacBio long-read sequencing and high-throughput chromosome conformation capture (Hi-C) sequencing (Methods, Supplementary Table 1, Supplementary Fig. 3). The final assembly was 98.6% complete according to Benchmarking Universal Single-Copy Orthologs (BUSCO)^16^ analysis and the largest 24 scaffolds contained 99.5% of the assembly, representing 24 pseudochromosomes. Genome annotation was informed by Illumina paired-end RNA sequencing (RNA-seq) of *W. somnifera* root, leaf, and berry tissue and protein-level data from the Viridiplantae clade, resulting in the identification of 36,057 genes.

To identify potential biosynthetic gene clusters, we performed a BLAST search against the *W. somnifera* genome using the previously published *24ISO* sequence^15^ from *W. somnifera* as the query. This search returned three genes with 99.8% (*24ISO2*), 93.5% (*24ISO1*), and 86.7% (*24ISO\**) amino acid sequence identity to the query. We discovered two putative biosynthetic gene clusters by investigating the annotated genes surrounding the hits. Cluster 1 spans 1.14 Mb on pseudochromosome 10 and contains 18 full-length genes, 16 of which belong to biosynthetically relevant enzyme families such as P450s, sulfotransferases (SULFs), acyltransferases (ACTs), and short-chain dehydrogenases (SDHs) (Fig. 2b,d). Cluster 2 spans 1.4 Mb on pseudochromosome 2 and consists of 13 full-length genes annotated as P450s and 2OGDs (Fig. 2b,d). Microsynteny analysis between cluster 1 and cluster 2 suggests that these clusters evolved from each other by way of a duplication event (Fig. 2b), most likely via whole genome duplication, as pseudochromosome 2 and pseudochromosome 10 are syntenic (Fig. 2a). Compared to other members of the Solanoideae subfamily, *W. somnifera* has experienced whole genome duplication, evidenced by intra-genome synteny (Fig. 2a, Supplementary Fig. 4) and the chromosome number of 24 instead of 12, the typical number for Solanoideae plants^17^. Reported plant biosynthetic gene clusters contain three to 15 genes and typically range in size from ∼35 kb to a few hundred kb^18–20^. The largest plant biosynthetic gene cluster by gene number is the benzylisoquinoline alkaloid cluster in *Papaver somniferum* with 15 genes^21^. Therefore, the biosynthetic gene clusters we identified are among the largest in plants discovered to date. To determine whether genes in the clusters were co-expressed, we performed RNA-seq read alignment and quantification. Surprisingly, we found that both gene clusters were split into two differentially expressed subclusters: one highly expressed in root tissue and the other highly expressed in leaf tissue (Fig. 2c,e). In cluster 1, the first six collinear genes (from *CYP87G1-1* to *SDH1)* form the root-expressed subcluster and genes from *ACT1* to *SULF1* form the leaf-expressed subcluster. In cluster 2, genes from *CYP88C8-2* to *OGD1* form the root-expressed subcluster and genes from *CYP749B2-5* to *CYP749B2-3* form the leaf-expressed subcluster, with the exception of *OGD7*, which is not highly expressed in any tissue. We corroborated the root and leaf subcluster delineations, determined by read counts, by computing pairwise Pearson correlation coefficients (Fig. 2e). Most cluster genes have at least one paralog in a leaf subcluster and at least one in a root subcluster (Supplementary Table 2). This led us to question whether the leaf- and root-expressed paralogs were functionally distinct, contributing to observed differences in withanolide profile between leaf and root (Fig 1, Supplementary Fig. 2). We characterized the paralogs of *24ISO, CYP87G1*, *CYP88C7, SDH,* and *SULF* but found no major differences in product profile (Supplementary Fig. 5).

**Fig. 2.**
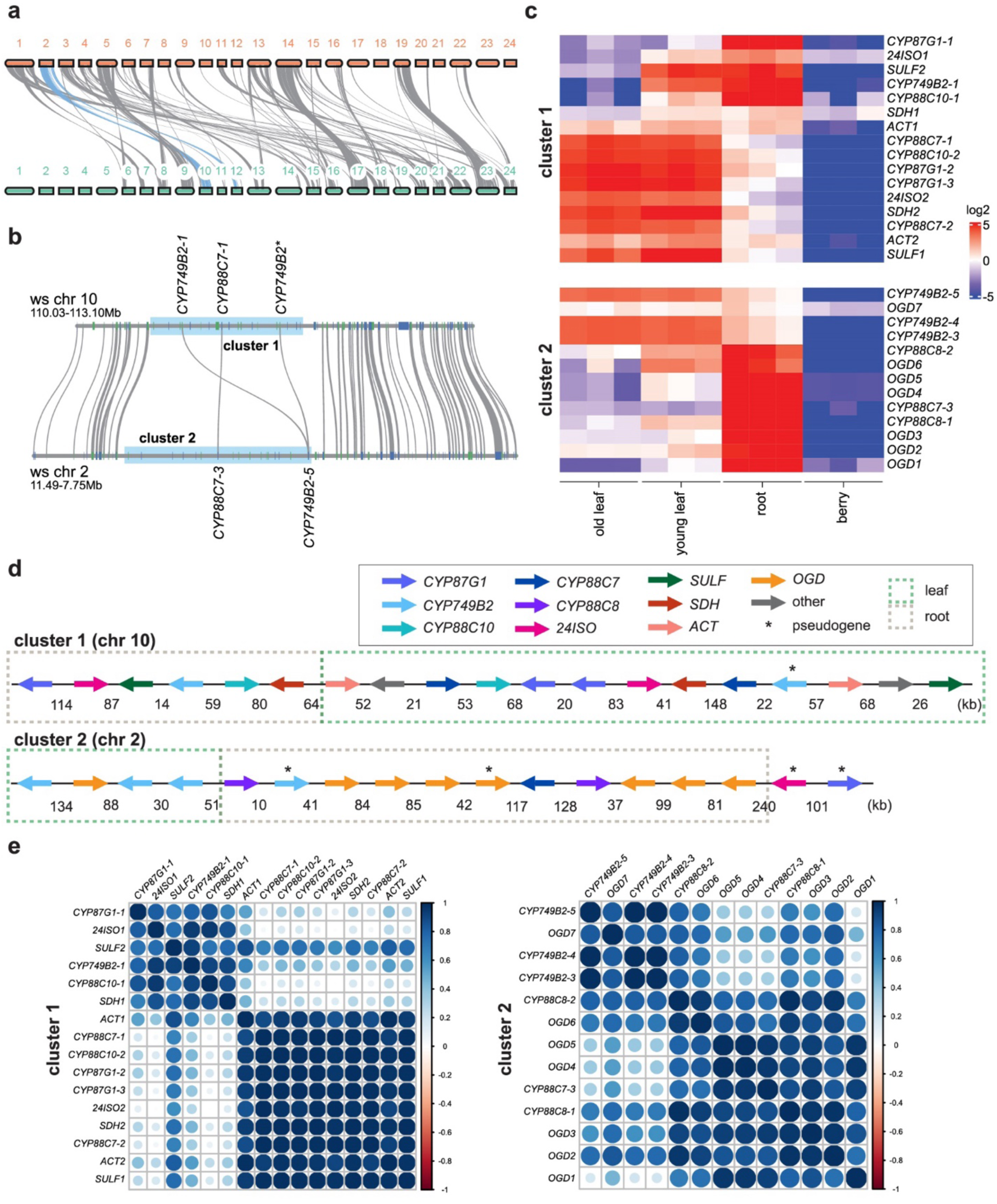
Discovery of withanolide biosynthetic gene clusters in *W. somnifera*. **a**, Synteny plot showing the 24 pseudochromosomes of *W. somnifera*. Syntenic blocks from pseudochromosome 2 are highlighted in blue to demonstrate synteny with pseudochromosome 10. **b**, Microsynteny plot of withanolide biosynthetic gene clusters. **c**, Heatmap displaying normalized log2-transformed RNA sequencing read counts for the biosynthetic cluster genes across four tissue types (old leaf, young leaf, root, and berry), n=3 biological replicates. Reads are centered around zero and values above 5 or below -5 are displayed as solid red or blue respectively. **d**, Withanolide biosynthetic gene clusters with genes colored by enzyme family (2OGD = 2-oxoglutarate-dependent dioxygenase, 24ISO = Δ^24^-isomerase, SULF = sulfotransferase, SDH = short-chain dehydrogenase, ACT = acyltransferase) or colored by specific enzyme for the P450s (CYP87G1, CYP749B2, CYP88C10, CYP88C7, and CYP88C8). Genes marked with an asterisk are pseudogenes. The green or brown dotted box indicates whether the genes belong to a root of leaf co- expressed subcluster. Distances between genes in kilobases (kb) are indicated. **e**, Correlation matrices of pairwise Pearson correlation coefficients calculated for every pair of genes within each cluster.

### Leveraging yeast metabolic engineering for candidate gene testing

Withanolides are de novo synthesized in both roots and leaves of *W. somnifera*^22^, indicating that the core biosynthetic genes are expressed in both tissues. Therefore, we limited our candidate list to genes with paralogs in both leaf and root subclusters. We further restricted the list to genes that were less than 75% identical on an amino acid level to avoid functional redundancy. Although only present in the root subcluster, *CYP88C8* was included due to its high expression level and membership in the CYP88 P450 family, members of which can hydroxylate triterpenes^23,24^. This resulted in a candidate list of seven genes in addition to *24ISO*: *CYP87G1*, *CYP88C7*, *CYP88C8*, *CYP749B2*, *CYP88C10*, *SDH2*, and *SULF1* (Supplementary Table 2).

Initially we screened candidate genes via *Agrobacterium*-mediated transient expression in *Nicotiana benthamiana* (tobacco), however, we were unable to detect new products upon expression of the candidate genes. We suspect contributing factors were low abundance of **1** in tobacco and poor ionization of sterols in our liquid chromatography-mass spectrometry (LC-MS) system. Another group independently reported that low **1** levels limited accumulation of **2** in tobacco, leading them to co-express phytosterol pathway genes along with *24ISO*^25^. Therefore, we shifted to screening candidate genes via heterologous expression in yeast.

We engineered a yeast base strain (Y7126) to produce **2** by integrating 7-dehydrocholesterol reductase from *Solanum tuberosum* (*7RED*) and *24ISO* from *W. somnifera* into a strain with deletions of ergosterol pathway genes *ERG4* and *ERG5* (Y4108) (Supplementary Table 3). *ERG4*/*ERG5* deletion causes an accumulation of ergosta-5,7,24(28)-trienol, which is reduced by 7RED to **1**, and then converted by 24ISO to **2** (Extended Data Fig. 1). Production of **1** and **2** in Y7126 was confirmed by gas chromatography mass spectrometry (GC-MS) (Extended Data Fig. 1). A cytochrome P450 reductase from *Arabidopsis thaliana* (*ATR1*) was integrated into every strain expressing a plant P450. Candidate genes were integrated into Y7126 and these strains were cultured, extracted, and tested for new products using LC-MS.

### Elucidation of lactone ring formation in withanolide biosynthesis

From the candidate enzymes tested in Y7126, only expression of CYP87G1 resulted in new products (Fig. 3, Extended Data Fig. 2). CYP87G1 produced **3**, with a mass corresponding to hydroxylated **2** and three products with masses corresponding to double hydroxylated **2**. Based on UV-vis absorbance, **3** is the major product of CYP87G1 and the double hydroxylated products likely represent overoxidation products (Extended Data Fig. 2). We purified **3** from large-scale yeast culture and elucidated the structure by nuclear magnetic resonance spectroscopy (NMR). The structure of **3** was confirmed to be (22*R*)-hydroxylated **2** (Supplementary Fig. 6-11, Supplementary Tables 4-5). We investigated the in vivo role of CYP87G1 by performing viral-induced silencing (VIGS) in *W. somnifera* plants. *CYP87G1*-silenced plants accumulated **1** and **2**, indicating that CYP87G1 acts downstream of these compounds in *W. somnifera*, consistent with the hypothesis that CYP87G1 is the second enzyme in the withanolide biosynthetic pathway after 24ISO (Supplementary Fig. 12).

**Fig. 3.**
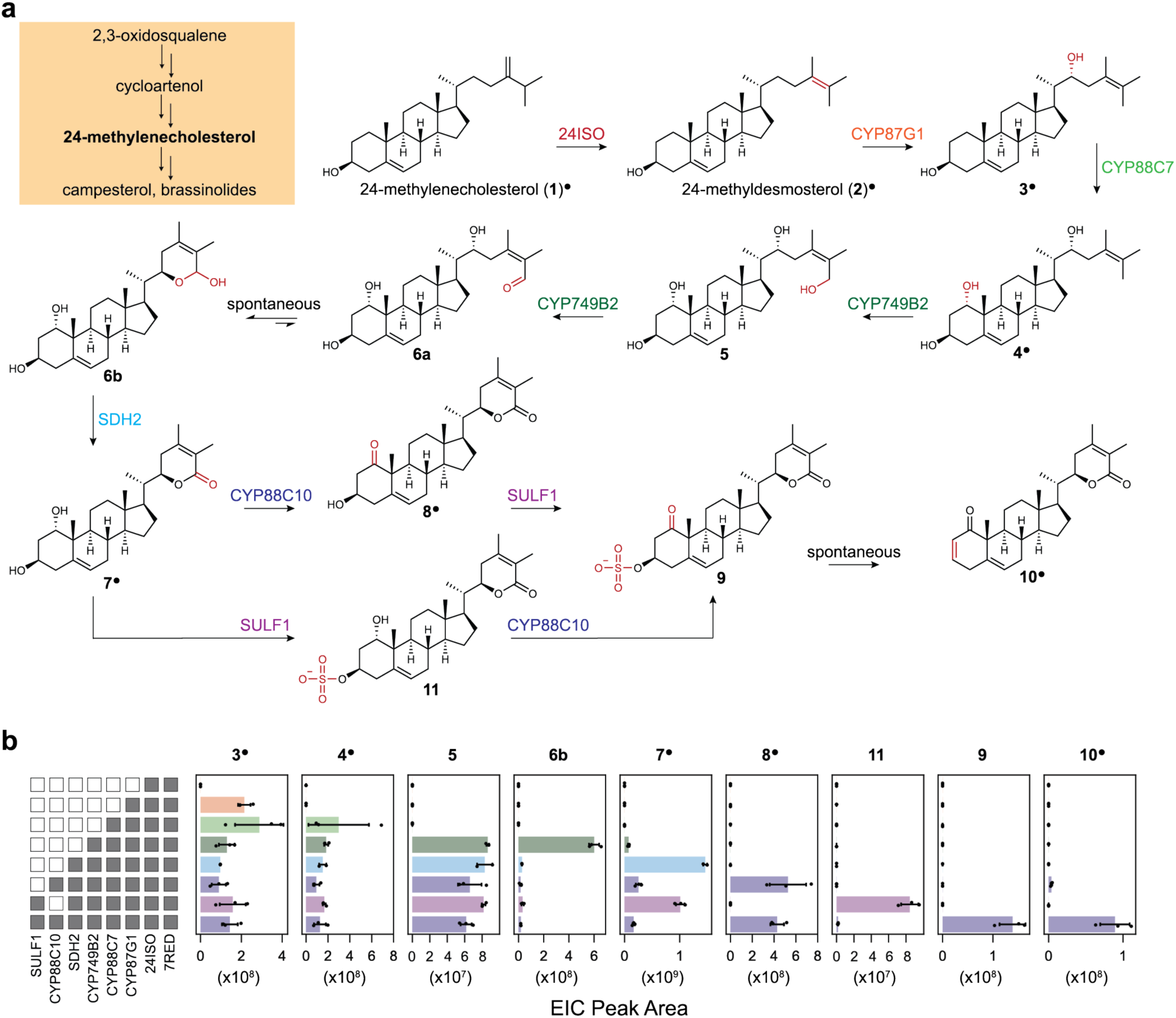
Discovery of the core withanolide biosynthesis pathway. **a,** Schematic showing steps from the pathway precursor 24-methylenecholesterol (**1**) to withanolide intermediate **10**. Superscript circles indicate that the compound structure has been confirmed by comparison with a standard or NMR analysis. **b**, Integrated extracted ion chromatogram (EIC) peak area of compounds produced by yeast strains expressing *7RED*, *24ISO*, and the six newly identified withanolide biosynthetic enzymes. The enzymes expressed in each strain are indicated by the grey boxes on the left-hand panel. Mean values are plotted of n=3 biological replicates and error bars indicate the standard deviation. Quantified ion is [M-H_2_0+H]^+^ for compounds **3**, **4**, **5**, and **6b**; [M+H]^+^ for compounds **7**, **8**, and **10**; [M-H]^-^ for compounds **9** and **11**.

We tested the remaining candidate enzymes by integration into the **3**-producing yeast strain (Y7037) and expression of two enzymes, CYP88C7 and CYP88C8, resulted in new products (Fig. 3, Extended Data Fig. 3). The major product of CYP88C7 (**4**) and the major product of CYP88C8 (**12**) have masses corresponding to hydroxylated **3**, suggesting that **3** is the substrate for both enzymes. Initially, we tested candidate enzymes in strains with both CYP88C7 and CYP88C8, but we discovered that the inclusion of CYP88C8 prevented accumulation of downstream withanolide intermediates (Extended Data Fig. 4). We suspect this occurs due to promiscuous oxidation of on-pathway intermediates by CYP88C8. We elucidated **4** and found that the additional hydroxyl group was installed at position C_1_ (Supplementary Fig. 13-18, Supplementary Tables 4-5). We silenced *CYP88C7* and *CYP88C8* in *W. somnifera* using VIGS and found this led to increased **1** when CYP88C7 was silenced and increased **1** and **2** when CYP88C8 was silenced, indicating these enzymes act downstream of withanolide pathway precursors in *W. somnifera* (Supplementary Fig. 12).

We tested the remaining candidate enzymes by integration into the **4**-producing yeast strain (Y7106) and the expression of CYP749B2 led to new products (Fig. 3, Extended Data Fig. 5). The major product of CYP749B2 (**5**) has a mass corresponding to hydroxylated **4**. CYP749B2 produced two additional products: **6b** and **7**. We purified **7** and confirmed the structure as the aglycone of withanoside V, which contains the lactone ring moiety that is characteristic of withanolides (Supplementary Fig. 19-24, Supplementary Tables 4-5). **5** is predicted to contain a hydroxyl group on C_26_, as this position must be oxygenated to enable lactonization. **6b** has a mass corresponding to the loss of two hydrogens from **5** and most likely results from a second sequential oxidation of C_26_ by CYP749B2, producing an unstable aldehyde intermediate (**6a**, not observed) which rapidly converts to **6b** by nucleophilic attack of the C_22_ alcohol at C_26_ to form a hemiacetal ring. It is unclear whether **7** is produced by a third sequential oxidation of CYP749B2 at C_26_ or by off-target activity of native yeast dehydrogenases on **6b**.

### Optimization of withanoside V aglycone (7) production in yeast

When SDH2 is expressed in combination with CYP87G1, CYP88C7, and CYP749B2, **6b** disappears and there is an approximately 10-fold increase in **7**. (Fig. 3, Extended Data Fig. 5). To determine the substrate of SDH2, we performed in vitro assays with recombinant SDH2 and extract containing both **5** and **6b** as the substrate. We observed consumption of **6b** and an approximately 10-fold increase in **7**, while the level of **5** stayed constant (Fig. 4a, Extended Data Fig. 6). Thus, SDH2 promotes lactone ring formation by oxidizing the hemiacetal ring of **6b** into the lactone ring of **7** rather than acting upon **5**. We performed docking analysis using a computationally generated SDH2 structure, which positioned the lactone ring of **7** in a catalytically sensible orientation relative to the nicotinamide moiety of the cofactor NADH, as well as the conserved catalytic residues serine and tyrosine^26^ (Supplementary Fig. 25).

**Fig. 4.**
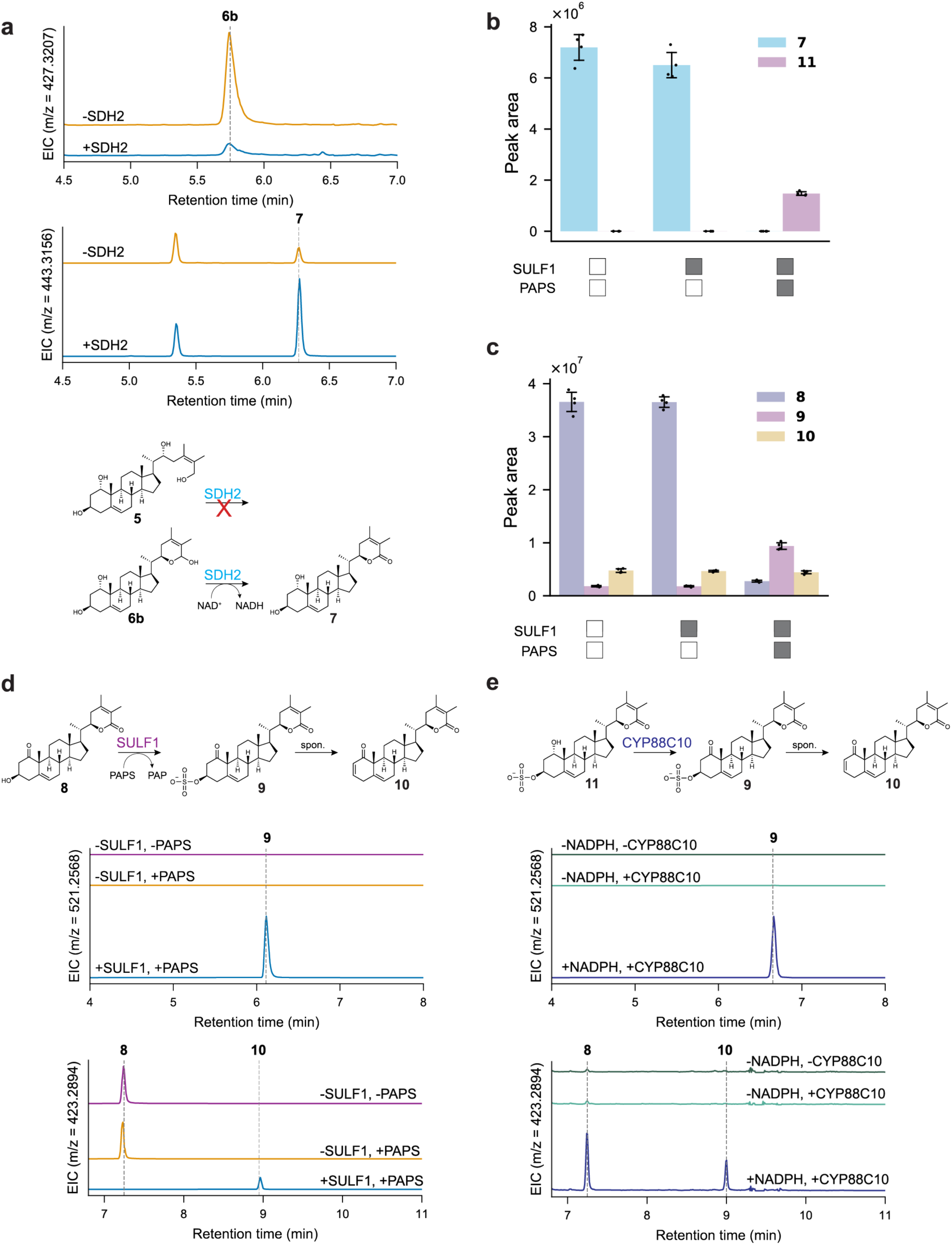
In vitro enzyme assays with SDH2, SULF1, and CYP88C10. **a,** In vitro enzyme assay with recombinant SDH2, supplied with NAD^+^ as the cofactor and extract from Y7567 as the substrate. Representative traces are shown out of n=4 replicates. **b**, In vitro enzyme assays with recombinant SULF1, supplied with 3’-phosphoadenosine-5’-phosphosulfate (PAPS) as the cofactor and extract from Y7752 as the substrate. Bar graphs display the mean value of n=4 replicates and the error bars indicate the standard deviation. Peak areas for compounds **7** and **11** are shown. **c**, In vitro enzyme assays with recombinant SULF1 as described in **b**, peak areas for compounds **8**, **9**, and **10** are shown. **d**, In vitro enzyme assays with recombinant SULF1 supplied with PAPS as the cofactor and purified **8** as the substrate. EICs shown for **9** ([M+H]^+^ ion; *m/z* = 521.2568) and **10** ([M+H]^+^ ion; *m/z* = 423.2894). Representative traces are shown of n=3 replicates. **e**, Assays with CYP88C10 microsomes supplied with NADPH as the cofactor and a mixture of **7** and **11** as the substrate. EICs shown for **9** ([M+H]^+^ ion; *m/z* = 521.2568) and **10** ([M+H]^+^ ion; *m/z* = 423.2894). Representative traces are shown of n=3 replicates.

When scaling up production of **7** for compound purification, we discovered that CYP749B2 requires higher oxygen conditions compared to CYP87G1 and CYP88C7. We successfully purified **3** (product of CYP87G1) and **4** (product of CYP88C7) from 1 L flask cultures, but production of **7** (product of CYP749B2 and SDH2) was insufficient for compound purification from these culture conditions. Concentrations of **3** and **4** were comparable between 3 mL and 1 L cultures, but concentration of **7** fell to negligible levels at the 1 L scale. To further investigate this phenomenon, we grew **4**- and **7**-producing strains (Y7106 and Y7610) with shaking at 250 rpm and 350 rpm in 3 mL cultures. We found that while **3** and **4** production was higher when cultures were grown at 250 rpm for both strains, **7** production was approximately 10-fold higher when cultures were grown at 350 rpm (Extended Data Fig. 7). This led us to employ a bioreactor with controllable dissolved oxygen concentration to produce **7**, a strategy that greatly improved production and enabled structural elucidation.

### Elucidation of C1 ketone formation and C2-C3 desaturation

Expression of the remaining candidate enzymes in a **7**-producing yeast strain (Y7610) revealed a sulfotransferase (SULF1) and a fourth P450 (CYP88C10) involved in withanolide biosynthesis. Earlier steps are strictly linear but **7** represents a branchpoint in the pathway as both CYP88C10 and SULF1 can consume **7**. Expression of CYP88C10 led to the appearance of a new product (**8**) with a mass corresponding to the loss of two hydrogens from **7** (Fig. 3, Extended Data Fig. 8). The structure of **8** was elucidated (Supplementary Fig. 26-32, Supplementary Tables 4-5) and differed from **7** by the oxidation of the C_1_ hydroxyl to a ketone. The C_1_ ketone is a widespread chemical feature among the withanolides and thus CYP88C10 represents a core withanolide biosynthetic enzyme. Expression of SULF1 resulted in the detection of a new product (**11**) with a mass corresponding to sulfated-**7** (Fig. 3, Extended Data Fig. 8). The MS/MS spectrum of **11** matches that of **7**, corroborating the structure of **11** as sulfated-**7** (Extended Data Fig. 8). We silenced *SULF1* in *W. somnifera* using VIGS and found this led to increased **1**, providing evidence that this enzyme is involved in withanolide biosynthesis. (Supplementary Fig. 12).

Co-expression of SULF1 and CYP88C10 in Y7610 resulted in a decrease in **11** and the appearance of two new products: **9** with a mass corresponding to sulfated-**8**/oxidized-**11** and **10** with a mass corresponding to dehydrated **8** (Fig. 3, Extended Data Fig. 8). We did not structurally elucidate **11** or **9** but two lines of evidence support the proposal that C_3_ is sulfated: a) sulfated withanolides detected in *W. somnifera* are always sulfated at the C_3_ position^27,28^ and b) the only free hydroxyl available for sulfation on **8** is at C_3_. We elucidated the structure of **10** via NMR as an intermediate containing all characteristic chemical features of withanolides: lactone ring, C_1_ ketone, and C_2_-C_3_ unsaturated bond on the A-ring (Supplementary Fig. 33-37, Supplementary Tables 4-5). To cross-validate our results in an orthogonal system, we reconstituted the six identified withanolide biosynthetic enzymes by transient expression tobacco (Extended Data Fig. 9). Early withanolide pathway intermediates were undetectable but compounds **6b**-**10** detected, cross-validating the results from pathway reconstitution in yeast. We detected **7**, **9**, and **10** in methanolic extracts of young leaf *W. somnifera* tissue, providing additional support for the biological relevance of these intermediates and the enzymes that produce them (Extended Data Fig. 10).

We performed in vitro enzyme assays to determine the order of SULF1 and CYP88C10 in the pathway to **10**. Incubation of SULF1 with extract from a yeast strain expressing all pathway enzymes (Y7752) led to consumption of **7** and **8** with concurrent production of sulfated-**7** (**11**) and sulfated-**8** (**9**) (Fig. 4b,c), while the level of **10** was unchanged. However, we wondered if the baseline **10** in Y7752 extract could be obscuring **10** generated during the assay. We performed additional SULF1 enzyme assays with purified **8** as the substrate and found that both **9** and **10** were produced (Fig. 4d). Additionally, we performed assays with microsomes isolated from a yeast strain expressing *CYP88C10* (Y7215) and used either purified **7** or a mixture of **7** and **11** as the substrate (Fig. 4e). **8** was produced in both cases, but compounds **9** and **10** were only produced when **11** was present in the substrate mixture. These results indicate there are two possible routes to **9** (Fig 3a) and it is not clear that one is preferred over the other. The production of **10** in both CYP88C10 and SULF1 assays suggests that **10** can form spontaneously from **9**.

To address the question of whether **10** can form spontaneously from **9**, we sought to purify **9**. We converted **8** to a mixture of **9** and **10** in a large-scale SULF1 reaction. We then separated **9** from **10** using preparative high-performance liquid chromatography (HPLC) and used this semi-pure **9** in further assays. Incubation of **9** in potassium phosphate buffer (pH 7.5) overnight at 30 °C led to spontaneous production of **10** (Supplementary Fig. 38).

## Discussion

In this study, we produced a chromosome-scale genome assembly for the important medicinal plant *W. somnifera*. Using this resource, we identified and partially characterized two biosynthetic gene clusters related to withanolide biosynthesis. The two biosynthetic gene clusters we identified are among the largest and most complex identified in plants, and additionally they exhibit an unusual tissue-specific subcluster structure. Although co-expression among genes in a biosynthetic gene cluster is common, there are only a few reported cases of tissue-specific expression differing within a cluster and the biological reason for this organization is unclear^29,30^. We explored the possibility that root- and leaf-expressed paralogs of the same enzyme might be responsible for observed differences in withanolide profile between root and leaf, but did not find any divergent activity between paralogs. More plausibly, enzymes specific to the leaf subcluster (*ACT*) or to the root subcluster (*CYP88C8* and *OGDs*) may be responsible for tissue-specific derivatization of the core withanolide scaffold.

Further work is necessary to produce withanolides that have proven therapeutic potential, such as withaferin A and withanolide A. The remaining uncharacterized genes across the clusters include *CYP88C8*, the *2OGDs*, and *ACT*. While we demonstrated that CYP88C8 can oxidize withanolide intermediates, we have not fully characterized this enzyme. The observation that CYP88C8 suppresses the formation of on-pathway withanolide intermediates when co-expressed with other biosynthetic enzymes is puzzling. It is possible that the catalysis of CYP88C8 is controlled in *W. somnifera* through a mechanism such as metabolon formation or sub-compartmentalization of the withanolide biosynthesis pathway across different cell types that is not reproduced in our heterologous expression system. Root-expressed enzymes CYP88C8 and the 2OGDs may represent the remaining steps in the biosynthesis to root-produced withanolides such as withanolide A. However, it is likely that non-clustered enzymes are involved in the biosynthesis of leaf-produced withanolides such as withaferin A. These non-clustered enzymes could be identified in future studies from co-expression analyses using the enzymes reported here as bait.

In summary, we discovered a short-chain dehydrogenase (SDH2), a sulfotransferase (SULF1), and four cytochrome P450s (CYP87G1, CYP88C7, CYP749B2, CYP88C10) involved in withanolide biosynthesis. SDH2 is important for lactone ring formation: expression increases production of **7** by approximately 10-fold, making SDH2 crucial for engineering withanolide production in yeast. Our discovery aligns with previous reports of short-chain dehydrogenases involved in lactone ring formation in momilactone and noscapine biosynthesis^31–33^,. SULF1 is capable of sulfating **8** to produce **9**, which can then spontaneously convert to **10**. The presence of sulfotransferases in the biosynthetic gene cluster was initially confusing because sulfated withanolides are rare^10^ and while the role of sulfated natural products in plants is largely unknown^34^, sulfation is considered a tailoring step in vertebrate sterol biosynthesis where it aids in transport and excretion of sterols by increasing solubility^35^. The discovery of SULF1 as a core pathway enzyme in withanolide biosynthesis challenges the conventional view of sulfation as a tailoring reaction and hints at a larger role for sulfotransferases in plant secondary metabolism.

Co-expression of the six withanolide biosynthetic enzymes discovered here results in production of **10** containing all defining chemical features of withanolides: the lactone ring, C_1_ ketone, and C_2_-C_3_ A-ring unsaturation. The structure of **10** differs from the “commonly diversifiable withanolide intermediate” in the published synthetic route to withanolides by only one hydroxylation^14^. In the process of enzyme characterization, we built yeast strains capable of producing milligram-per-liter levels of withanolide pathway intermediates including **10**, representing an important starting point for the scale-up of withanolide production and the development of withanolide-derived pharmaceuticals.

## Extended Data Figures

**Extended Data Fig. 1.**
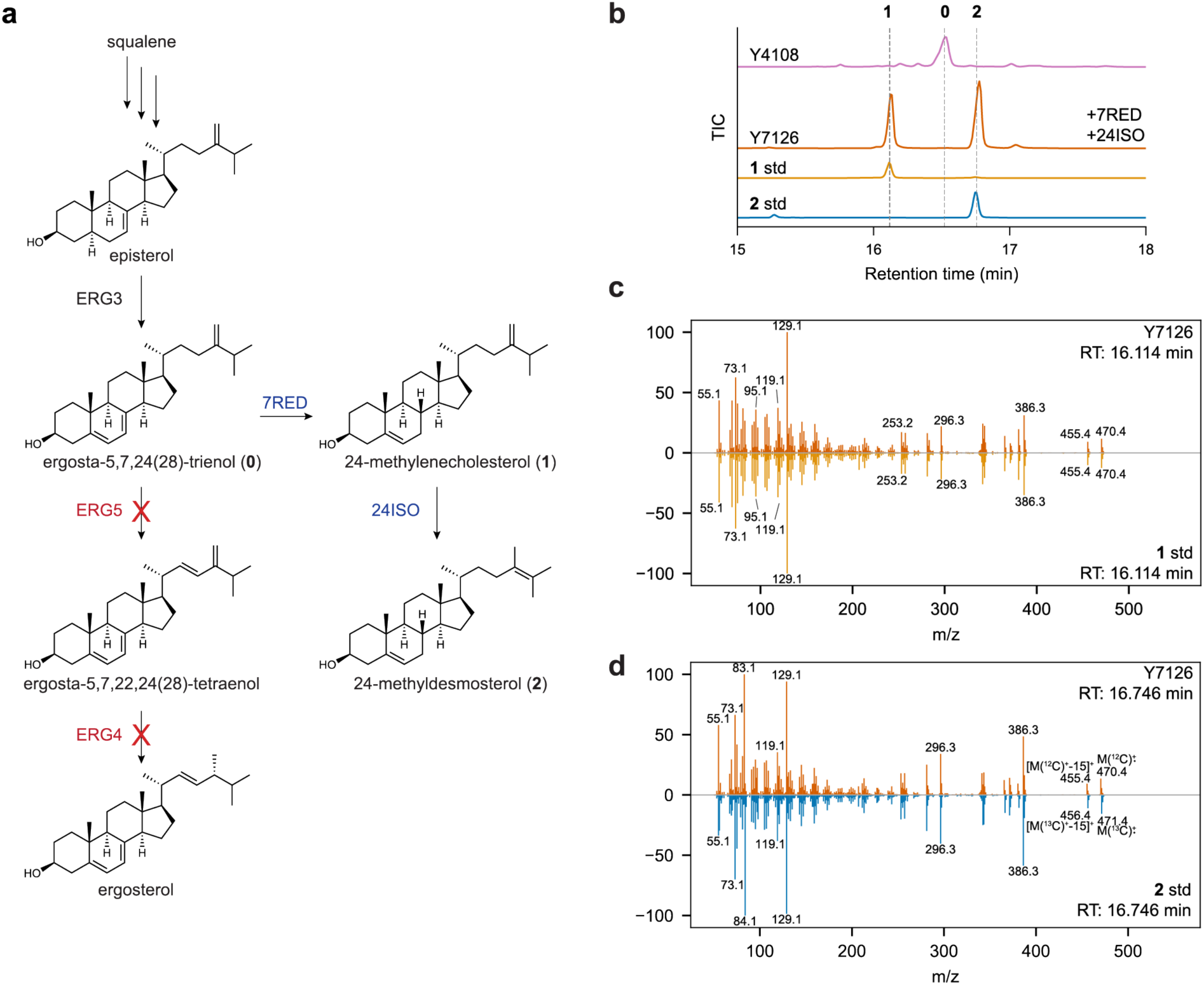
Metabolic engineering of 2-producing yeast strain Y7126. **a,** Schematic of metabolic engineering to create **2**-producing yeast strain Y7126. **b**, Confirmation of **1** and **2** production by Y7126. Representative gas chromatography-mass spectrometry (GC-MS) total ion chromatograms (TICs) are shown of n=3 biological replicates. Products of *ERG4*/*ERG5* deletion strain (Y4108) and **2**-producing base strain (Y7126) are indicated, along with an authentic standard for **1** and a [26-^13^C]-labeled standard for **2**. **0** is the ergosterol pathway intermediate ergosta-5,7,24(28)-trienol. **c**, MS/MS spectra of Y7126 product and authentic **1** standard. **d**, MS/MS spectra of Y7126 product and [26-^13^C]-labeled **2** standard.

**Extended Data Fig. 2.**
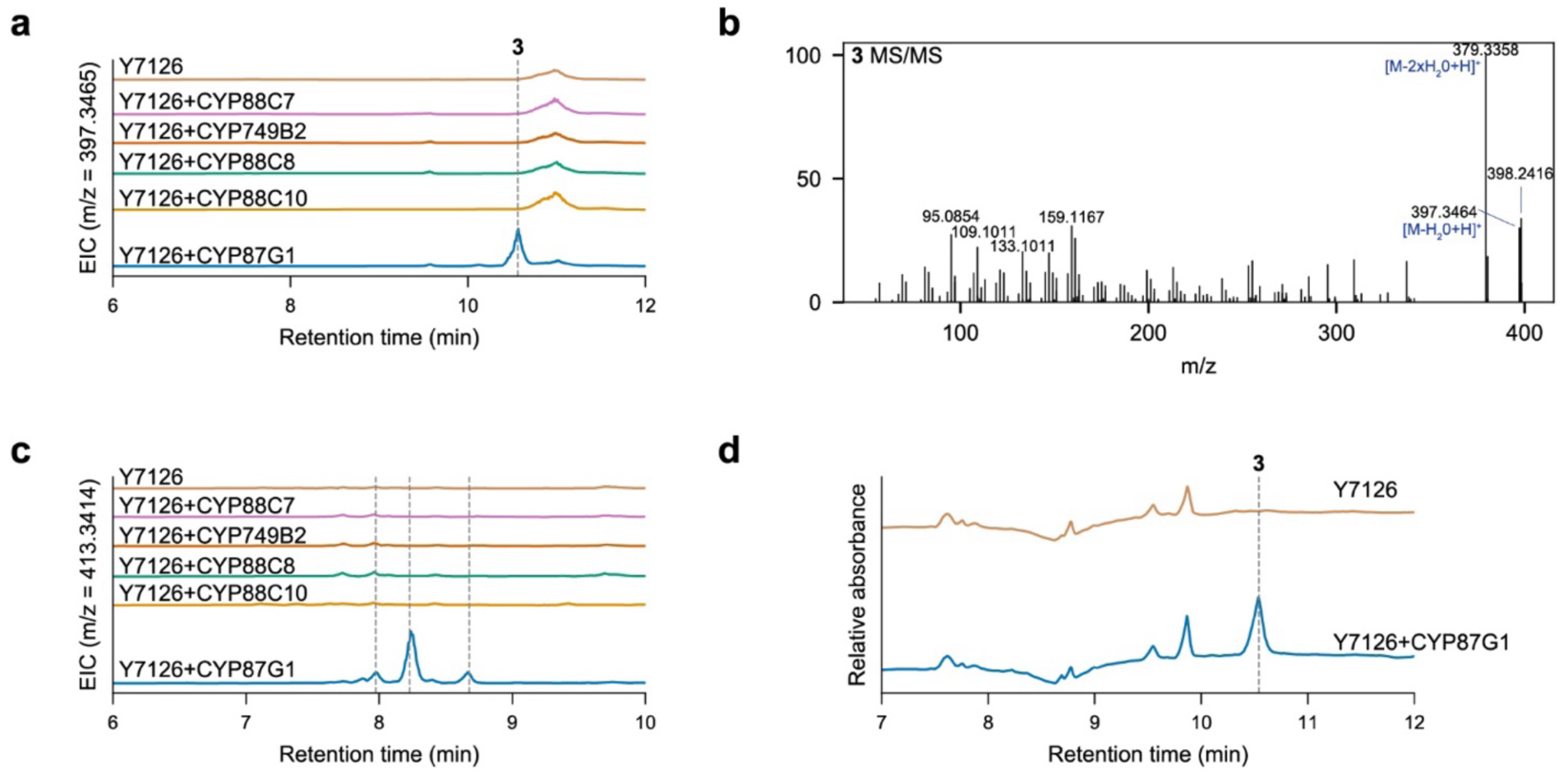
Testing candidate genes in yeast strain Y7126. **a,** Candidate cytochrome P450s were expressed in **2**-producing yeast strain Y7126. EIC for [M-H_2_0+H]^+^ ion corresponding to hydroxylated **2** (C_28_H_46_O_2_; *m/z* = 397.3465). **b**, MS/MS spectrum from **3** produced by Y7037 (Y7126+CYP87G1). Precursor ion was *m/z* = 397.3465 at retention time 10.59 mins. **c**, EIC for [M-H_2_0+H]^+^ ion corresponding to double hydroxylated **2** (C_28_H_46_O_3_; *m/z* = 413.3414). **d**, Relative absorbance (UV-vis 190-800 nm) as measured by the photo diode array (PDA) detector on the same LC method as in **a** and **c**.

**Extended Data Fig. 3.**
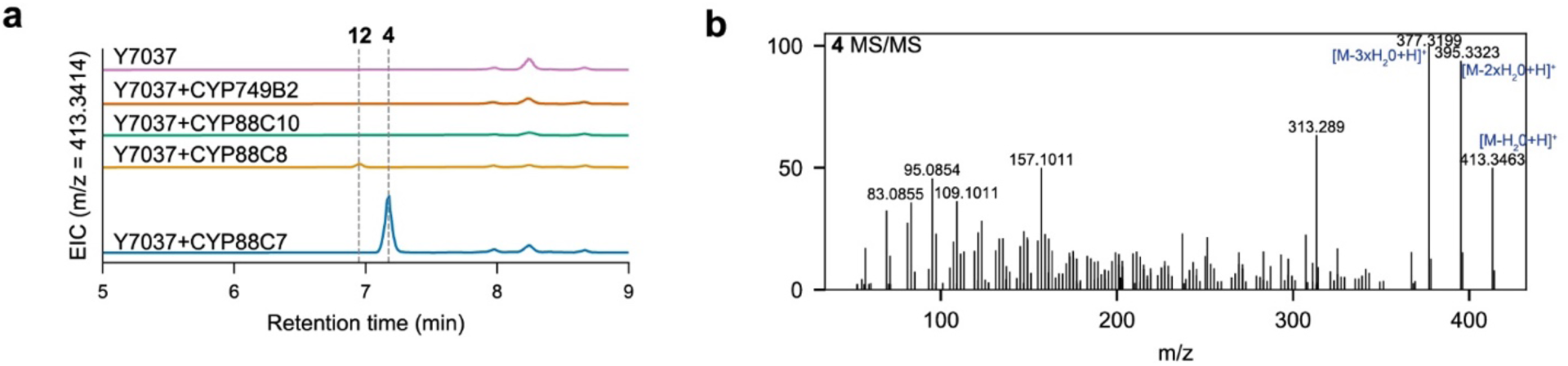
Testing candidate genes in yeast strain Y7037. **a**, Candidate cytochrome P450s were expressed in **3**-producing yeast strain Y7037. EIC for [M-H_2_0+H]^+^ ion corresponding to hydroxylated **3** (C_28_H_46_O_3_; *m/z* = 413.3414). **b**, MS/MS spectrum for **4** produced by Y7106 (Y7037+CYP88C7). Precursor ion was *m/z* = 413.3412 at retention time 7.19 mins.

**Extended Data Fig. 4.**
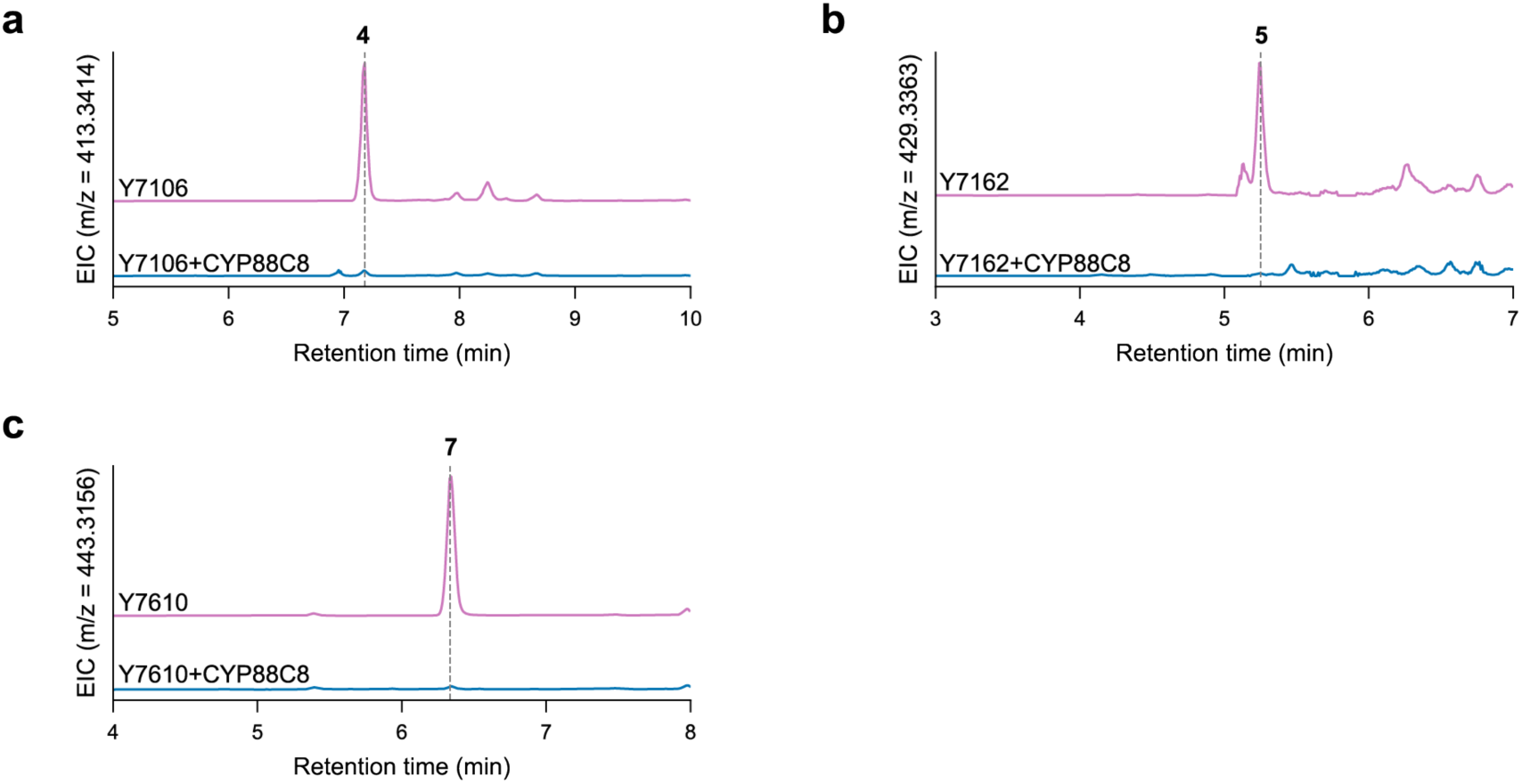
Expression of CYP88C8 suppresses formation of on-pathway intermediates. **a,** When CYP88C8 is expressed in a **4**-producing yeast strain, **4** accumulation is diminished. EIC for [M- H_2_0+H]^+^ ion corresponding to **4** (C_28_H_46_O_3_; *m/z* = 413.3414). **b**, When CYP88C8 is expressed in a **5**- producing yeast strain, **5** no longer accumulates. EIC for [M-H_2_0+H]^+^ ion corresponding to **5** (C_28_H_46_O_4_; *m/z* = 429.3363). **c**, When CYP88C8 is expressed in a **7**-producing yeast strain, **7** accumulation is diminished. EIC for [M+H]^+^ ion corresponding to **7** (C_28_H_42_O_4_; m/z = 443.3156).

**Extended Data Fig. 5.**
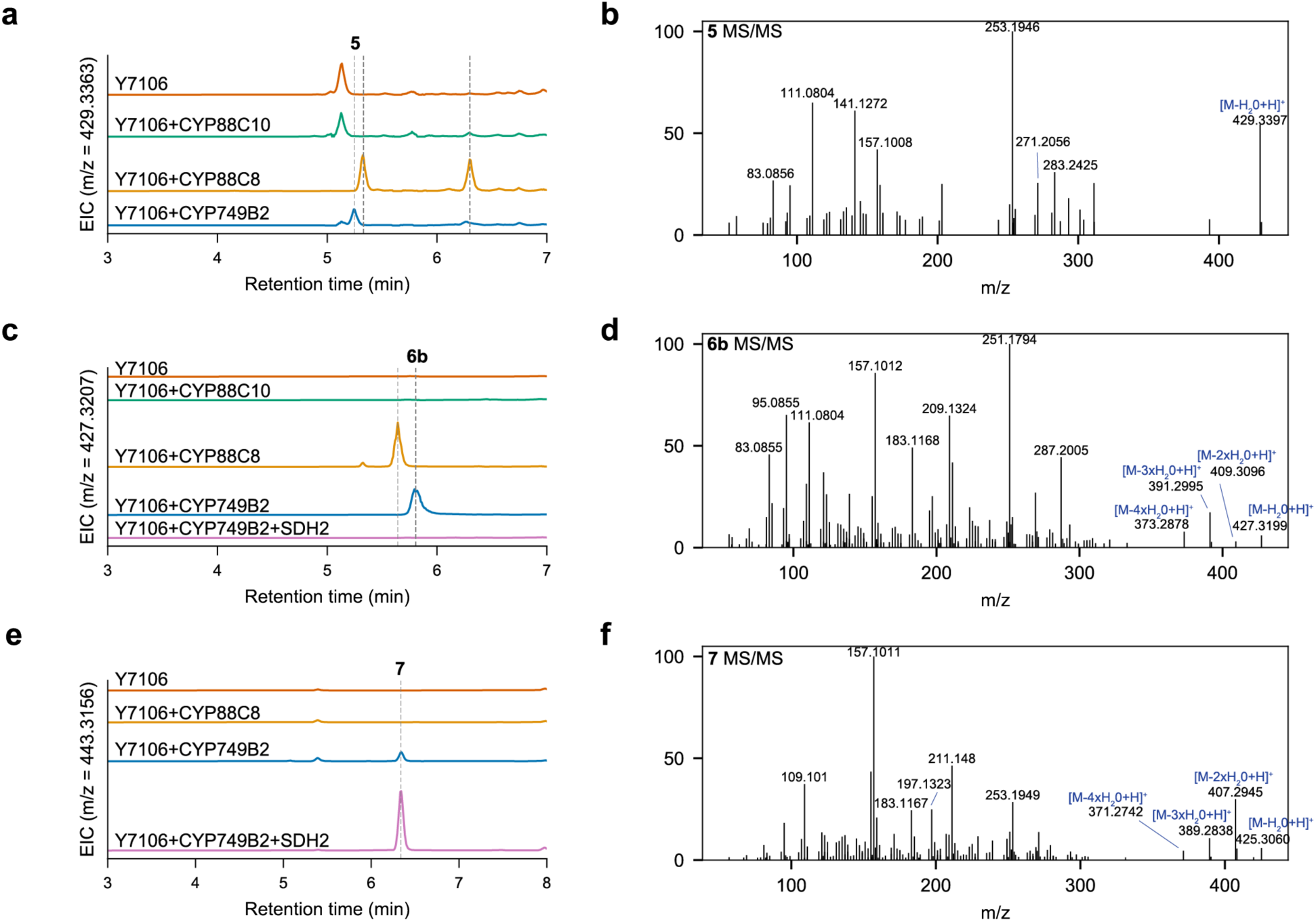
Testing candidate genes in yeast strain Y7106. **a,** Candidate cytochrome P450s were expressed in **4**-producing yeast strain Y7106. EIC for [M-H_2_0+H]^+^ ion corresponding to hydroxylated **4** (C_28_H_46_O_4_; m/z = 429.3363). **b**, MS/MS spectrum for **5** produced by Y7162 (Y7106+CYP749B2). Precursor ion was *m/z* = 429.3361 at retention time 5.26 mins. **c**, EIC for [M-H_2_0+H]^+^ ion corresponding to the formula C_28_H_44_O_4_ (*m/z* = 427.3207). **d**, MS/MS spectrum for **6b** produced by Y7162 (Y7106+CYP749B2). Precursor ion was *m/z* = 427.3205 at retention time 5.8 mins. **e**, EIC for [M+H]^+^ ion corresponding to the formula C_28_H_42_O_4_ (*m/z* = 443.3156). **f**, MS/MS spectrum for **7** produced by Y7610 (Y106+CYP749B2+SDH2). Precursor ion was *m/z* = 443.3152) at retention time 6.35 mins.

**Extended Data Fig. 6.**
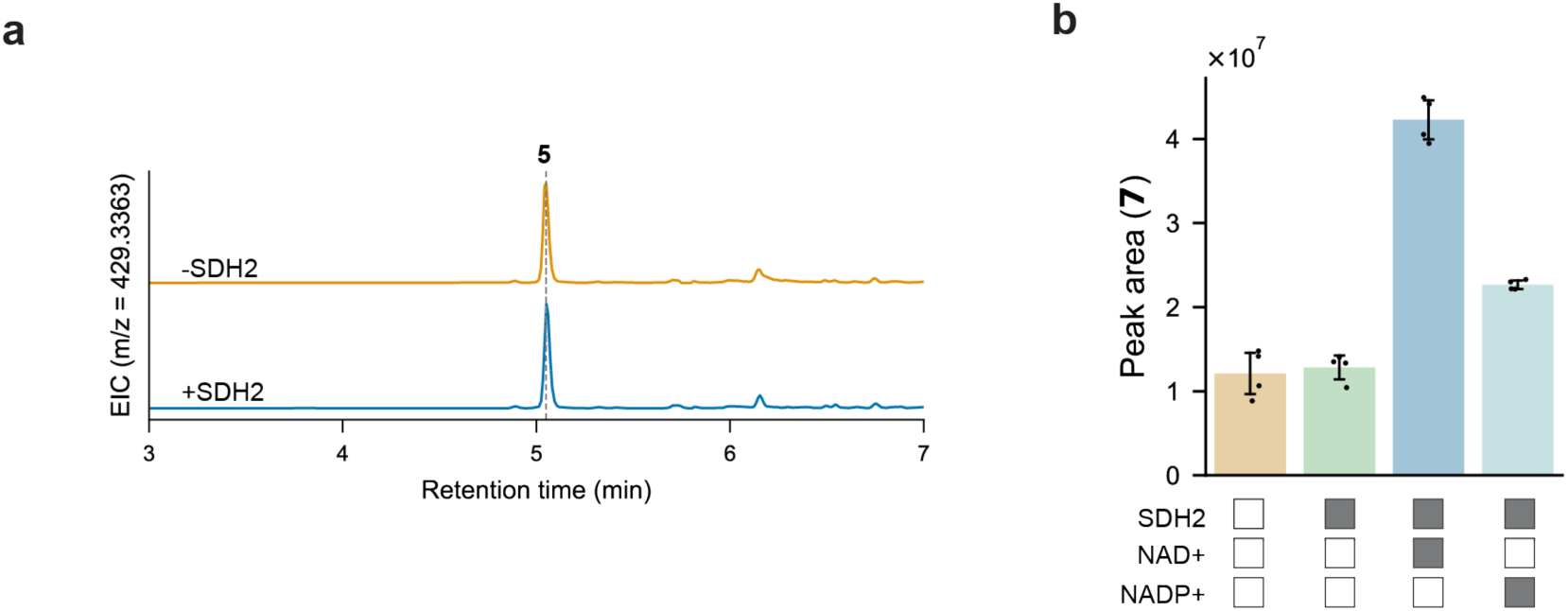
SDH2 in vitro enzyme assays. **a,** In vitro enzyme assays with recombinant SDH2, supplied with NAD^+^ as the cofactor and extract from Y7567 as the substrate. EIC for [M-H_2_0+H]^+^ ion for **5** (*m/z* = 429.3363). Representative traces are shown of n=4 replicates. **b**, Peak area of **7** for SDH2 in vitro enzyme assays with either NAD^+^ or NADP^+^ supplied as the cofactor and extract from Y7567 as the substrate. Bar graphs show the mean value of n=4 replicates and error bars indicate the standard deviation.

**Extended Data Fig. 7.**
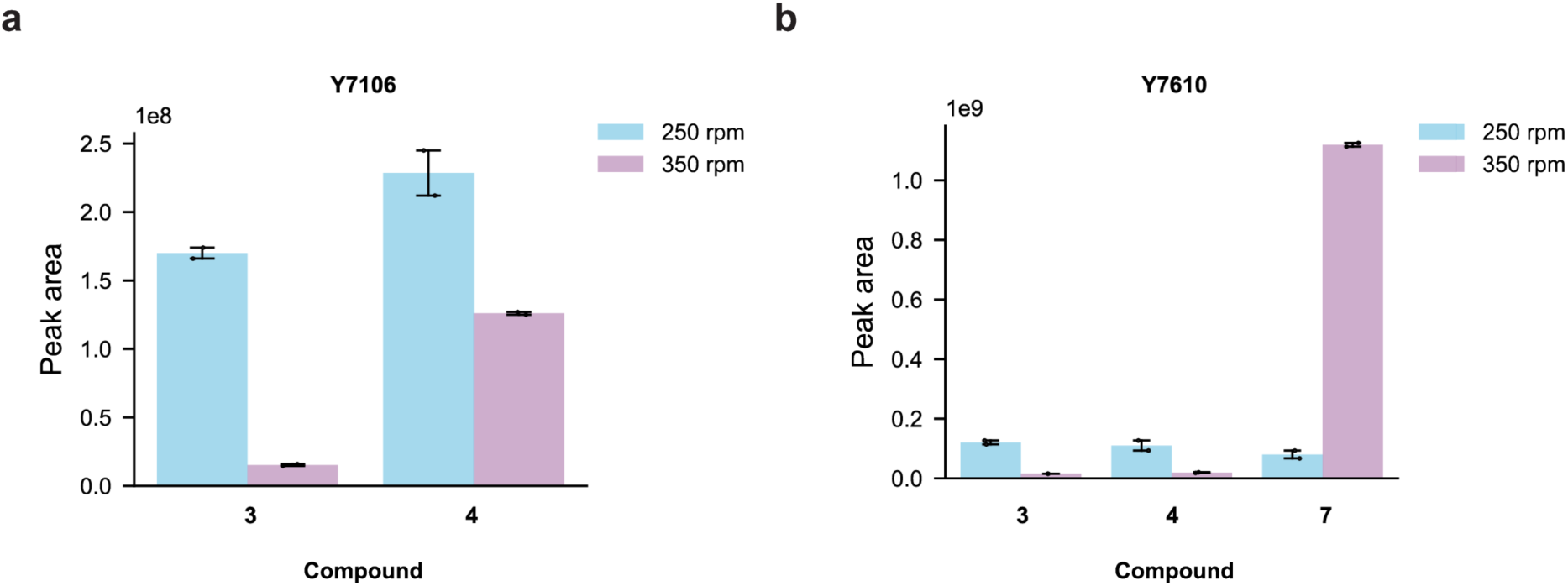
Effect of shaking speed on compound production in strains Y7106 and Y7610. **a,** Compound production by **4**-producing strain Y7106 grown with shaking at 250 or 350 rpm. Bar graphs shown the mean value of n=2 replicates and error bars indicate the standard deviation. **b**, Compound production by **7**-producing strain Y7610 grown with shaking at 250 or 350 rpm. Bar graphs shown the mean value of n=2 replicates and error bars indicate the standard deviation.

**Extended Data Fig. 8.**
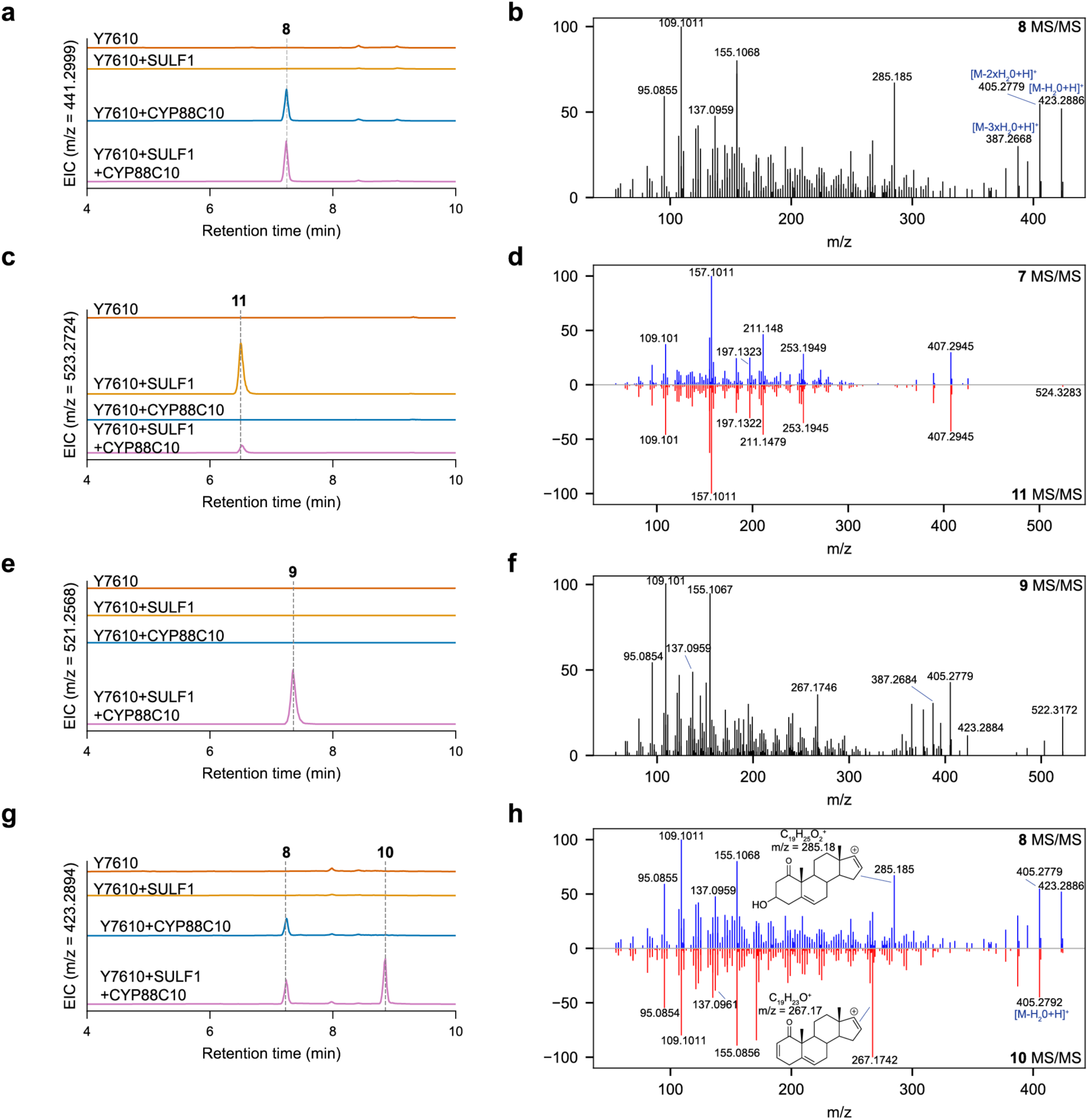
Characterization of products of CYP88C10 and SULF1. **a,** Candidate genes SULF1 and CYP88C10 were expressed in **7**-producing yeast strain Y7610. EIC for [M+H]^+^ ion corresponding to C_28_H_40_O_4_ (*m/z* = 441.2999). **b**, MS/MS spectrum for **8** produced by Y7658 (Y7610+CYP88C10). Precursor ion was *m/z* = 441.2997 at retention time 7.26 mins. **c**, Candidate genes SULF1 and CYP88C10 were expressed in **7**-producing yeast strain Y7610. EIC for [M+H]^+^ ion corresponding to C_28_H_42_O_7_S (*m/z* = 523.2724). **d**, MS/MS spectrum for **11** produced by Y7748 (Y7610+SULF1) and comparison to MS/MS spectrum for **7**. Precursor ion for **11** MS/MS was *m/z* = 523.2723 at retention time 6.54 mins. **e**, Candidate genes SULF1 and CYP88C10 were expressed in **7**- producing yeast strain Y7610. EIC for [M+H]^+^ ion corresponding to C_28_H_40_O_4_S (*m/z* = 521.2568). **f**, MS/MS spectrum for **9** produced by Y7752 (Y7610+SULF1+CYP88C10). Precursor ion was *m/z* = 521.2567 at retention time 7.35 mins. **g**, Candidate genes SULF1 and CYP88C10 were expressed in **7**- producing yeast strain Y7610. EIC for [M+H]^+^ ion corresponding to C_28_H_38_O_3_ (*m/z* = 423.2894). **h**, MS/MS spectrum for **10** produced by Y7752 (Y7610+SULF1+CYP88C10) and comparison to MS/MS spectrum for **8**. Precursor ion for **10** was *m/z* = 423.2894 at retention time 8.84 mins.

**Extended Data Fig. 9.**
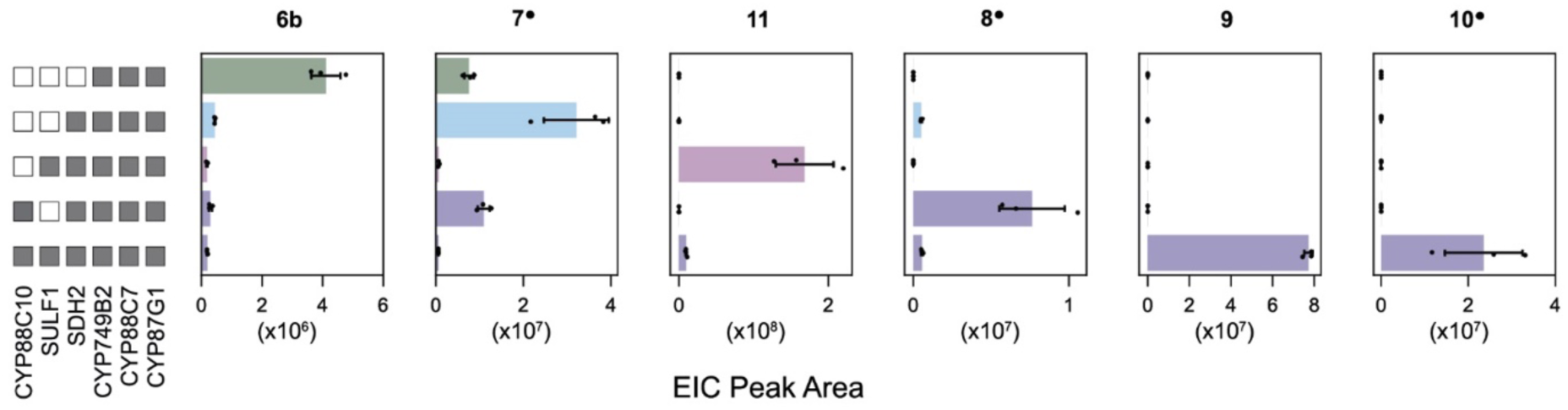
Cross-validation of the core withanolide biosynthesis pathway in tobacco. EIC peak area of compounds produced by *Agrobacterium*-mediated transient expression of withanolide biosynthetic enzymes in tobacco (compounds **6b**-**10**, see Fig. 3a for structures). Superscript circles indicate that the compound structure has been confirmed by either comparison with a standard or by NMR analysis. The enzymes co-expressed in each experiment are indicated by the grey boxes on the left-hand panel. Mean values are plotted of n=3 biological replicates and error bars indicate the standard deviation. Quantified ion is [M+H]^+^ for compounds **6b**, **7**, **8**, and **10** and [M-H]^-^ for compounds **9** and **11**.

**Extended Data Fig. 10.**
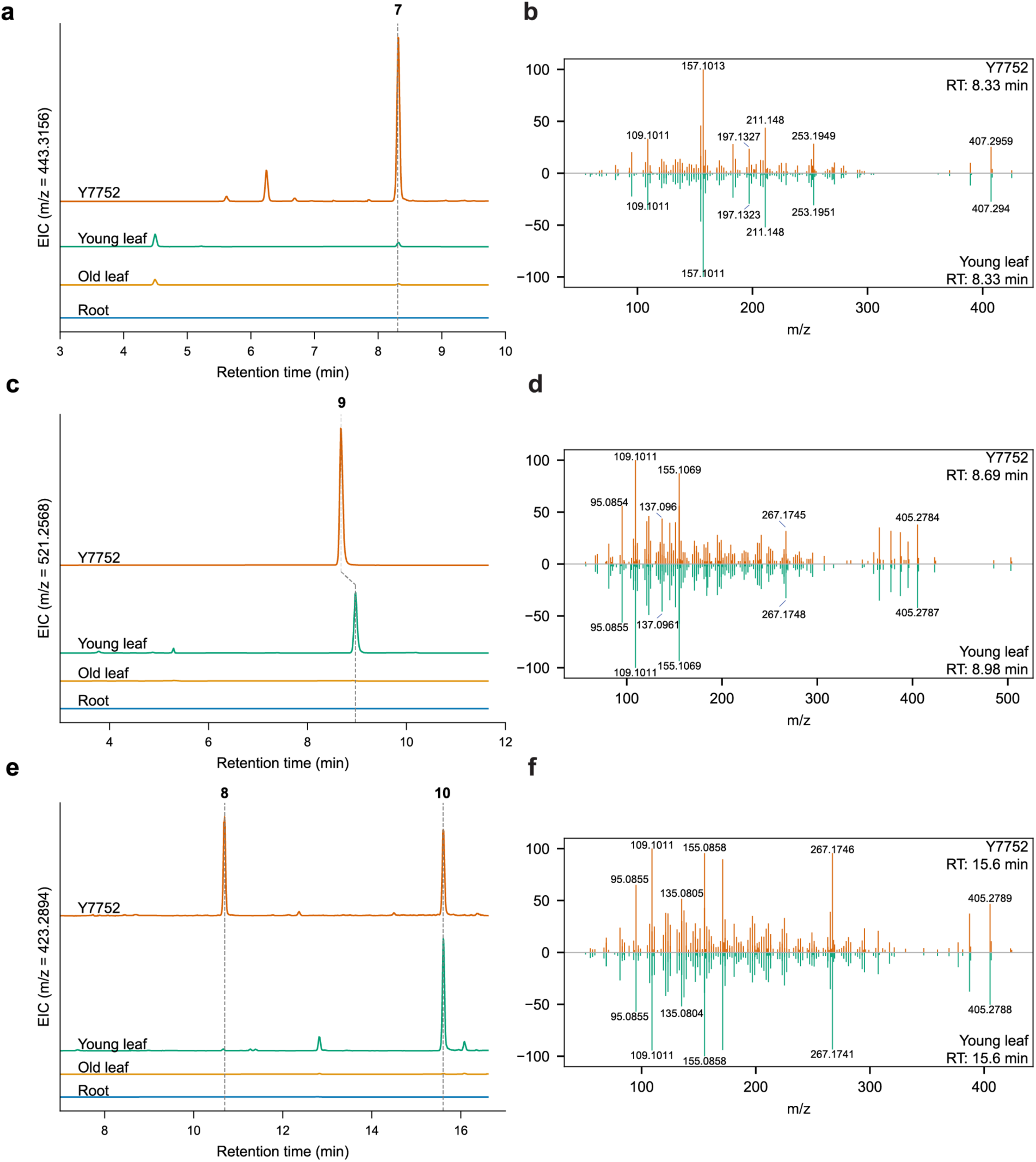
Intermediates in the withanolide biosynthesis pathway accumulate in *W. somnifera* tissue. **a**, Methanolic extracts of young leaf, old leaf, and root *W. somnifera* tissue and ethyl acetate extracts from **10**-producing yeast strain Y7752. EIC for **7** ([M+H]^+^ ion; *m/z* = 443.3156). **b**, MS/MS spectra for **7** produced by Y7552 and young leaf tissue. **c**, Methanolic extracts of young leaf, old leaf, and root *W. somnifera* tissue and ethyl acetate extracts from **10**-producing yeast strain Y7752. EIC for **9** ([M+H]^+^ ion; *m/z* = 521.2568). Sulfated compounds such as **9** exhibit retention time shifting on our LC-MS method, even within a run. **d**, MS/MS spectra for **9** produced by Y7552 and young leaf tissue. **e**, Methanolic extracts of young leaf, old leaf, and root *W. somnifera* tissue and ethyl acetate extracts from **10**-producing yeast strain Y7752. EIC for **10** ([M+H]^+^ ion; *m/z* = 423.2894). **8** is detected ([M-H_2_0+H]^+^ ion; *m/z* = 423.2894) in Y7752 extract only. **f**, MS/MS spectra for **10** produced by Y7552 and young leaf tissue.

## ACKNOWLEDGMENTS

This work was supported by the Keck Foundation and the Schooner Foundation. E.R. was supported by a National Science Foundation Graduate Research Fellowship. E.K. was supported by Japan Society for the Promotion of Science (JSPS). We thank the Genome Technology Core at Whitehead Institute for performing RNA sequencing and Prof. David Nelson (University of Tennessee) for helping with the nomenclature of cytochrome P450 enzymes. We thank Jason Matos for assisting with SDH2 purification. We also acknowledge valuable discussions and suggestions from members of the Weng Lab. The authors express gratitude for the staffing and instrumentation support provided by the Northeastern University NMR Core Facility. This work utilized an NMR spectrometer that was purchased with funding from a National Institutes of Health SIG grant (S10OD032452). The content is solely the responsibility of the authors and does not necessarily represent the official views of the National Institutes of Health.

## AUTHOR CONTRIBUTIONS

E.R. and J.K.W. led and designed the experiments. B.C. performed the initial work on the project. M.X. purified SULF1 and contributed to the initial project work, including cloning candidate P450s into the pEAQ-HT vector. F.S.L. performed NMR and elucidated compound structures. E.R., M.T., and F.S.L. purified compounds. E.K. performed VIGS experiments. E.R. conducted the assembly and annotation of the *W. somnifera* genome, synteny analysis, RNA sequencing, co-expression analysis, metabolic engineering in yeast, bioreactor fermentations, transient expression in *N. benthamiana*, purification of SDH2, in vitro enzyme assays, and LC-MS and GC-MS analysis. M.T. assisted with various experiments, including yeast engineering and metabolic profiling of *W. somnifera* tissues. T.M. supported numerous experiments, including yeast engineering and cloning SDH2 into the pHis8b vector. J.H. contributed to several experiments, including yeast engineering and generating the RNA sequencing read count heatmap for the clusters. J.K.W. performed docking analysis and computational modeling of SDH2 structure. T.M. and E.R. wrote Python scripts to process data and generate graphs. E.R., M.T., T.M., J.H., M.X., E.K., B.C., F.S.L., and J.K.W. contributed to writing the manuscript.

## DATA AVAILABILITY

The fully assembled and annotated *W. somnifera* genome sequences and RNA-seq dataset will be deposited in NCBI upon formal publication of this manuscript in a peer-reviewed journal.

## Supplemental information

### Materials and Methods

#### Analytic standards

Standards were obtained from the following sources: withaferin A (Millipore Sigma), withanolide A (Cayman Chemical), and 24-methylenecholesterol (Avanti polar lipids, inc.). [26-^13^C]-labeled 24-methyldesmosterol was prepared and structurally confirmed in a previous study^1^. Quantification of withanolide A and withaferin A for metabolic profiling of *W. somnifera* plant tissues was based on a standard curve.

#### Plant materials

*W. somnifera* seeds were acquired from “All good things organic seeds” on Amazon.com in March 2017*. N. benthamiana* seeds were a gift from the S. L. Lindquist laboratory, Whitehead Institute. For growing *W. somnifera* and *N. benthamiana*, seeds were sown on potting mix with added vermiculite and vernalized for three to five days at 4 °C under a humidified lid. They were then grown on in-door growth shelves installed with full-spectrum LED growth lights (80–120 μmol photons m^−2^ s^−1^) under a 16 h light/8 h dark photoperiod at 22 °C.

#### *W. somnifera* metabolic profiling

30 ± 2 mg fresh weight of *W. somnifera* tissue was flash frozen in liquid nitrogen in 2 mL microfuge tubes containing 2 mm zirconia/silica disruption beads. Samples were lyophilized overnight and dry weight was measured. Samples were homogenized using a Qiagen TissueLyser II at 30 Hz for 2 minutes. 1:50 dry weight/volume methanol was added to each sample, then homogenized at 30 Hz for 4 minutes. Samples were incubated at room temperature for 20 mins, then diluted 1:5 v/v in methanol. Diluted samples were centrifuged at 21130 rcf for 2 mins, then 400 μL of supernatant was removed and filtered using 0.2 µm PTFE filters. Filtrate was run on the LC-MS using the method described in LC-MS analysis.

#### Genome sequencing and assembly

Genomic DNA from *W. somnifera* was extracted from 1.5 g of leaf tissue flash frozen in liquid nitrogen and ground to powder using a mortar and pestle. Powder was processed using the NucleoBond Takara Bio kit. PacBio long-read sequencing of genomic DNA (using 4 Sequel II SMRT cells) was performed by Icahn School of Medicine at Mount Sinai. Genome assembly was performed in-house using the Improved Phased Assembly (IPA) workflow^2^ resulting in a draft genome assembly with 532 contigs, an N50 of 9 Mbp, and a total length of 2.79 Gbp. Hi-C sequencing was performed on flash frozen *W. somnifera* leaf tissue by Phase Genomics. Juicer^3^ was used to align Hi-C reads to the draft genome assembly and 3d-dna^4^ was used to generate chromosome-length scaffolds. The output from 3d-dna was viewed in Juicebox for manual curation. The 26 pseudochromosomes identified by 3d-dna were manually curated to 24 pseudochromosomes because karyotyping analysis on *W. somnifera* determined the haploid chromosome number to be n=24 in previous studies^5^. The final pseudochromosome-level assembly contained 419 scaffolds with a total length of 2.79 Gbp and an N50 of 119.6 Mb. The 24 pseudochromosomes accounted for 99.5% of the genome assembly and with only 13.7 Mb of remaining contigs. The assembly was 98.6% complete according to BUSCO^6^ analysis using the eudicots_odb10 database.

#### RNA sequencing

RNA was extracted from four different tissue types of *W. somnifera* (young leaves, old leaves, berry, and root) with three biological replicates per tissue. Young leaves were defined as leaves smaller than 2 cm and were plucked from the top of the plant, near the budding region. Young leaves, old leaves, and root tissue were harvested from ∼6-week-old plants and berry tissue was harvested from ∼6-month-old plants. 100 mg wet weight of old leaf and berry tissue was harvested, 30 mg wet weight young leaf tissue, and 150-200 mg wet weight of root tissue was harvested. Harvested tissues were flash frozen in liquid nitrogen and ground to a fine powder in liquid nitrogen using a mortar and pestle. RNA was extracted using the Qiagen RNeasy Plant Mini Kit. Extracted RNA samples were flash frozen in liquid nitrogen and submitted to the Genome Technology Core at Whitehead Institute for Illumina paired-end RNA sequencing. RNA library prep was performed with xGen RNA Library Prep kit (IDT) and sequencing was carried out on a NovaSeqS4 PE150.

#### Genome annotation

Repeat-masking was performed on the *W. somnifera* genome using Earl Grey, a fully automated pipeline for transposable element annotation that identifies and masks known repeats using RepeatMasker, followed by analysis with RepeatModeler2 for de novo TE identification, followed by several more steps using other programs such as LTR finder^7^. Repeat analysis of the *W. somnifera* genome revealed that 81% of the genome consisted of repetitive elements, mostly classified as long terminal repeats (Supplementary Fig. 39). The repeat-masked genome was annotated using the BRAKER3 pipeline ^8–19^, a combination of GeneMark-ET and AUGUSTUS that uses a combination of user-provided transcriptomic and protein data to train the gene prediction tools. Raw RNA sequencing reads for all four tissue types (generated as described in RNA sequencing) were provided to BRAKER3 along with protein data from the Viridiplantae clade generated from OrthoDB v. 11^20^. This resulted in annotation of 36,057 genes. Synteny analyses were performed using MCScanX, MCScan (python version), and JVCI tools^21^.

#### RNA-seq data analysis

We performed RNA-seq read alignment to the *W. somnifera* genome using STAR, quantification using featureCounts, and count normalization using DESeq2. RNA-seq reads were aligned to the assembled *W. somnifera* genome using STAR (https://github.com/alexdobin/STAR), a splice-aware mapper^22^. Reads mapped to annotated genes were quantified using the featureCounts program in the Subread v. 2.0.1 package in fractional multi-mapping mode^23^. 84% of reads were successfully mapped to annotated features. The R package DESeq2^24^ was then used to log2 transform the read count data and normalize for factors such as library size. The heatmap was generated in R and the values were centered around zero.

Log2 values above 5 (32-fold overexpressed compared to the mean for a particular gene) or below -5 (32- fold underexpressed) appear solid red or blue, respectively, to aid in heatmap readability (Fig. 2c). Unadjusted heatmap available in Supplementary Fig. 40. Co-expression analysis was performed in R by computing Pearson correlation coefficients for every pairwise combination of genes in the biosynthetic gene clusters.

#### Viral induced gene silencing in *W. somnifera*

Gene silencing was performed using a tobacco rattle virus (TRV)-based VIGS system^25^, following the procedure described previously^26^. Inserts of interest were cloned into the pTRV2 vector at the *EcoRI* and *KpnI* restriction sites. The resulting constructs were introduced into *Agrobacterium tumefaciens* strain GV2260 harboring the GabT helper plasmid^27^. *Phytoene desaturase* (*PDS*) was used as positive control for silencing and a fragment of *GFP* as negative control. To induce silencing, 3-week-old *W. somnifera* seedlings were syringe-infiltrated with a 1:1 mixture of *Agrobacterium* cultures carrying pTRV1 and pTRV2. Six plants were infiltrated per construct, and successful gene silencing was verified by qPCR at harvest (∼4 weeks post-infiltration), coinciding with visible bleaching in *PDS*-silenced plants (∼3 weeks post-infiltration). The five uppermost leaves were collected from each plant and pooled for further analyses. Sterol extraction and analysis was done as described previously^26^.

#### Preparation of cDNA from *W. somnifera*

100 mg wet weight *W. somnifera* leaf or root tissue was harvested, flash frozen in liquid nitrogen, and ground to powder using a mortar and pestle in liquid nitrogen. The Qiagen RNeasy Plant Mini Kit was used to extract RNA and concentration was measured on the Nanodrop One Microvolume UV-vis Spectrophotometer (Thermo Fisher Scientific), then flash frozen with liquid nitrogen. cDNA was synthesized using the SuperScript™ III One-Step RT-PCR kit.

#### Metabolic engineering of *S. cerevisiae*

Candidate genes were amplified from *W. somnifera* cDNA using Q5 polymerase (NEB) or synthesized by Twist bioscience. Primers were used to add 30 bp homology arms for homology-based cloning. Candidate genes were cloned into yeast integration vectors from the EasyClone-MarkerFree Vector set^28^, a gift from Irina Borodina (Addgene kit #1000000098), using NEBuilder HiFi Assembly cloning (NEB). Candidate genes were placed under the GAL1 promoter, amplified from pYTK030 from the MoClo-YTK plasmid kit^29^, or under the bidirectional GAL1/10 promoter, amplified from pJK372 (gift from T. J. Kappock; Addgene plasmid # 73597; http://n2t.net/addgene:73597; RRID:Addgene_73597). The MoClo-YTK plasmid kit was a gift from John Dueber (Addgene kit # 1000000061). Assembled integration vectors were transformed into NEB DH5-α competent *E. coli.* Positive clones were verified by whole plasmid sequencing. Integration constructs were linearized using NotI-HF (NEB) digestion and genomically integrated following the lithium acetate/ssDNA/PEG yeast transformation method^30^. Colony PCR using primers from the EasyClone-MarkerFree Vector set^28^ were used to confirm site-specific integration at the desired locus.

#### Construction of Y7126 and Y7037

7-dehydrocholesterol reductase from *Solanum tuberosum* (*7RED*) and *24ISO* from *W. somnifera* were cloned into a LEU2 integration vector containing the GAL1/10 promoter assembled with the MoClo-YTK plasmid kit^29^. This integration vector was verified by whole plasmid sequencing and transformed into Y4108, with selection on synthetic defined (SD) medium without leucine for 3-4 days at 30 °C. Y4108 is a BY4742- derived strain constructed as described in a previous study^31^ (see Supplementary Table 3 for genotypes of all yeast strains). Correct integration of the *7RED*/*24ISO* construct was verified by colony PCR. *CYP87G1* from *W. somnifera* and the cytochrome P450 reductase *ATR1* from *Arabidopsis thaliana* were cloned into a URA3 integration vector containing GAL1/10 assembled with the MoClo-YTK plasmid kit^29^. The integration vector was verified by full plasmid sequencing and transformed into Y4108 to create strain Y6798, with selection on SD medium without uracil for 3-4 days at 30 °C and verification by colony PCR. This strain was subsequently transformed with the *7RED*/*24ISO* LEU2 integration construct described above and selected on SD medium without leucine for 3-4 days at 30 °C to create Y6852. To enable genome editing with CRISPR-Cas9, Y6852 was transformed with a hygromycin-marked plasmid containing the Cas9 under the TEF1 promoter (pcfB2312-Hyg). Correct transformants were selected on yeast peptone dextrose (YPD) medium with 200 μg/mL hygromycin B for 2-3 days at 30 °C. Colonies containing the plasmid were verified by colony PCR. pcfB2312-Hyg was adapted from the pcfB2312 plasmid from the EasyClone-MarkerFree Vector set^28^. The HygR resistance marker was amplified from the pYTK079 plasmid from the MoClo-YTK plasmid kit^29^.

#### Metabolite extraction from yeast

Yeast strains were struck on yeast peptone dextrose (YPD) plates with 200 μg/mL of hygromycin B and incubated for two days at 30 °C. Biomass from plates was used to inoculate 3 mL YPD cultures with 200 μg/mL hygromycin B and incubated overnight in a shaking incubator at 30 °C with 250 r.p.m shaking. Starter cultures were diluted into fresh 3 mL YPD cultures to an optical density at 600 nm (OD_600_) of 0.25. These were grown at 30 °C in a shaking incubator at 350 r.p.m for 48 h. To induce the cultures, they were centrifuged at 3000 rcf for 4 min, resuspended in 3 mL of sterile water to wash, centrifuged again, and resuspended in 3 mL of yeast peptone (YP) media with 2% galactose (YP-Gal). The cultures were incubated for 72 h at 30 °C with shaking at 350 r.p.m. Cell pellets were harvested by centrifugation at 3000 rcf for 4 min. For LC-MS analysis, pellets were resuspended with 500 μL water and transferred to 1.5 mL microcentrifuge tubes. 500 μL of ethyl acetate was added and samples were vortexed to mix. Samples were centrifuged at 21130 rcf to separate layers, and 250 μL of organic layer was transferred to a clean tube. 500 μL of ethyl acetate was added and samples were vortexed and centrifuged a second time. 500 μL of organic layer was removed and combined with the previously removed ethyl acetate extract. The combined extract was dried down under nitrogen or in a SpeedVac (Thermo Scientific) to evaporate solvent. Dried extracts were resuspended in 100 μL methanol, centrifuged at 21130 rcf for 5-10 mins, and 60 μL supernatant was aliquoted into LC-MS tubes. For GC-MS analysis, pellets were resuspended in 500 μL 40% potassium hydroxide in 50% methanol solution and transferred to 1.5 mL microcentrifuge tubes. Samples were incubated for 1 h at 90 °C in a heat block. 600 μL hexane was added to each sample, the samples were vortexed to mix, and centrifuged at 21130 rcf to separate layers. 500 μL of hexane layer was transferred to a clean set of tubes and this extraction process was repeated two additional times. The hexane extract was dried down under nitrogen and resuspended in 20 μL pyridine with 20 μL N-Methyl-N-trimethylsilyltrifluoroacetamide (MSTFA) to derivitize the sterols. The samples were incubated at room temperature for 20 mins and then dried down under nitrogen. The samples were resuspended with 100 μL hexane, centrifuged for 10 mins at 21130 rcf, and 50 μL of supernatant was transferred to glass GC-MS vials.

#### Transient expression in *Nicotiana benthamiana*

Candidate genes were amplified from *W. somnifera* cDNA or synthesized by Twist Bioscience and cloned into the pEAQ-HT vector^32^. Vectors were transformed into electrocompetent *Agrobacterium tumefaciens* LBA4404 by electroporation (2.2 kV, cuvettes with 0.1 cm gap). Transformants were selected on solid Luria-Bertani (LB) plates containing 50 μg/mL kanamycin, 100 μg/mL streptomycin, and 50 μg/mL rifampicin with incubation at 30 °C for 3 days. Positive clones were confirmed by colony PCR and struck on fresh plates. Biomass from plates was used to inoculate 5 mL liquid cultures of the same medium and grown at 30 °C with 250 r.p.m. shaking for ∼24 h. The cultures were centrifuged at 5000 rcf for 5 mins and resuspended in induction buffer (10mM MES buffer, 10mM MgCl_2_, and 150 μM acetosyringone, pH 5.6). Cultures were diluted to an OD_600_ of 0.4 using induction buffer and incubated at room temperature for 1 h with occasional inversion. Cultures of *Agrobacterium* strains containing plasmids to be co-infiltrated were then combined (4 mL per strain) and centrifuged at 5000 rcf for 5 mins. The supernatant was discarded and the cell pellet resuspended in 4 mL induction buffer. 4-6 week-old *N. benthamiana* plants were watered one day prior to infiltration. Plants were syringe infiltrated on the abaxial side of the leaves, then kept in the dark overnight. After ∼18 h, plants were watered and returned to the light and leaf tissue was harvested after five days. 100 ± 2 mg of fresh weight from each infiltrated leaf was added to 2mL microfuge tubes containing 2 mm zirconia/silica disruption beads, flash frozen in liquid nitrogen, and kept at -80 °C until sample processing. To extract metabolites for LC-MS analysis, 500 μL methanol was added to each sample. Samples were homogenized for 4 mins at 30 Hz in a Qiagen TissueLyser II, incubated at room temperature for 20 mins, and centrifuged for 2 mins at 21130 rcf. Supernatant was passed through 0.2 µm PTFE filters and aliquoted into LC-MS vials.

#### LC-MS analysis

Liquid chromatography was performed using a Vanquish Flex Binary UHPLC system equipped with a Vanquish Variable Wavelength Detector (Thermo Fisher Scientific). Solvent A was water with 0.1% formic acid and solvent B was acetonitrile with 0.1% formic acid. Reverse phase separation of analytes was performed on a Kinetex C18 column, 2.6 μm particle size, 150 x 3mm, 100Å (Phenomenex). The column oven was held at 30 °C and the flow rate was 0.5 mL/min. 2 µL of each sample was injected. MS analysis was performed on a high-resolution Orbitrap Exploris 120 benchtop mass spectrometer (Thermo Fisher Scientific). The default method for analyzing withanolide intermediates followed this gradient: 0-2 min, 40% B; 2-7 min, 40-95% B; 7-12 min 95% B; 12-15 min, 40% B. The electrospray ionization (H-ESI) source was operated in positive ionization mode with a full scan range of 300-600 *m/z* and data-dependent acquisition of MS/MS scans of the top four highest intensity precursor ions with dynamic exclusion. HCD collision energy was 30% for data-dependent MS/MS collection. The orbitrap resolution was 120,000 with RF lens of 70% and static spray voltage of 3500 V. To quantify sulfated compounds a modified version of the default method was occasionally used, in which the scan range was extended to 300-1000 *m/z* and the MS was operated in negative mode with a static spray voltage of 2500 V. For metabolic profiling of *W. somnifera* tissues, the default method was modified to the following gradient: 0-2 min, 40% B; 2-17 min, 40-95% B; 17-22 min 95% B; 22-25 min, 40% B. This method was run with MS collection in positive mode with a scan range of 300-1000 *m/z* and otherwise the same parameters as described for the default method or in negative mode with the same parameters as described above.

#### GC-MS analysis

GC-MS analysis was carried out on an Agilent 7890B GC system with an Agilent 5977 MSD and an Agilent HP-5MS column (5% Phenyl Methyl Silox, 30 m x 250 µm x 0.25 µm). Helium was used as the carrier gas with a flow rate of 1 mL/min and the injection volume was 1 µL at 300 °C (splitless mode). The oven program was 80 °C for 1 min, ramp at rate 20 °C/min to 300 °C, hold for 18 mins. Mass spectra were obtained with a scan range of 50-1000 *m/z*.

#### Recombinant protein expression and purification

SDH2 and SULF1 sequences were cloned into protein overexpression vector pHis8b containing an N-terminal 8xHis tag followed by a tobacco etch virus (TEV) cleavage site. Vectors were transformed into *E. coli* BL21 (DE3) for heterologous expression. Strains were inoculated from glycerol stock into 40 mL Terrific Broth (TB) with 50 µg/mL kanamycin in 250 mL flasks and cultivated overnight with incubation at 37 °C and shaking at 250 r.p.m. Starter cultures were diluted 50-fold into 6x 1 L TB cultures with 50 µg/mL kanamycin in 2.5 L Ultra Yield baffled flasks (Thomson). Flasks were incubated at 37 °C until OD_600_ of cultures reached 0.6-0.8. At that point cultures were induced with 0.15 mM IPTG and moved to incubators set to 18 °C with shaking at 250 r.p.m. After ∼16-18 h, cell pellets were harvested by spinning at 3000 rcf for 10 minutes in a floor centrifuge cooled to 4 °C. The cell pellets were resuspended in lysis buffer (25 mM Tris, 150 mM NaCl, 5% glycerol, 1 mg/mL lysozyme, 1 mM phenylmethylsulfonyl fluoride, 1 mM dithiothreitol (DTT), pH 8) and disrupted by sonication in an ice bath. Cell debris was removed by centrifugation at 20,000 r.p.m. for 40 mins at 4 °C. The supernatant was filtered using a 0.22 µm syringe filter unit and applied to a Ni-NTA column. The column was washed with four elution buffers (same components as lysis buffer but containing 20 mM, 50 mM, 100 mM, or 250 mM imidazole). Flow-through from the 250 mM imidazole elution was dialyzed overnight in 4 L dialysis buffer (25 mM Tris, 150 mM NaCl, 5% glycerol, pH 8) with purified his-tagged TEV at 4 °C. After dialysis the TEV-digested protein was applied to another Ni-NTA column to remove TEV and uncut protein. Flow-through was concentrated using Amicon Ultra-10 K centrifugal concentrator and final protein concentration was measured by bradford assay for SDH2 or by A_280_ on a NanoDrop for SULF1.

#### In vitro enzyme assays for SDH2 and SULF1 activity

In vitro assays to determine substrate preference of SDH2 were performed in a total volume of 200 µL with 1 µM SDH2, 2 mM NAD^+^, and extract from Y7567 in assay buffer 1 (20 mM potassium phosphate with 5 mM methyl beta cyclodextrin, pH 7.5). Reactions were vortexed to mix and incubated at 30 °C overnight. In vitro assays to determine cofactor preference of SDH2 were performed with 0.01 µM SDH2, extract from Y7567, and either 2 mM NAD^+^, 2 mM NADP^+^, or no cofactor added. Reactions were vortexed to mix and incubated at 30 °C for 45 mins. In vitro assays for SULF1 activity were performed in a total volume of 100 µL with 1 µM SULF1, 1 mM 3’-phosphoadenosine-5’-phosphosulfate (PAPS), and extract from Y7752 in assay buffer 2 (50% 20 mM potassium phosphate with 5 mM methyl beta cyclodextrin, pH 7.5 and 50% phosphate-buffered saline, pH 7.4). Reactions were vortexed to mix and incubated at 30 °C for 30 mins. In vitro SULF1 enzyme assays with purified **8** were performed in a total volume 200 µL with 20 µM SULF1, 1 mM PAPS, and ∼27 µg/mL **8** purified as described in ‘Purification of **8**’ in assay buffer 1. Reactions were vortexed to mix and incubated at 30 °C overnight. Assays were performed with n=3 or n=4 replicates. Reactions were quenched by the addition of 200 µL ethyl acetate and vortexing the sample. Samples were centrifuged at 21130 rcf to separate the organic layer and 150 µL of organic layer was transferred to a clean tube. An additional 200 µL ethyl acetate was added, the samples were vortexed and centrifuged as before, and 150 µL organic layer was removed and combined with the first extract. Samples were dried down under nitrogen, resuspended in 100 µL methanol, centrifuged at 21130 rcf, and 80 µL of supernatant was transferred to LC-MS vials.

#### CYP88C10 microsome assays

Microsomes were isolated from yeast strains Y7193 and Y7215. Strains were struck on YPD plates with 200 μg/mL of hygromycin B and incubated for 2 days at 30 °C and biomass was used to inoculate 3 mL YPD cultures with 200 μg/mL hygromycin B. The cultures were then incubated at 30 °C with shaking at 250 r.p.m. overnight. Cultures were diluted 1:10 into 200 mL YPD in flasks and incubated at 30 °C with shaking at 250 r.p.m. for 24 h. Cultures were centrifuged at 3000 rcf for 4 mins, washed with sterile water, and resuspended with 200 mL YP-Gal. Cultures were incubated at 30 °C with shaking at 250 r.p.m. for ∼18 h.

Cell pellets were collected by centrifugation at 3000 rcf for 10 min, resuspended in 100 mL sterile water to wash, and centrifuged again. Cell pellets were resuspended in 20 mL digestion buffer (1.1 M sorbitol, 50 mM potassium phosphate pH 7.5, 1 mM DTT, 2500 U lyticase per 100 mL of digestion buffer) and incubated for 1 h at 30 °C with shaking at 250 r.p.m. After digestion, cells were collected by centrifugation at 1200 rcf for 10 mins. Cell pellets were resuspended in cold lysis buffer (PBS buffer pH 7.5, 20% glycerol, 1 mM DTT, 2 mini Roche cOmplete™ Protease Inhibitor Cocktail tablets per 50 mL of lysis buffer) and transferred to 50 mL falcon tubes containing an approximately equal volume of 0.5 mm Zirconia beads. In a 4 °C cold room, the cell suspensions were vortexed 5-10 times for 1 min with 1 min breaks on ice in between. The supernatant was transferred to a new 50 mL falcon tube and centrifuged for 3000 rcf for 10 mins at 4 °C. The supernatant was transferred to ultracentrifuge tubes and centrifuged at 100,000 rcf for 45 min at 4 °C. After ultracentrifugation, the microsomal pellet was resuspended in 2 mL storage buffer (PBS buffer pH 7.5, 20% glycerol) using a syringe and needle, flash frozen in liquid nitrogen, and stored at -80 °C until use. Microsome assays were performed in a total volume of 100 µL with 2 mM NADPH and a concentration of 0.7-0.8 mg/mL total microsomal protein in assay buffer 1. Purified 7 generated as described in ‘Purification of 7’ was used as the substrate or a mixture of 7 and 11. To generate the mixture of 7 and 11, purified 7 was reacted with 20 µM SULF1 and 1 mM PAPS in assay buffer 2 in a total volume of 200 µL. The reaction was incubated for 1 h at 30 °C with shaking at 350 r.p.m. The reaction was quenched with 1:1 v/v ethyl acetate and products were extracted with ethyl acetate. Ethyl acetate extract was dried under nitrogen, resuspended in assay buffer 1, and this was used in the microsome assay. Reactions were incubated at 30 °C with shaking at 350 r.p.m. overnight. Reactions were quenched by the addition of 100 µL ethyl acetate and vortexing the sample. Samples were centrifuged at 21130 rcf to separate the organic layer and 80 µL of organic layer was transferred to a clean tube. An additional 100 µL ethyl acetate was added, the samples were vortexed and centrifuged as before, and 80 µL organic layer was removed and combined with the first extract. Samples were dried down under nitrogen, resuspended in 80 µL methanol, centrifuged at 21130 rcf, and 60 µL of supernatant was transferred to LC-MS vials.

#### Assay for spontaneous 10 production

We performed a large-scale SULF1 enzyme reaction to convert ∼2.5 mg purified 8 to a mixture of 9 and 10. The reaction was performed in a total volume of 6 mL, with 20 µM SULF1 and 1 mM PAPS in assay buffer 3 (20 mM potassium phosphate with 10 mM methyl beta cyclodextrin, pH 7.5). Reaction was quenched and products extracted by adding 1:1 v/v n-butanol, vortexing to mix, and separating layers by centrifugation at 4000 rcf for 3 mins. N-butanol layer was removed and transferred to a clean glass vial. This procedure was repeated three times and the n-butanol extracts were combined and dried under nitrogen. The dried extract was resuspended in 800 µL methanol and injected on a Shimadzu preparative HPLC with a Kinetex 5 μm C18 100 Å 150 x 21.2 mm column, AXIA packed (Phenomenex). Solvent A was water, solvent B was acetonitrile, and the gradient was 60-95% B over 1 h with a flow rate of 10 mL/min. The fractions containing 9 were dried down and this semi-pure 9 was resuspended in assay buffer 1 and immediately sampled and run on the LC-MS to verify that there was no 10 in the sample. The semi-pure 9 was then incubated overnight at 30 °C and sampled the next day.

#### Computational modeling of SDH2 and docking analysis

The SDH2 protein sequence, compound 7, and NADH were uploaded to the Boltz-1 (AlphaFold3) online webserver (https://neurosnap.ai/service/Boltz-1%20(AlphaFold3))^33^ to generate an AlphaFold-based SDH2 structural model in complex with compound 7 and NADH. Next, DynamicBind (https://neurosnap.ai/service/DynamicBind)^34^, a deep learning method designed for dynamic protein-ligand docking, was used to dock compound 7 into the Boltz-1-generated structure. The top five poses from DynamicBind were consistent with those from Boltz-1, providing validation for the results. The final docking model from DynamicBind was used to create the figure shown in the Supplemental Information.

#### General considerations for NMR

NMR spectra were recorded on a 700MHz Bruker Avance Neo NMR spectrometer equipped with a Prodigy CryoProbe.

#### Purification of 3

12 L of yeast strain Y7037 were cultured in 12x 2.5 L Ultra Yield baffled flasks (Thomson) with 1 L YPD in each flask. Cultures were inoculated at an initial OD_600_ of 0.25 and grown for two days at 30 °C with shaking at 180 r.p.m. Cultures were induced by media exchange to YP+2% galactose and grown for an additional four days at 30 °C with shaking at 180 r.p.m. Cell pellets were separated from the supernatant by centrifugation at 3000 rcf for 10 mins. Cell pellets were combined and resuspended in ∼1 L 50% methanol 40% potassium hydroxide in a 2 L glass beaker with a stir bar. The solution was stirred and heated to 90 °C on a hot plate for 1 h. After the solution was cool, 1:1 volume of hexane was added to the resuspended cell pellet, shaken in a separatory funnel, and the hexane layer was separated and collected. This process was repeated three times and evaporated by rotary evaporation to yield 143.4 mg extract. The combined supernatant was extracted by adding 1:1 volume of ethyl acetate in a separatory funnel, shaken in the funnel, and the ethyl acetate layer was collected. This process was repeated twice and the ethyl acetate extract was evaporated by rotary evaporation. Pellet and supernatant extracts were separately purified on a Buchi Pure C-815 Flash chromatography system. For the pellet extract, a Buchi FlashPure 4 g EcoFlex Silica 50 µm irregular column was used and for the supernatant extract, a Buchi FlashPure 25 g EcoFlex Silica 50 µm irregular column was used. Solvent A was hexane and solvent B was ethyl acetate. For the pellet extract the gradient was 0-100% (over 10 mins), then held at 100% for an additional 3.27 mins. The flow rate was 15 mL/min and the run was 33.2 column volumes (CVs). For the supernatant extract the gradient was 0-100% (over 22 mins), then held at 100% for an additional 3.75 mins. The flow rate was 32 mL/min and the run was 25.8 CVs. 100 µL aliquots of each fraction were dried down, resuspended in 100 µL methanol, and run on an LC-MS. Fractions containing **3** were combined and dried down. This extract was resuspended in (1:1) methanol-chloroform and separated on a Sephadex LH20 column eluted with 50% methanol 50% chloroform. The purest fraction containing **3** was dried down for a final weight of 3.6 mg, enabling structural confirmation by nuclear magnetic resonance spectroscopy (NMR).

#### Purification of 4

7 L of yeast strain Y7567 were cultured in 14x 2.5 L Ultra Yield baffled flasks (Thomson) with 500 mL YPD in each flask. Cultures were inoculated at an initial OD_600_ of 0.25 and grown for two days at 30 °C with shaking at 300 r.p.m. Cultures were induced by media exchange to YP+2% galactose and grown for an additional three days 30 °C with shaking at 300 r.p.m. Compounds were extracted by adding 1:1 volume of ethyl acetate to the yeast culture and shaking at 125 r.p.m. for 1 h at 30 °C. The ethyl acetate layer was separated by centrifugation and use of a separatory funnel. Yeast cell culture was extracted twice with ethyl acetate and the extract was evaporated by rotary evaporation. This process yielded 4.072 g of dried crude extract, which was purified using a Buchi FlashPure 25 g EcoFlex Silica 50 µm irregular column on a Flash chromatography system. Solvent A was hexane, solvent B was ethyl acetate, and the gradient was 0-100% (over 13 minutes) and kept at 100% for an additional 10 minutes. The flow rate was 32 mL/min and the run was 23 CVs. 100 µL aliquots of each fraction were dried down, resuspended in 100 µL methanol, and run on an LC-MS. Fractions containing **4** were combined and dried down, yielding 86.1 mg. This extract was resuspended in methanol and separated on a Sephadex LH20 column eluted with methanol, yielding 47.4 mg of extract. This extract was resuspended in methanol and injected into a Shimadzu preparative high-performance liquid chromatography (HPLC) with a Kinetex 5 µm PFP 100 Å 150 x 21.2 mm column, AXIA packed (Phenomenex). Solvent A was water with 0.1% TFA, solvent B was acetonitrile with 0.1% TFA, and the gradient was 40-95% B over 40 mins with a flow rate of 10 mL/min. The fraction containing **4** was dried down for a final weight of 3.2 mg, enabling structural confirmation by NMR.

#### Bioreactor fermentation for purification of 7, 8, and 10

For isolation of compounds **7**, **8**, and **10**, engineered yeast strains Y7587, Y7771, and Y7841 respectively were fermented in a 5 L BaoXing bioreactor (BIOTECH-5BG). The bioreactor was filled with 3 L YPD and inoculated with a 150-300 mL starter culture of the desired yeast strain to a starting OD_600_ of ∼0.25-0.5. The pH was controlled at 5 with 7.5 M ammonium hydroxide and the dissolved oxygen was controlled by stir rate and kept at 30-40%. A constant air flow of ∼0.8-1.6 L/min was used. Temperature was controlled at 30 °C. After ∼24 h, the bioreactor was induced with 40% concentrated galactose solution to a final concentration of ∼2% galactose. During the remainder of the fermentation, the bioreactor was batch fed with galactose whenever the pH began to spike, indicating sugar was depleted. Production was ended 7 days after induction. Compounds were extracted from the ∼3.6 L bioreactor culture by adding 1:1 volume of ethyl acetate to the yeast culture and shaking at 125 r.p.m. for 1 h at 30 °C. The ethyl acetate layer was separated by centrifugation and use of a separatory funnel. Yeast cell culture was extracted twice with ethyl acetate and the extract was evaporated by rotary evaporation.

#### Purification of 7

This process yielded 4.013 g of dried crude extract, which was purified using a Buchi FlashPure 25 g EcoFlex Silica 50 µm irregular column on a Flash chromatography system. Solvent A was hexane, solvent B was ethyl acetate, and the gradient was 0-100% (over 9 minutes) and kept at 100% for an additional 18 minutes. The flow rate was 32 mL/min and the run was 27 CVs. Fractions containing **7** were combined and dried down, yielding 22.63 mg. This extract was resuspended in methanol and separated on a Sephadex LH20 column eluted with methanol. The fractions containing **7** were combined and dried. This extract was resuspended in methanol and injected into a Shimadzu preparative HPLC with a Kinetex 5 μm C18 100 Å 150 x 21.2 mm column, AXIA packed (Phenomenex). Solvent A was water, solvent B was acetonitrile, and the gradient was 60-95% B over 1 h with a flow rate of 10 mL/min. The fractions containing **7** were dried down for a final weight of 5 mg, enabling structural confirmation by NMR.

#### Purification of 8

This process yielded 6.72 g of dried crude extract, which was purified using a Buchi FlashPure 25 g EcoFlex Silica 50 µm irregular column on a Flash chromatography system. The extract was eluted on an isocratic method using 100% ethyl acetate. Eluent containing **8** was dried down yielding 2.85 g. This extract was resuspended in methanol and separated on a Sephadex LH20 column eluted with methanol. The fractions containing **8** were combined and dried. This extract was resuspended in methanol and injected into a Shimadzu preparative HPLC with a Kinetex 5 μm C18 100 Å 150 x 21.2 mm column, AXIA packed (Phenomenex). Solvent A was water with 0.1% TFA, solvent B was acetonitrile with 0.1% TFA, and the gradient was 60-95% B over 60 mins with a flow rate of 10 mL/min. Eluent was re-run through the preparative HPLC a second time using a gradient of 60-80% B over 60 minutes followed by 80-95% B from 60-90 min. The fraction containing **8** was dried down for a final weight of 3.6 mg, enabling structural confirmation by NMR.

#### Purification of 10

This process yielded 5.97 g of dried crude extract, which was purified using a Buchi FlashPure 25 g EcoFlex Silica 50 µm irregular column on a Flash chromatography system. Solvent A was hexane, solvent B was ethyl acetate, and the gradient was 0-100% (over 13 minutes) and kept at 100% for an additional 8 minutes. The flow rate was 32 mL/min and the run was 21 CVs. Fractions containing **10** were combined and dried down and this extract was resuspended in methanol and separated on a Sephadex LH20 column eluted with methanol. The fractions containing **10** were combined and dried. This extract was resuspended in methanol and injected into a Shimadzu preparative HPLC with a Kinetex 5 μm C18 100 Å 150 x 21.2 mm column, AXIA packed (Phenomenex). Solvent A was water, solvent B was acetonitrile, and the gradient was 60-95% B over 1 h with a flow rate of 10 mL/min. The fractions containing **10** were dried down for a final weight of 1.94 mg, enabling structural confirmation by NMR.

## Supplemental figures

**Supplementary Fig. 1.**
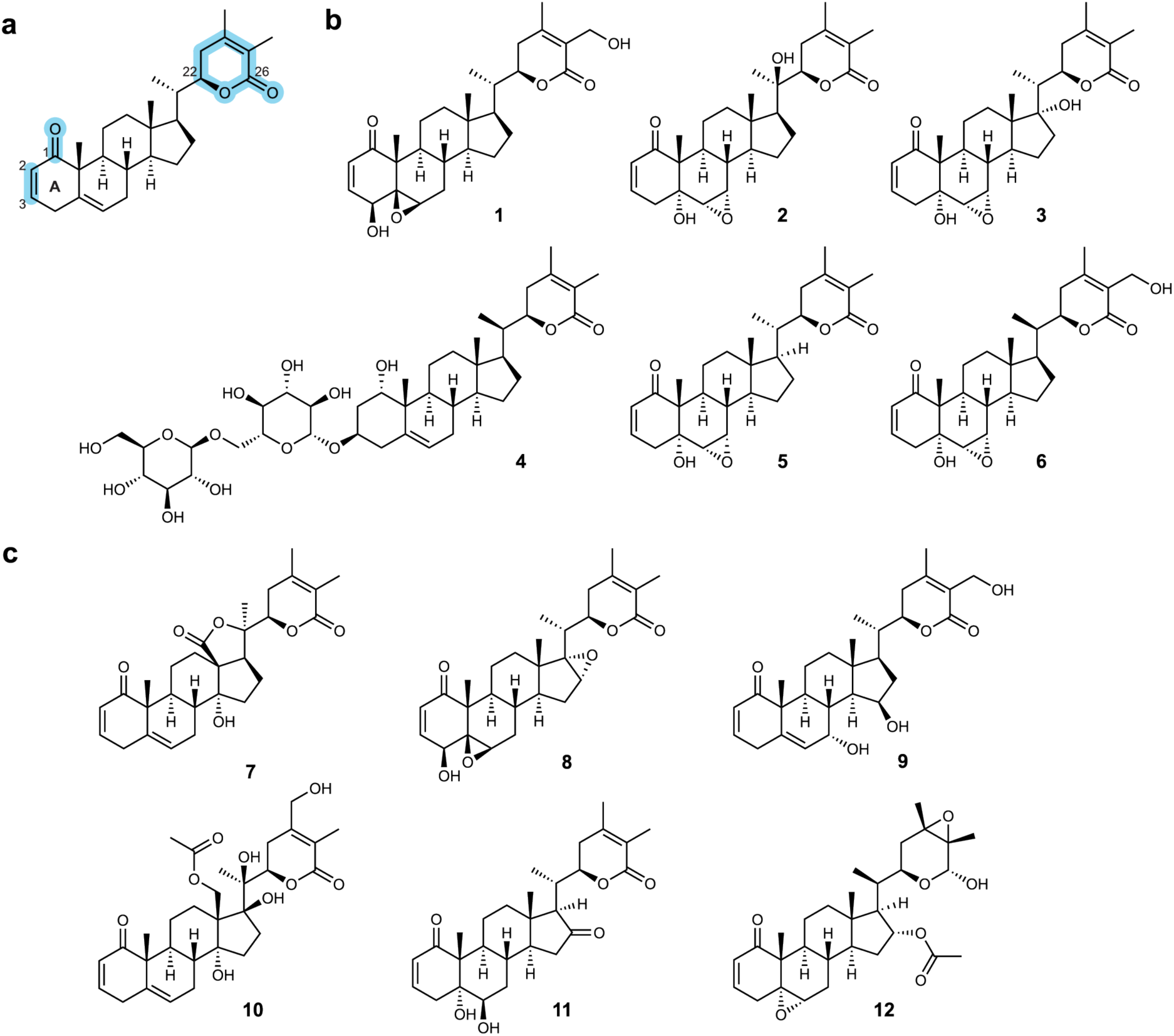
Withanolide core structure and select withanolides from *W. somnifera* and other Solanaceae plants. **a**, Schematic showing the core withanolide structure including the key chemical features of C_2_-C_3_ unsaturation, C_1_ ketone, and lactone ring (highlighted blue). **b**, Examples of withanolides identified in *W. somnifera* including withaferin A (**1**), withanolide A (**2**), withanone (**3**), withanoside V (**4**), withanolide B (**5**), and 12-deoxywithastramonolide (**6**). **c**, Examples of withanolides identified from various plants in the Solanaceae, including withaphysalin A from *Physalis minima* (**7**), tubocapsanolide A from *Tubocapsicum anomalum* (**8**), daturmetelide F from *Datura metel L.* (**9**), orizabolide from *Physalis orizabae* (**10**), 16-oxo-27-deoxyjaborosalactone D from *Nicandra john-tyleriana*, and salpichrolide O from *Salpichroa origanifolia* (**12**).

**Supplementary Fig. 2.**
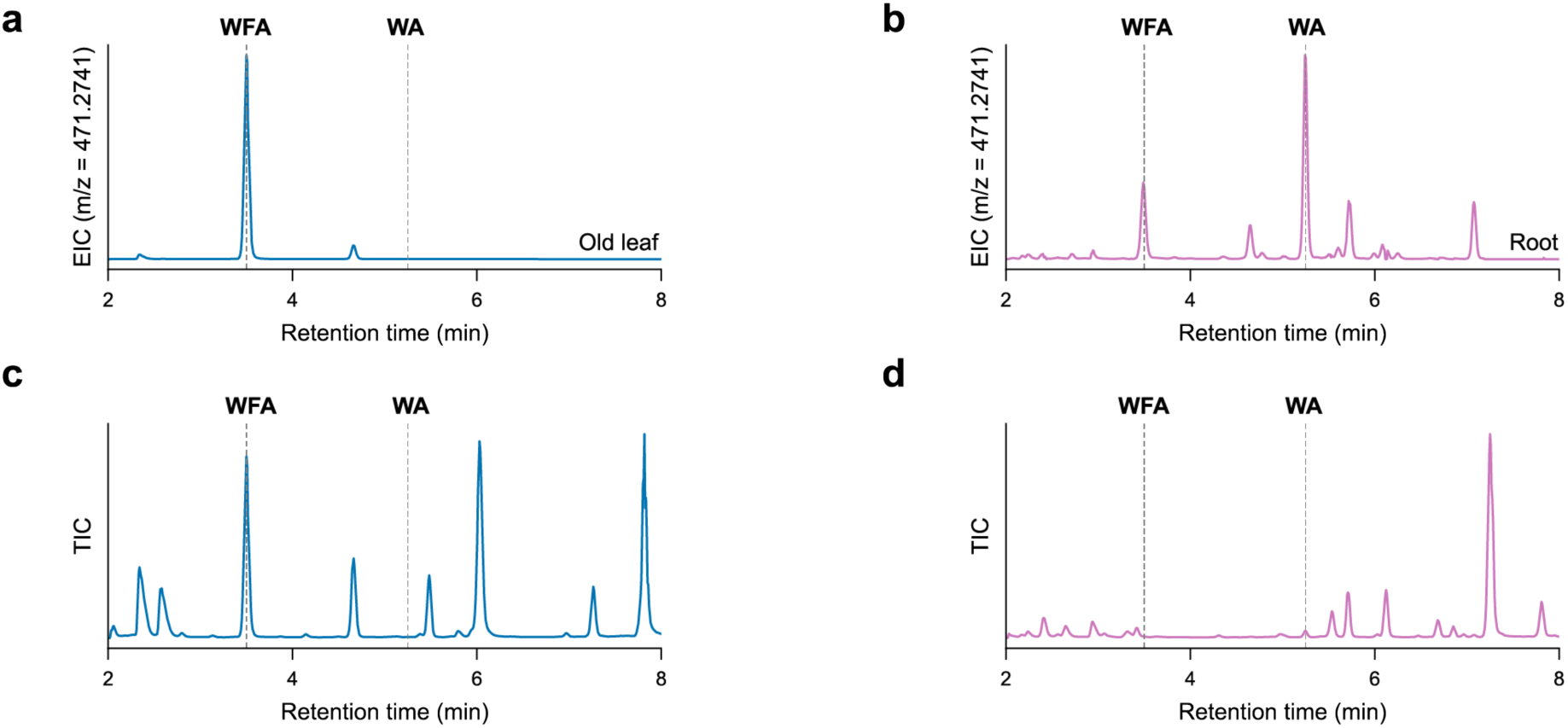
Metabolic profiling of *W. somnifera* old leaf and root tissue. **a**, Methanolic extract of *W. somnifera* old leaf tissue. Extracted ion chromatogram (EIC) for [M+H]^+^ ion corresponding to withaferin A (WFA) and withanolide A (WA) mass (C_28_H_38_O_6_; *m/z* = 471.2741). **b**, Methanolic extract of *W. somnifera* root tissue. EIC for [M+H]^+^ ion corresponding WFA and WA mass (C_28_H_38_O_6_; *m/z* = 471.2741). **c**, Methanolic extract of *W. somnifera* leaf tissue. Liquid chromatography-mass spectrometry (LC-MS) total ion chromatogram (TIC) with WFA and WA retention times indicated. **d**, Methanolic extract of *W. somnifera* root tissue. LC-MS TIC with WFA and WA retention times indicated.

**Supplementary Fig. 3.**
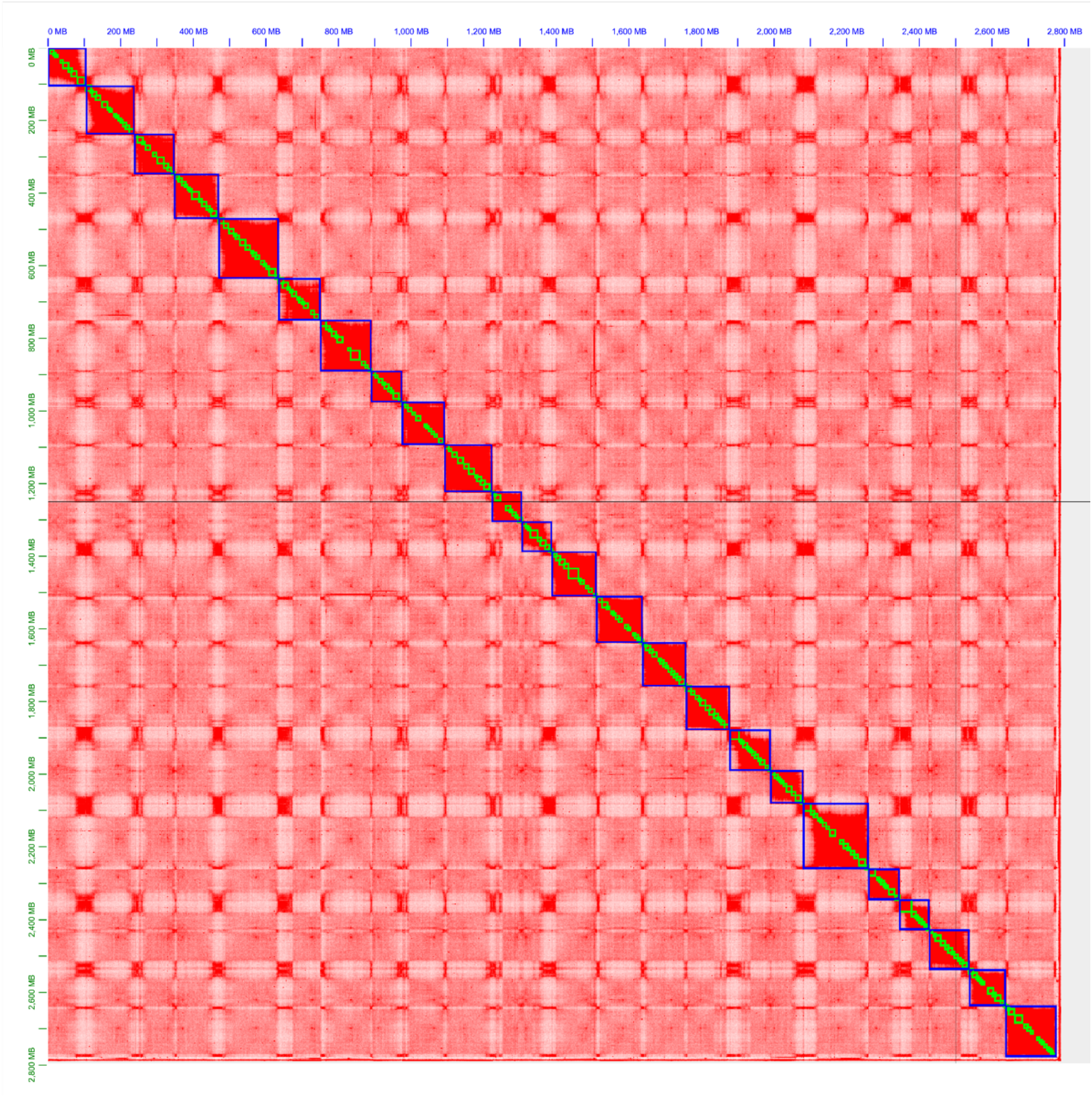
Hi-C contact heatmap of *W. somnifera* genome. Pseudochromosomes demarcated by blue squares and scaffolds demarcated by green squares. Contact density indicated by color (white=low, red=high). Heatmap generated in Juicebox at a resolution of 2.5 Mbp.

**Supplementary Fig. 4.**
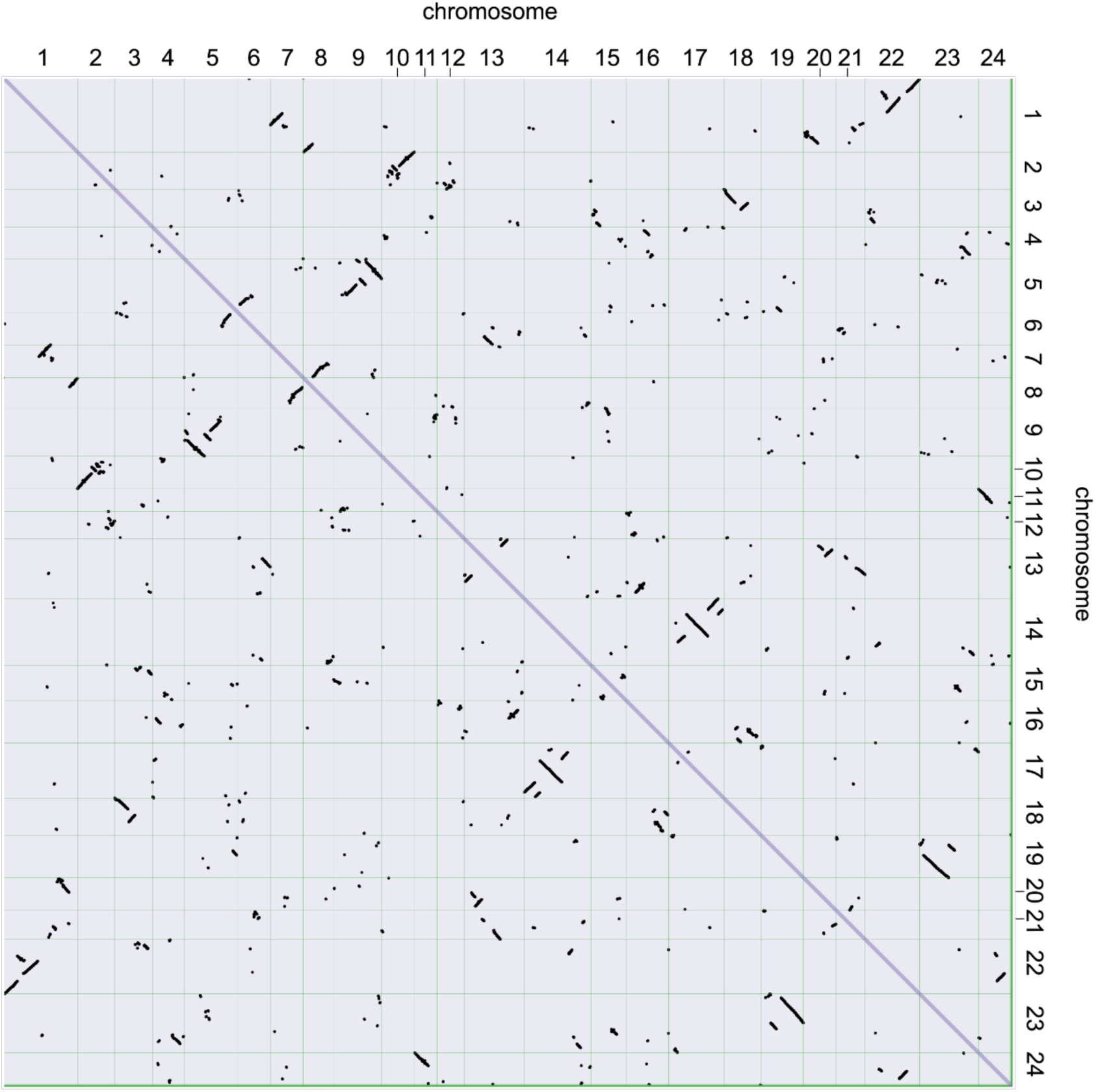
Intra-genomic synteny dot plot of *W. somnifera* genome. Syntenic gene pairs indicated by dots (9,570 gene pairs in total).

**Supplementary Fig. 5.**
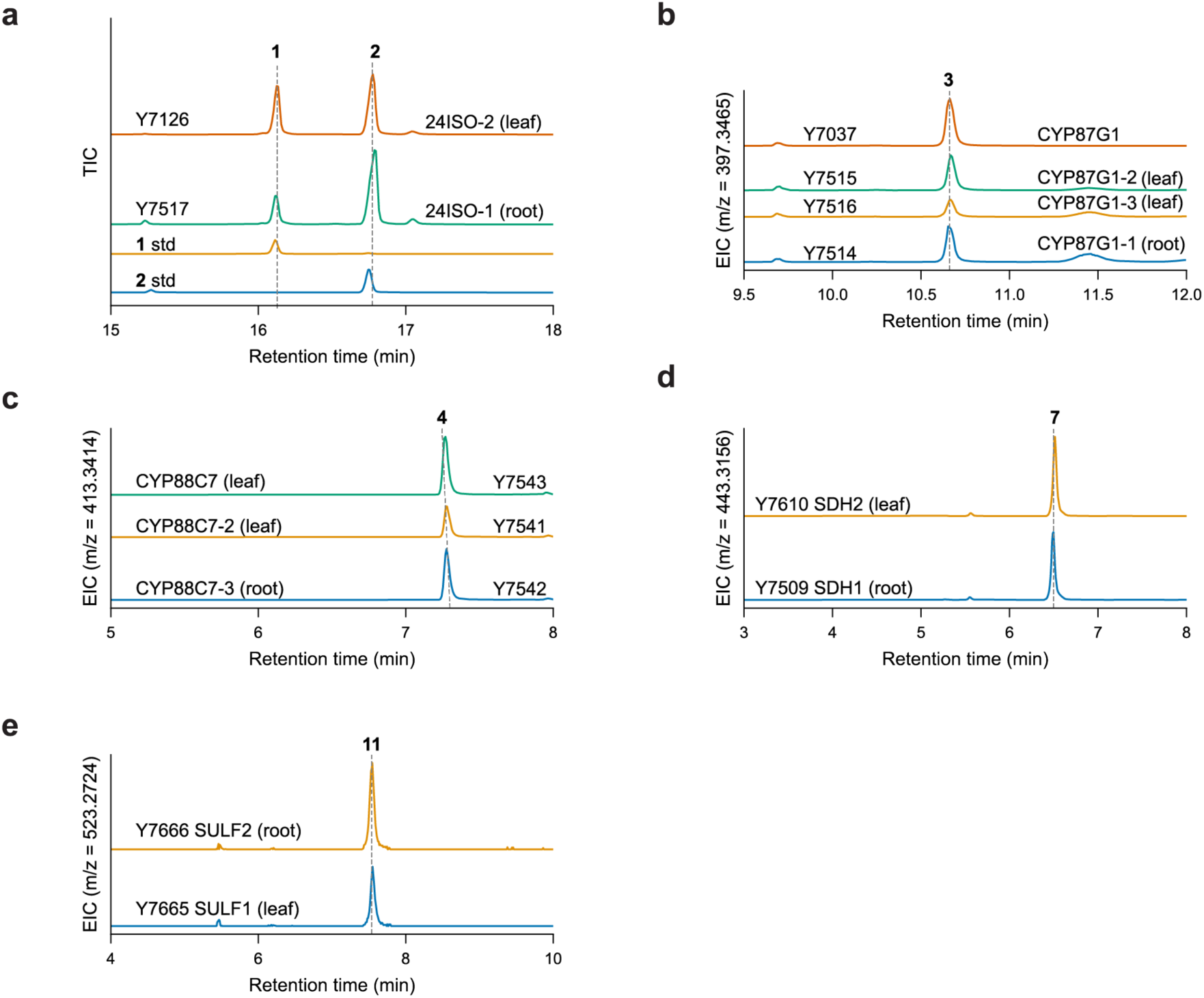
Leaf- and root-expressed paralogs of *24ISO*, *CYP87G1*, *CYP88C7*, *SDH*, and *SULF* make the same products. **a**, Gas chromatography-mass spectrometry (GC-MS) TICs of yeast strains expressing leaf and root *24ISO* paralogs. Authentic standard for 24-methylene cholesterol (**1**) and [26-^13^C]-labeled standard for 24-methyldesmosterol (**2**). **b**, Products of yeast strains expressing leaf and root *CYP87G1* paralogs. EIC for **3** ([M-H_2_0+H]^+^; *m/z* = 397.3465). Experiment performed in triplicate and representative traces are shown. **c**, Products of yeast strains expressing leaf and root *CYP88C7* paralogs. EIC for **4** ([M-H_2_0+H]^+^; *m/z* = 413.3414). Experiment performed in triplicate and representative traces are shown. **d**, Products of yeast strains expressing leaf and root *SDH* paralogs. EIC for **7** ([M+H]^+^; *m/z* = 443.3156). Experiment performed in duplicate and representative traces are shown. **e**, Products of yeast strains expressing leaf and root *SULF* paralogs. EIC for **11** ([M+H]^+^; *m/z* = 523.2724). Experiment performed in duplicate and representative traces are shown.

**Supplementary Fig. 6.**
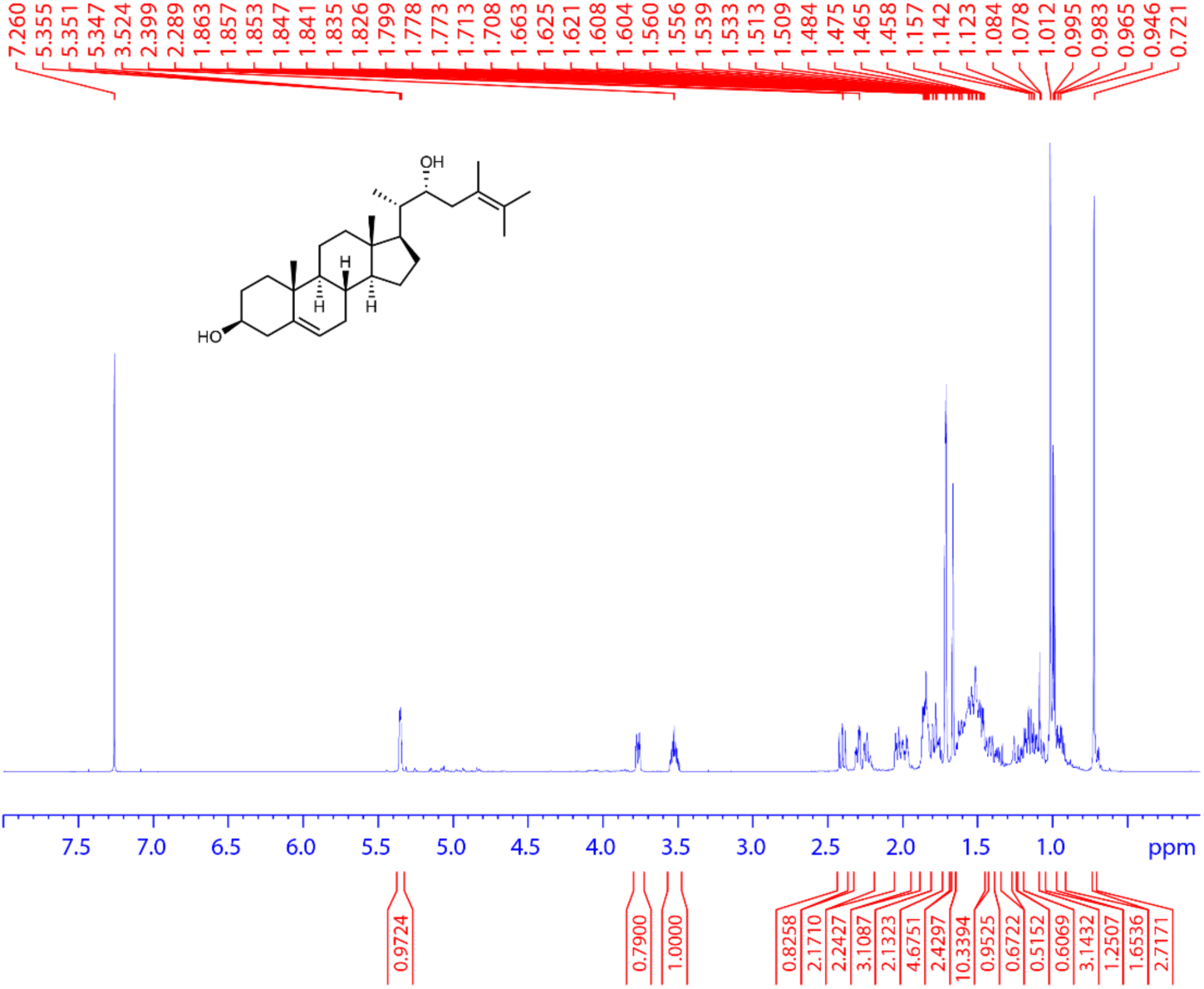
**^1^H spectrum of 3. (700 MHz, CDCl_3_, 298 K).**

**Supplementary Fig. 7.**
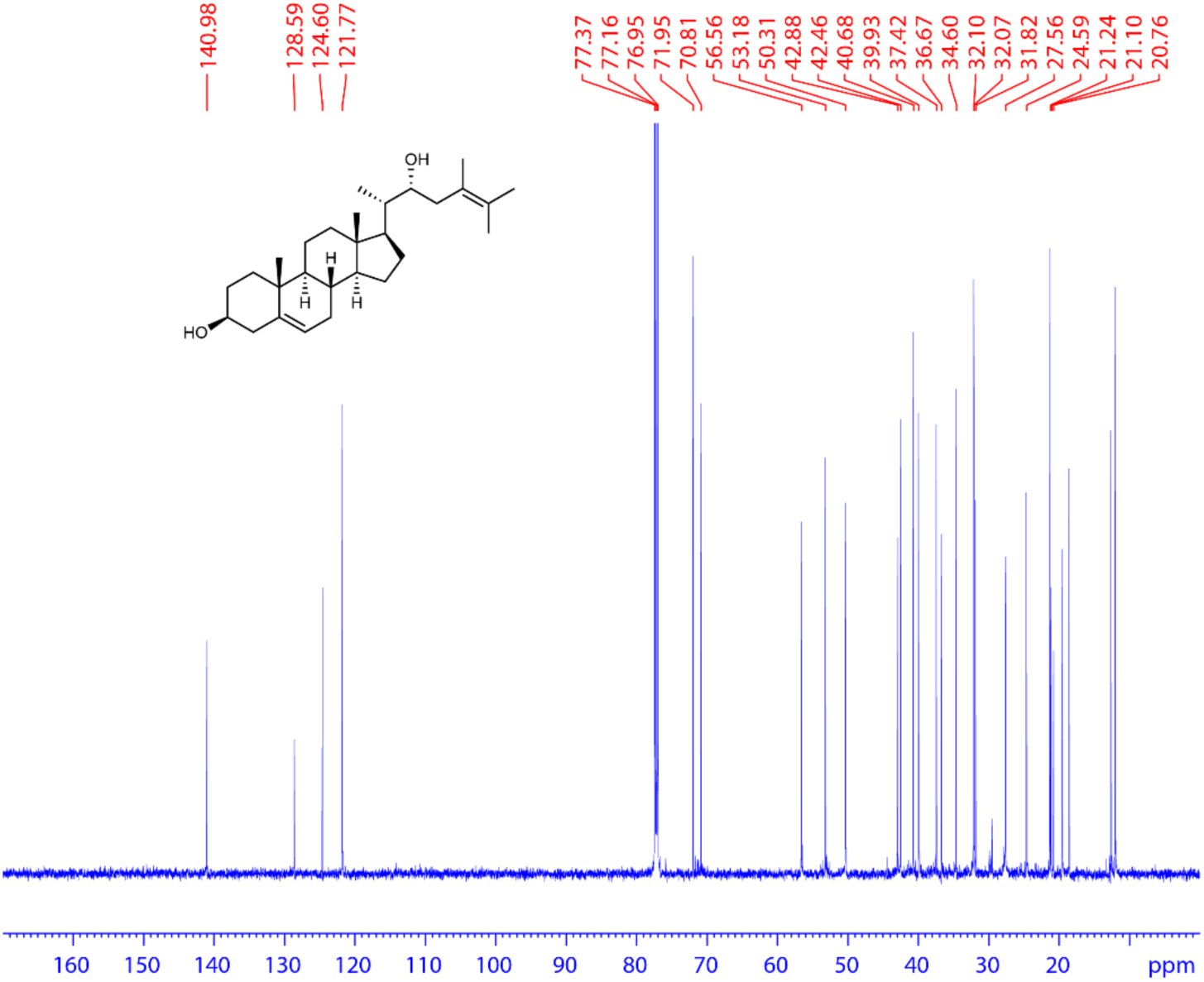
**^13^C spectrum of 3. (175 MHz, CDCl_3_, 298 K).**

**Supplementary Fig. 8.**
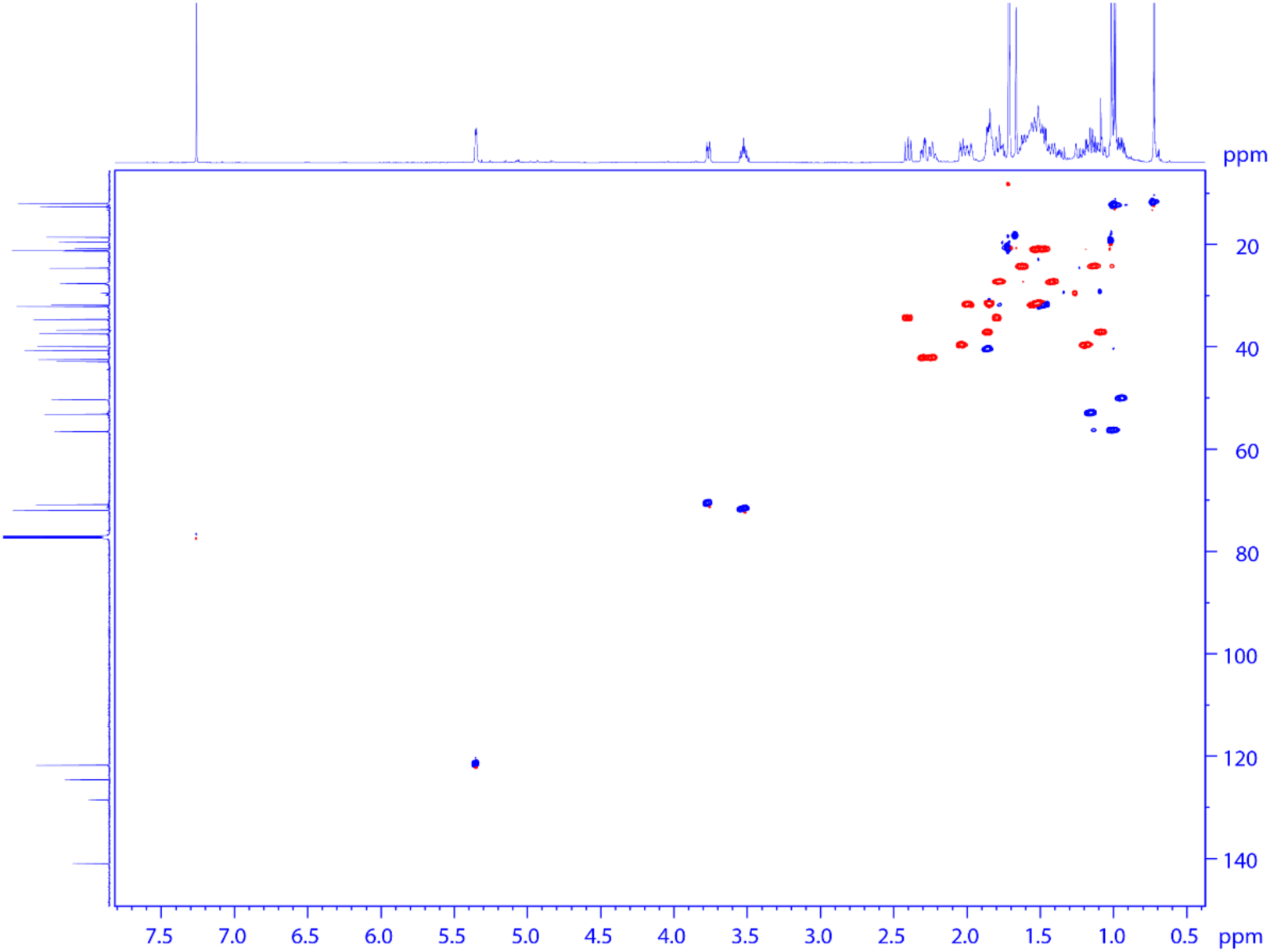
**HSQC spectrum of 3.**

**Supplementary Fig. 9.**
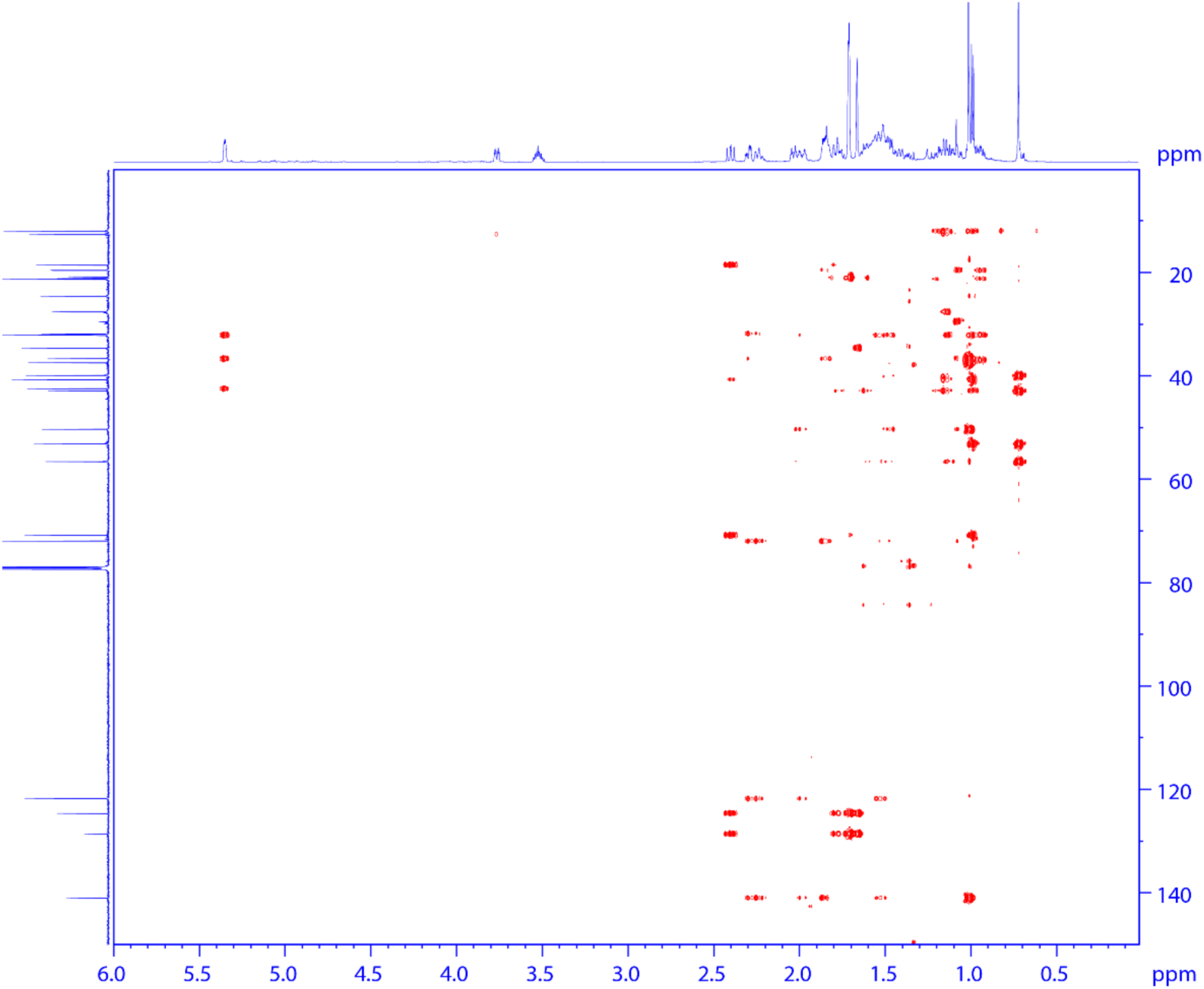
**HMBC spectrum of 3.**

**Supplementary Fig. 10.**
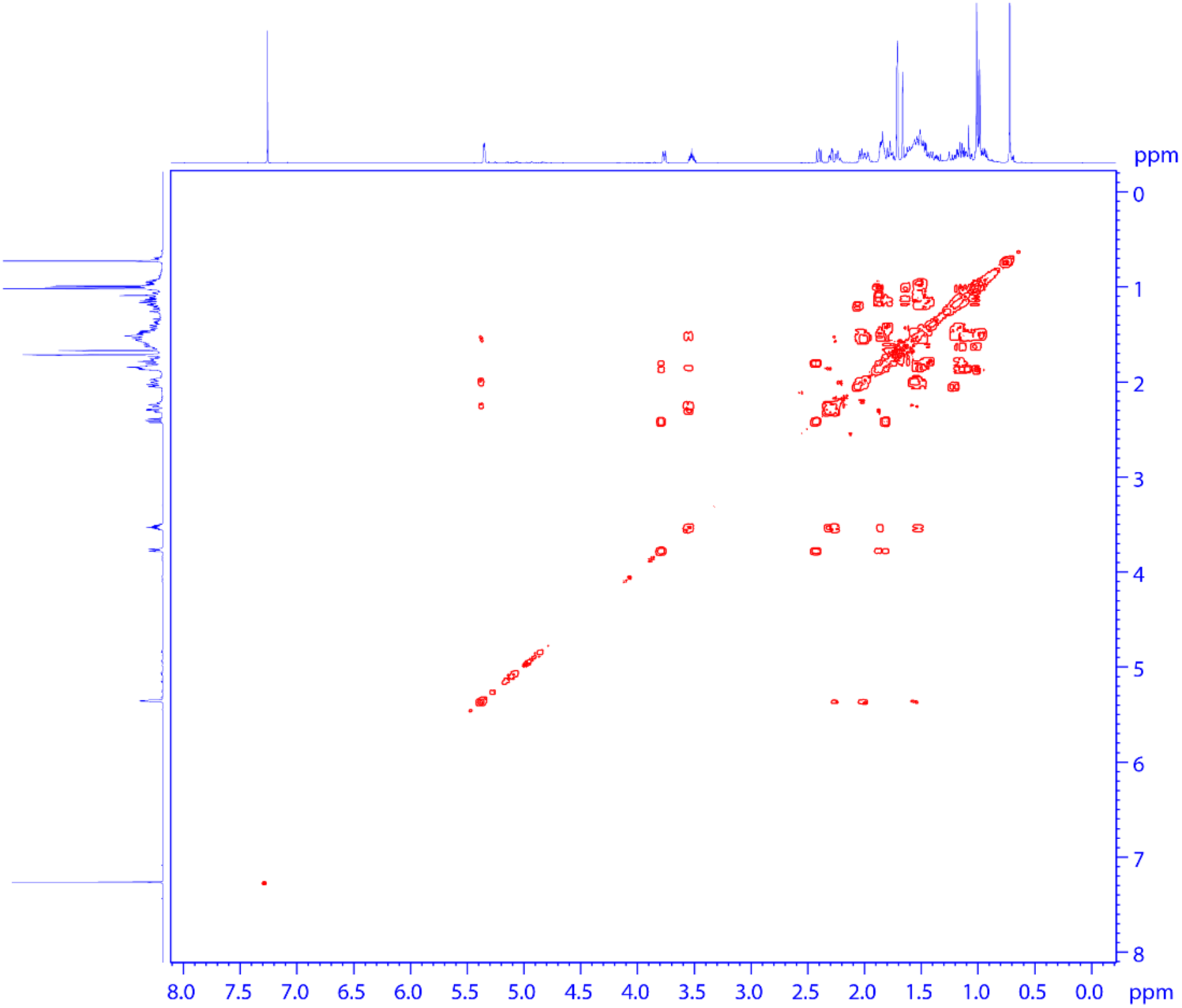
**HHCOSY spectrum of 3.**

**Supplementary Fig. 11.**
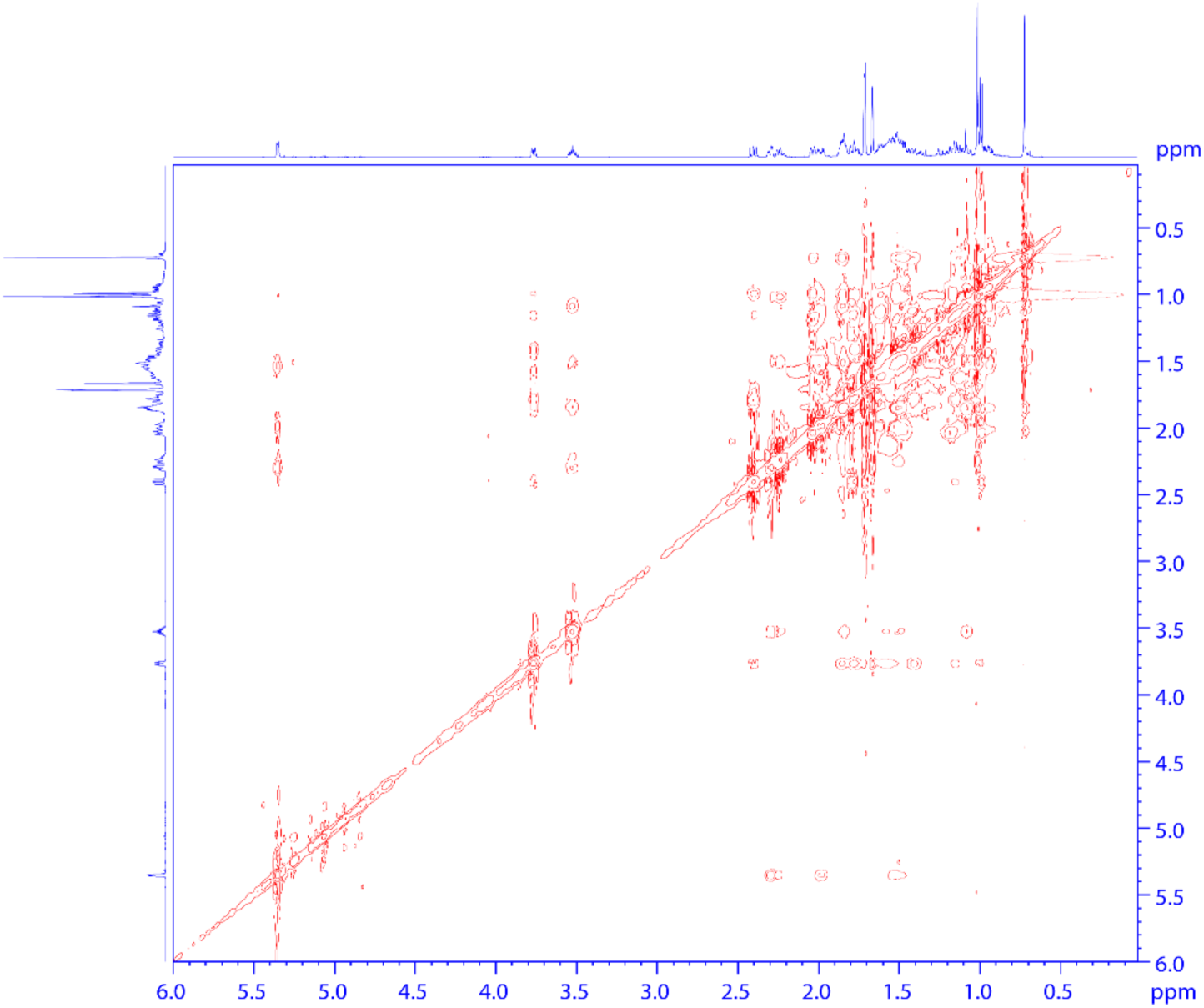
**NOESY spectrum of 3.**

**Supplementary Fig. 12.**
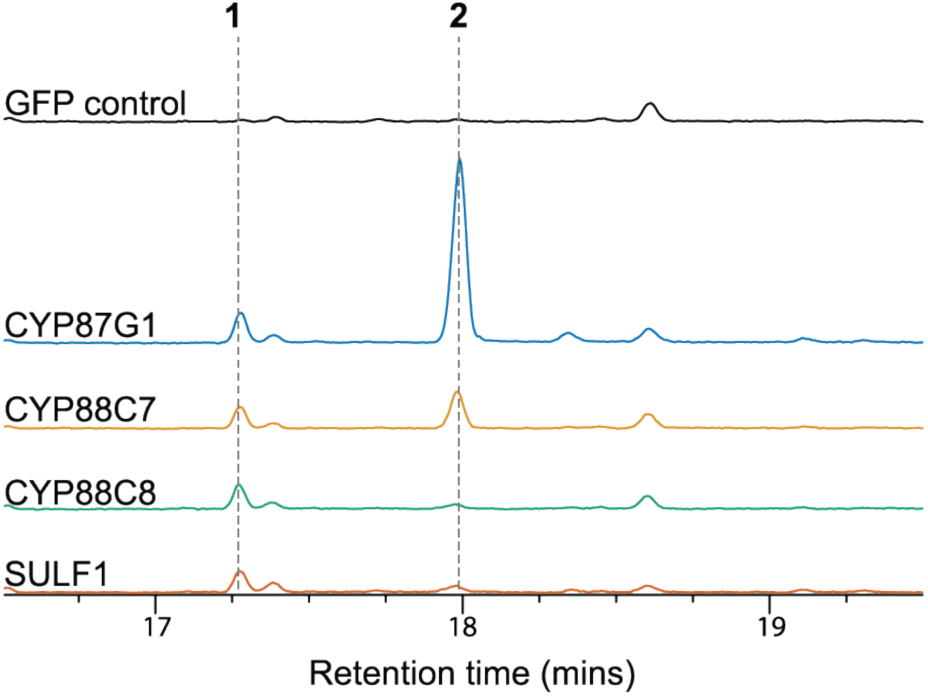
Viral-induced gene silencing of withanolide biosynthetic genes in *W. somnifera*. GC-MS TICs for samples from *W. somnifera* plants that had various withanolide biosynthetic genes silenced by viral-induced gene silencing (VIGS). Experiment was performed in triplicate and representative traces chosen. **1** is 24-methylene cholesterol and **2** is 24-methyldesmosterol.

**Supplementary Fig. 13.**
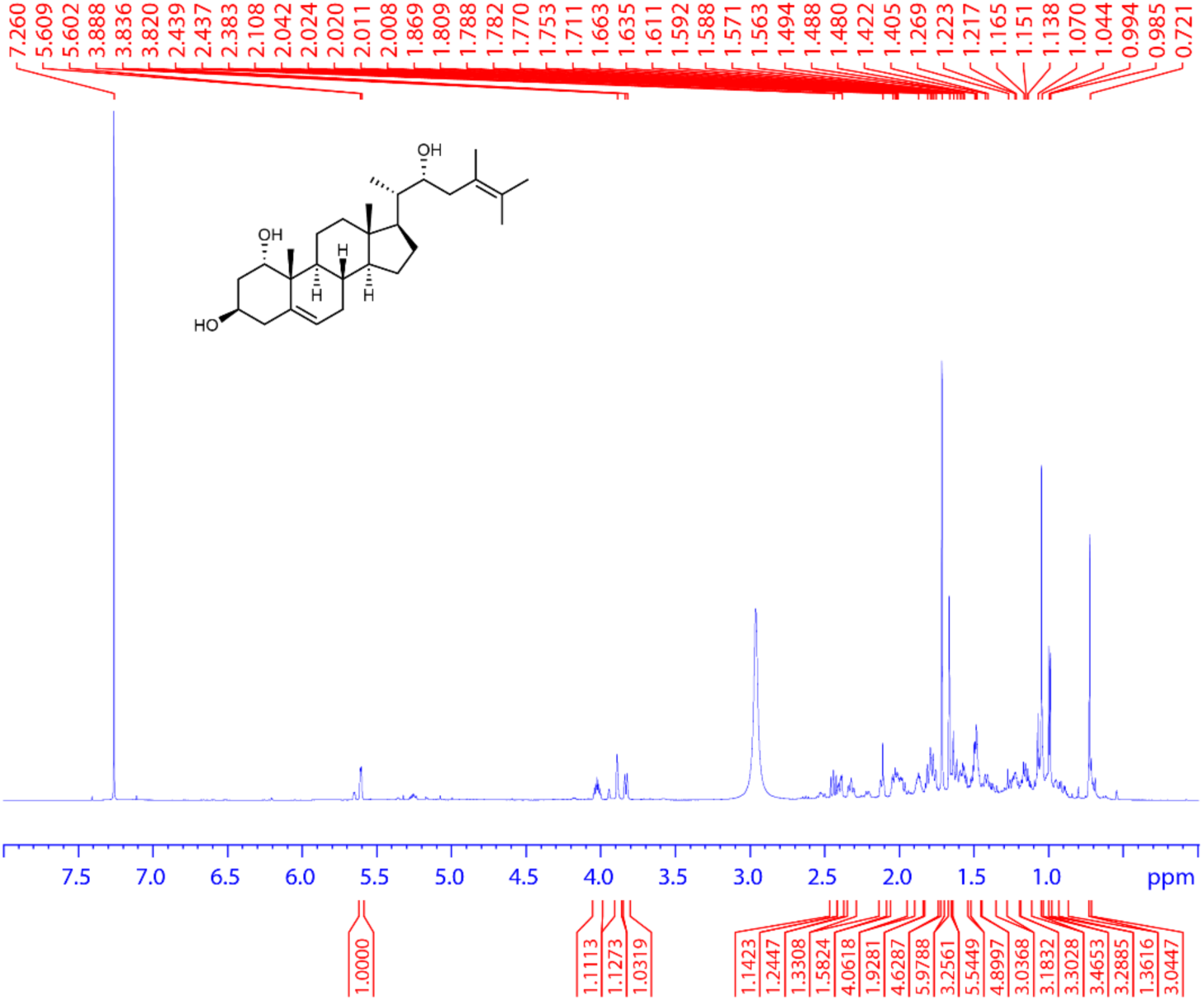
**^1^H spectrum of 4. (700 MHz, CDCl_3_, 298 K).**

**Supplementary Fig. 14.**
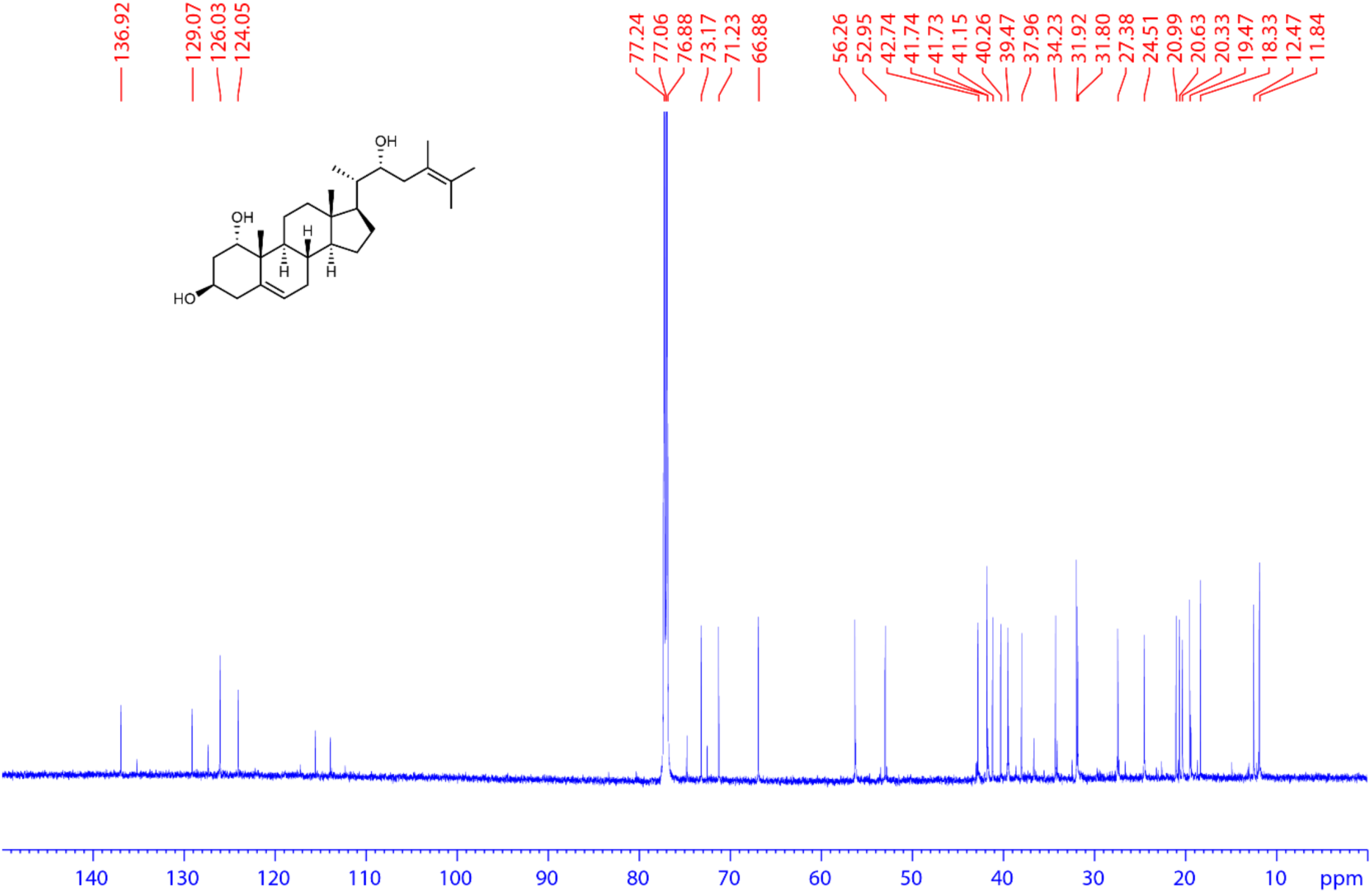
**^13^C spectrum of 4. (175 MHz, CDCl_3_, 298 K).**

**Supplementary Fig. 15.**
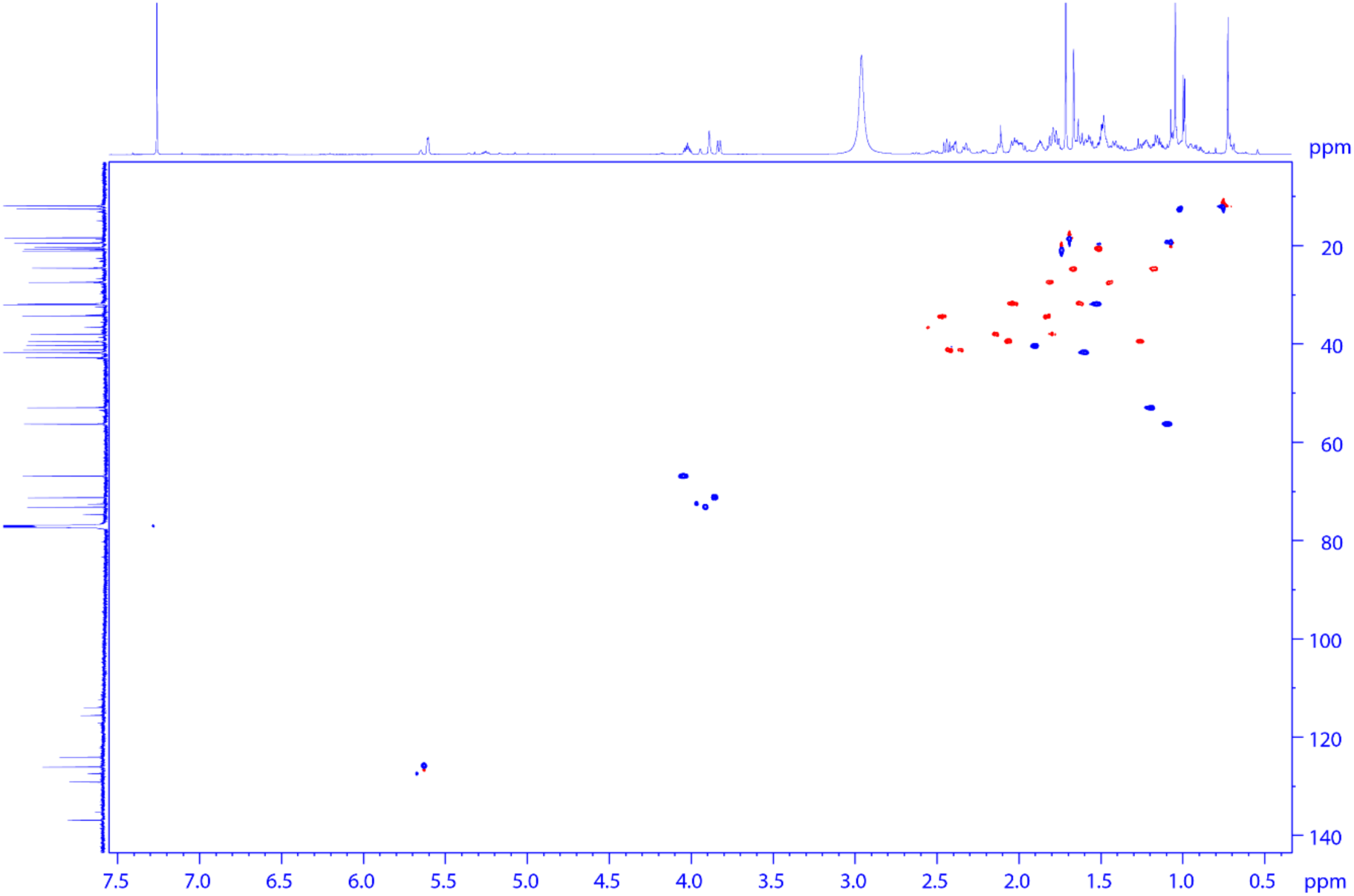
**HSQC spectrum of 4**

**Supplementary Fig. 16.**
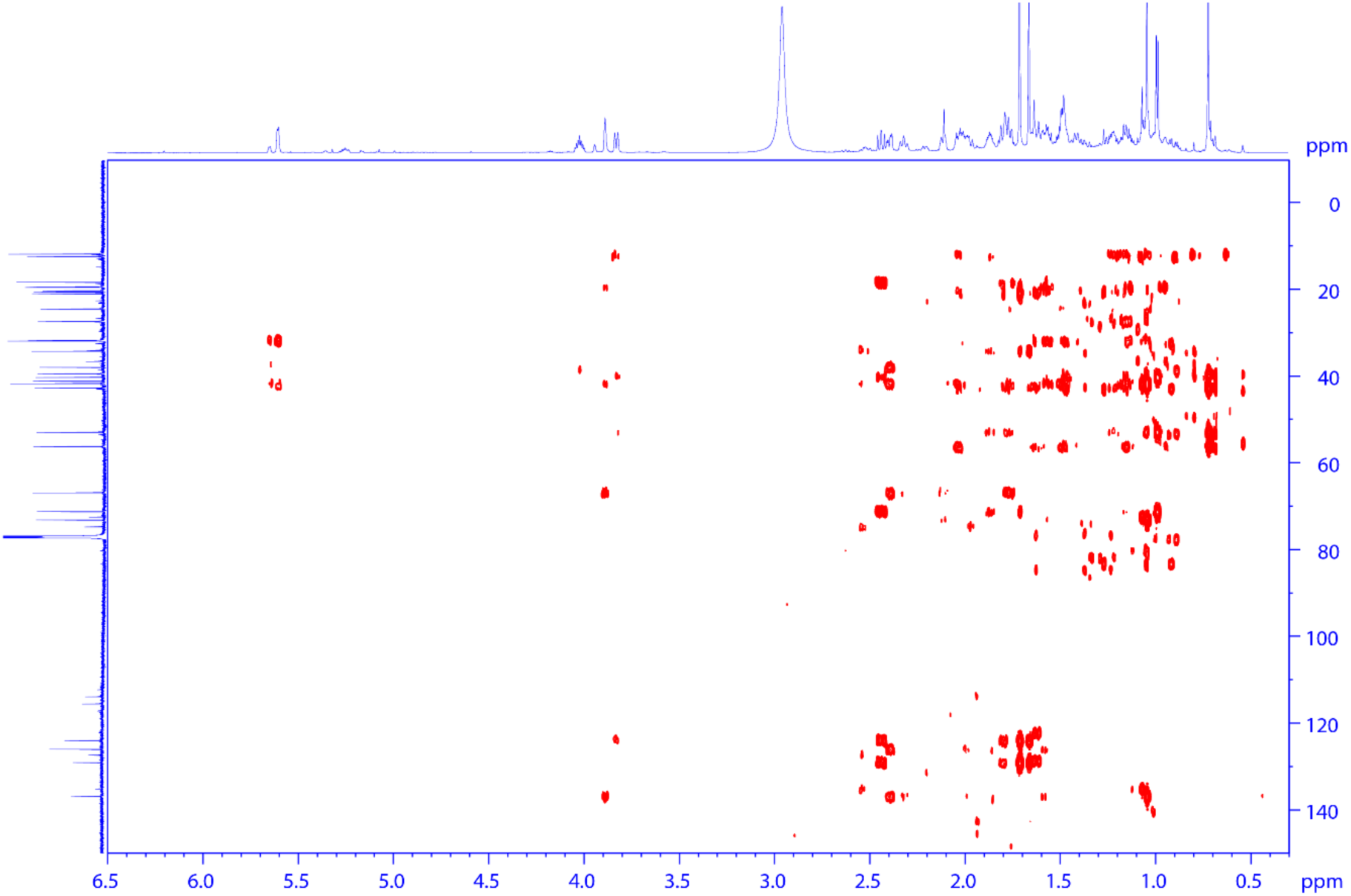
**HMBC spectrum of 4.**

**Supplementary Fig. 17.**
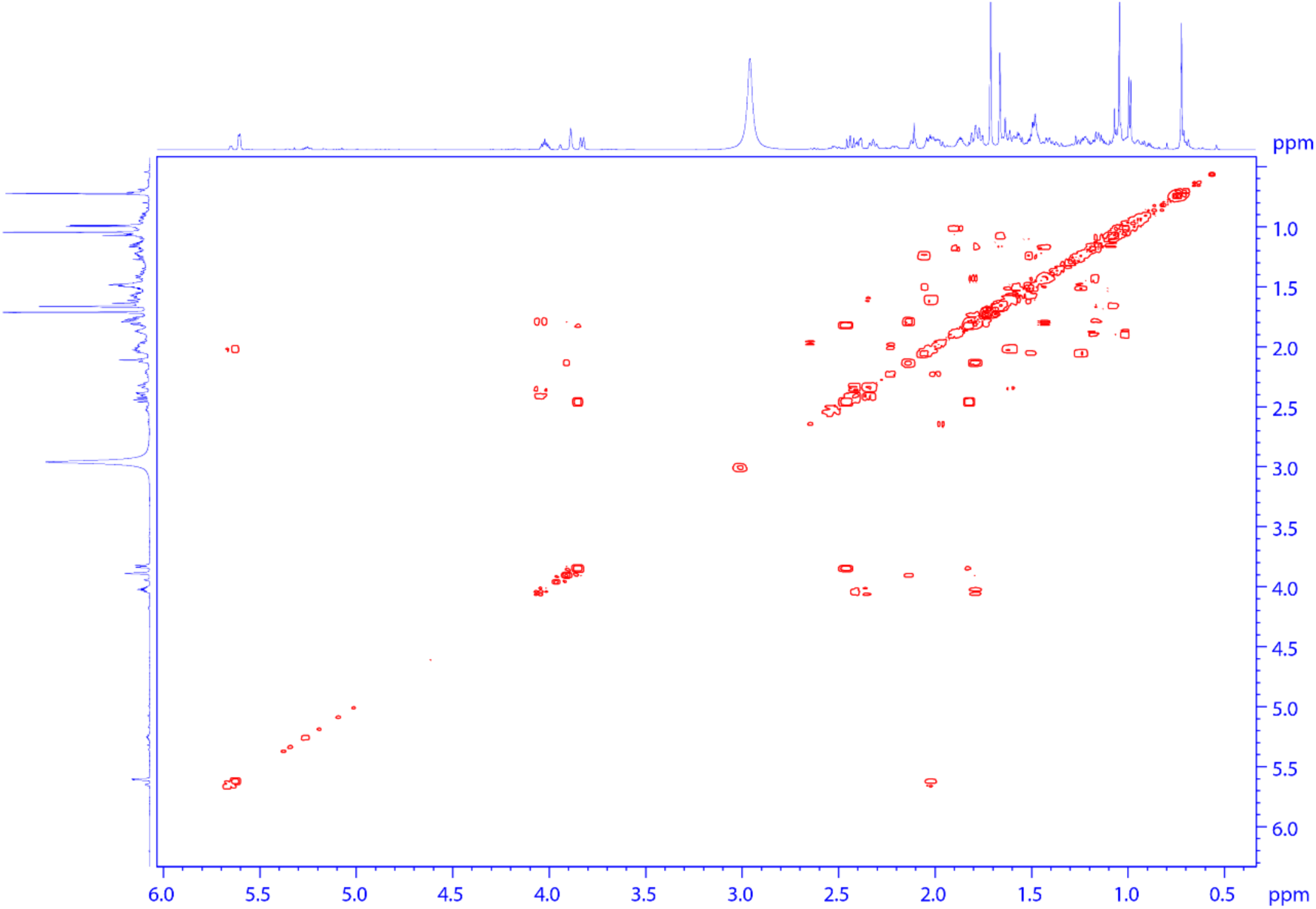
**HHCOSY spectrum of 4**

**Supplementary Fig. 18.**
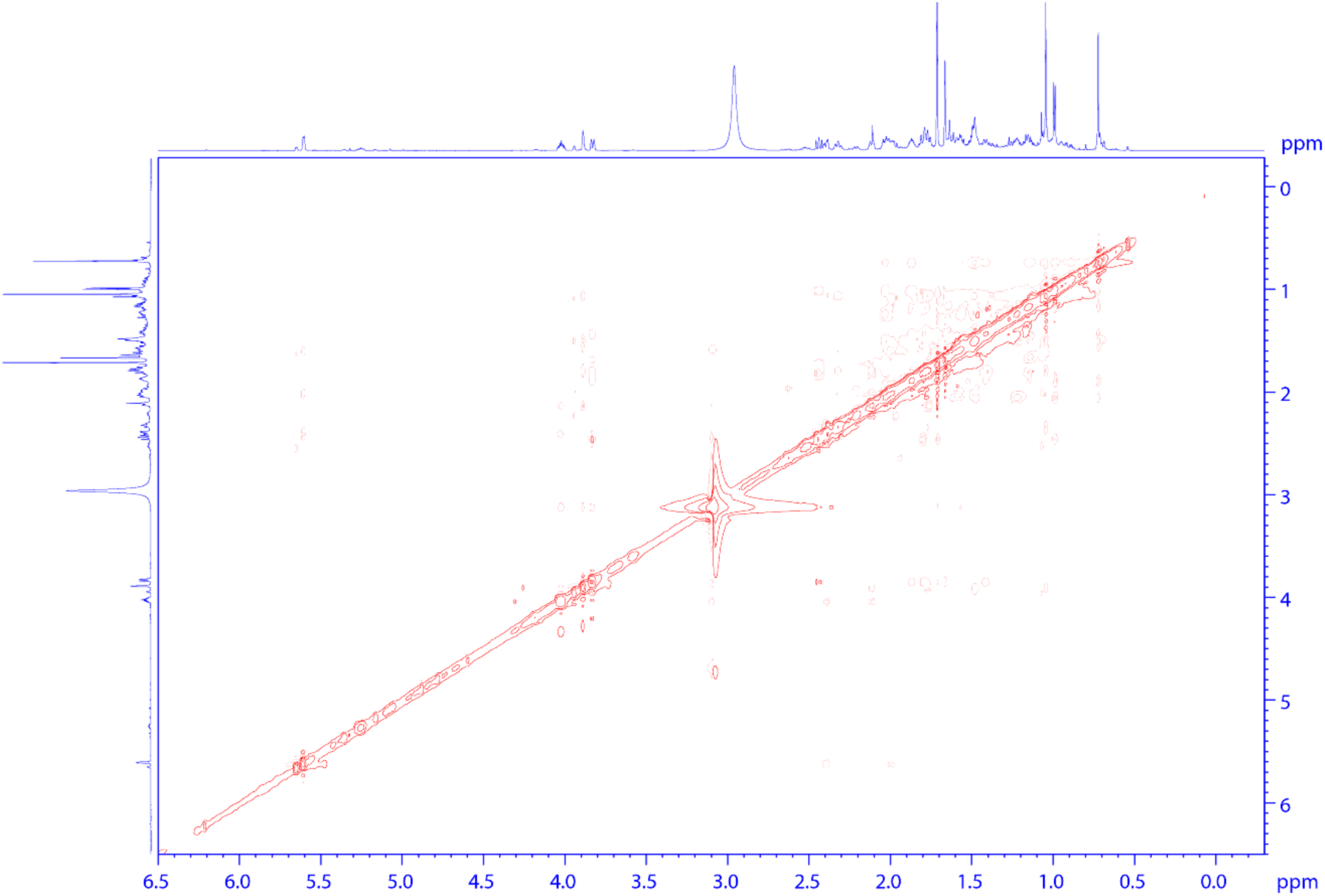
**NOESY spectrum of 4.**

**Supplementary Fig. 19.**
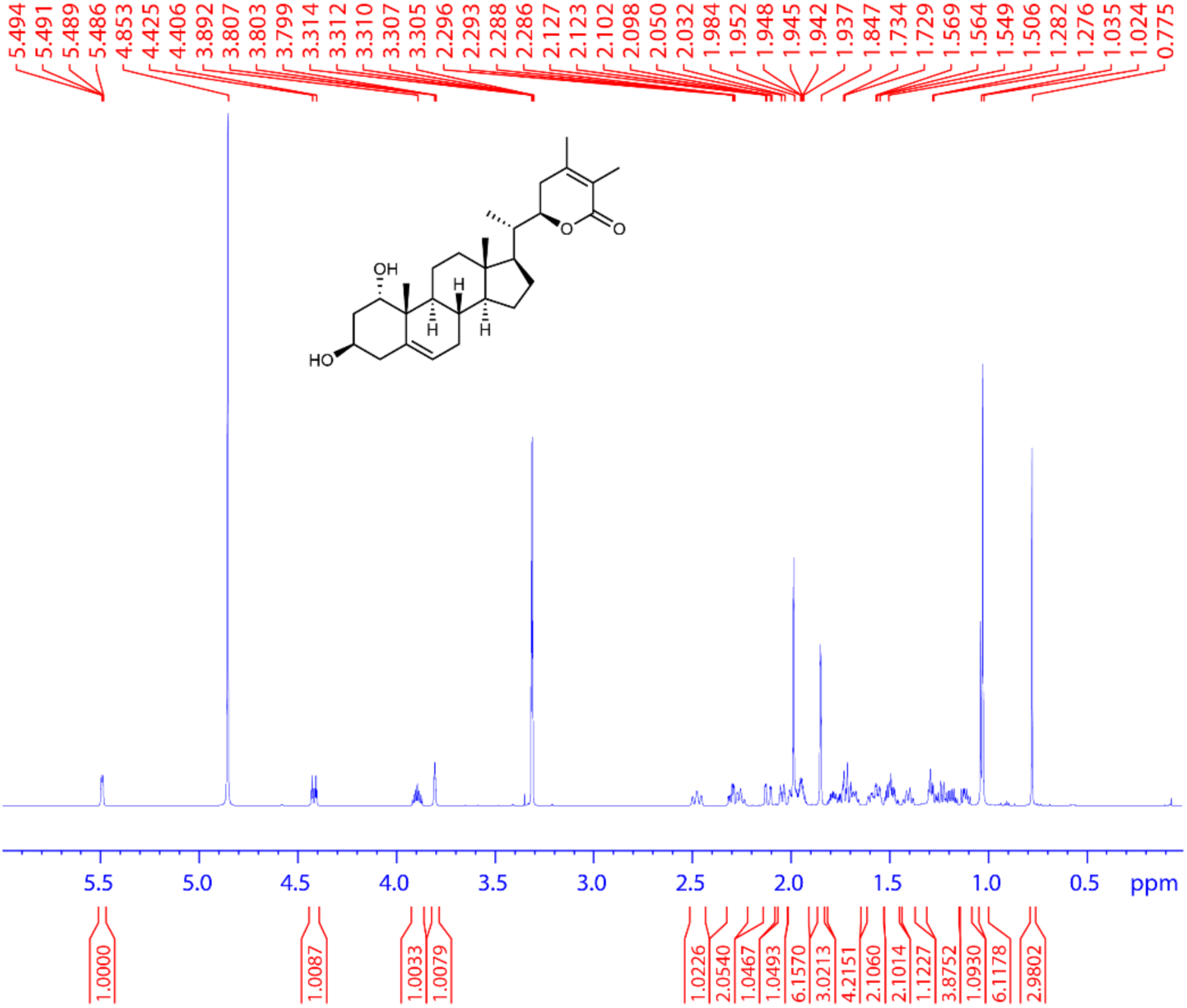
**^1^H spectrum of 7. (700 MHz, CD_3_OD, 298 K).**

**Supplementary Fig. 20.**
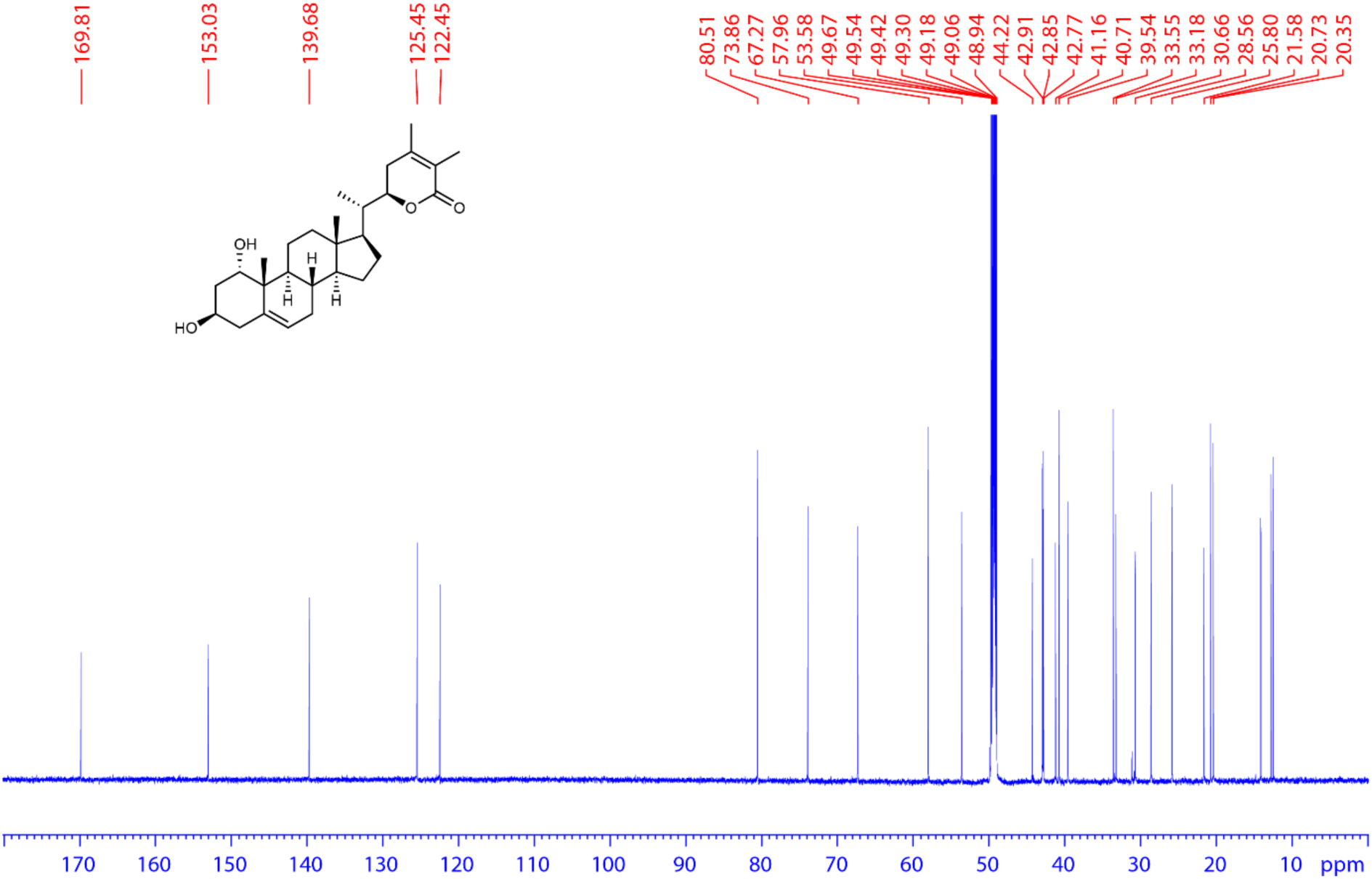
**^13^C spectrum of 7. (175 MHz, CD_3_OD, 298 K).**

**Supplementary Fig. 21.**
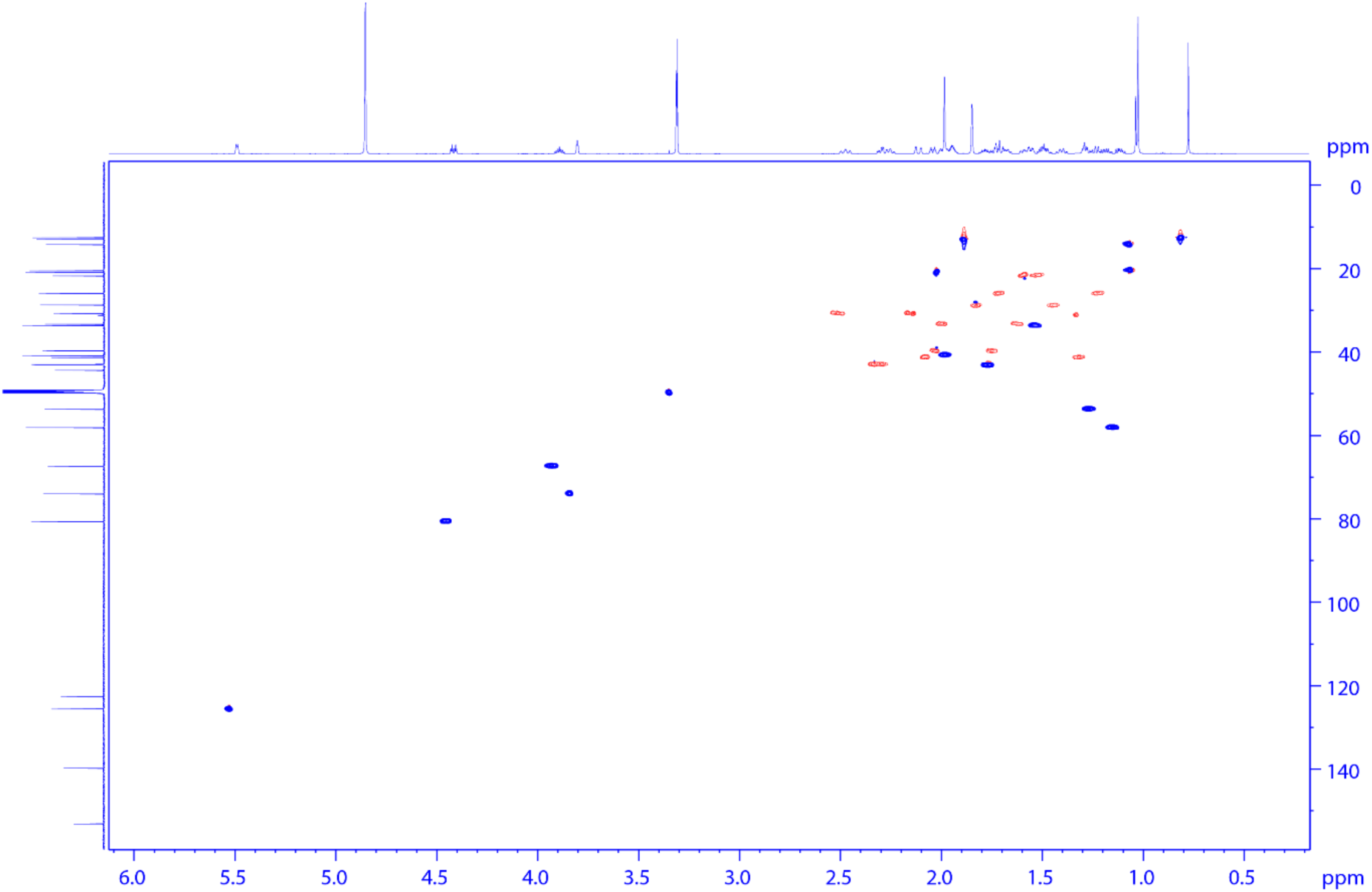
**HSQC spectrum of 7.**

**Supplementary Fig. 22.**
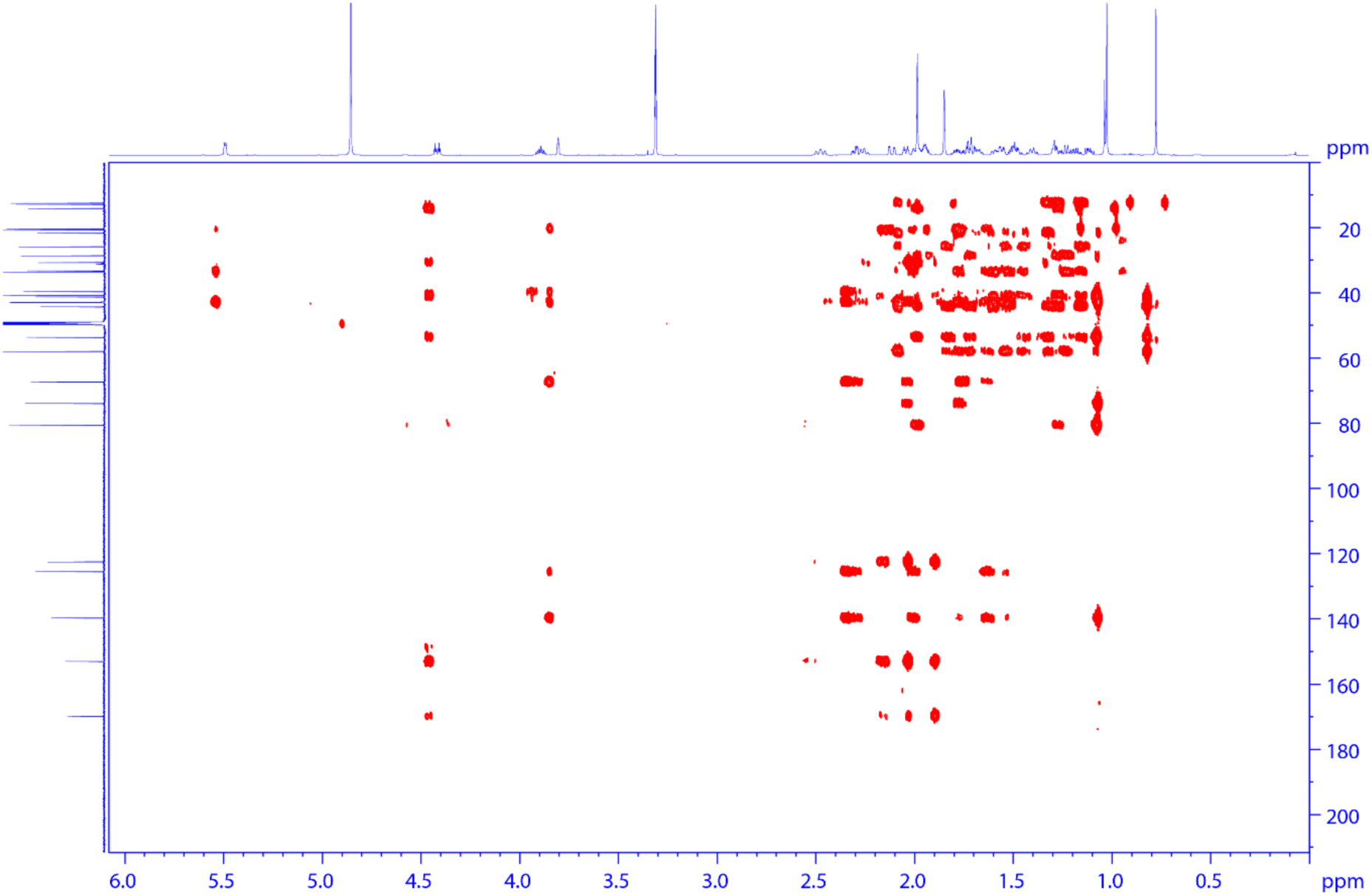
**HMBC spectrum of 7.**

**Supplementary Fig. 23.**
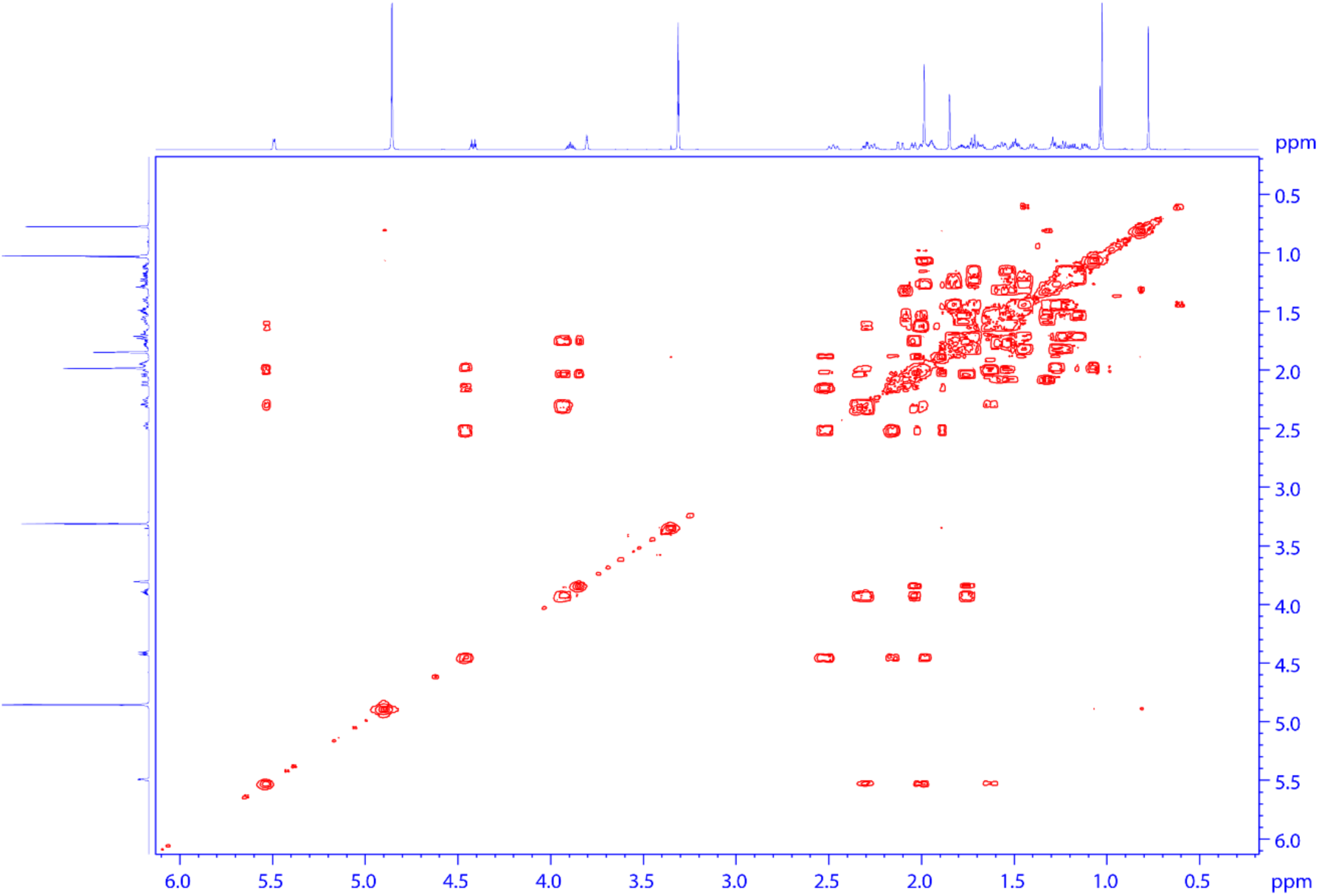
**HHCOSY spectrum of 7.**

**Supplementary Fig. 24.**
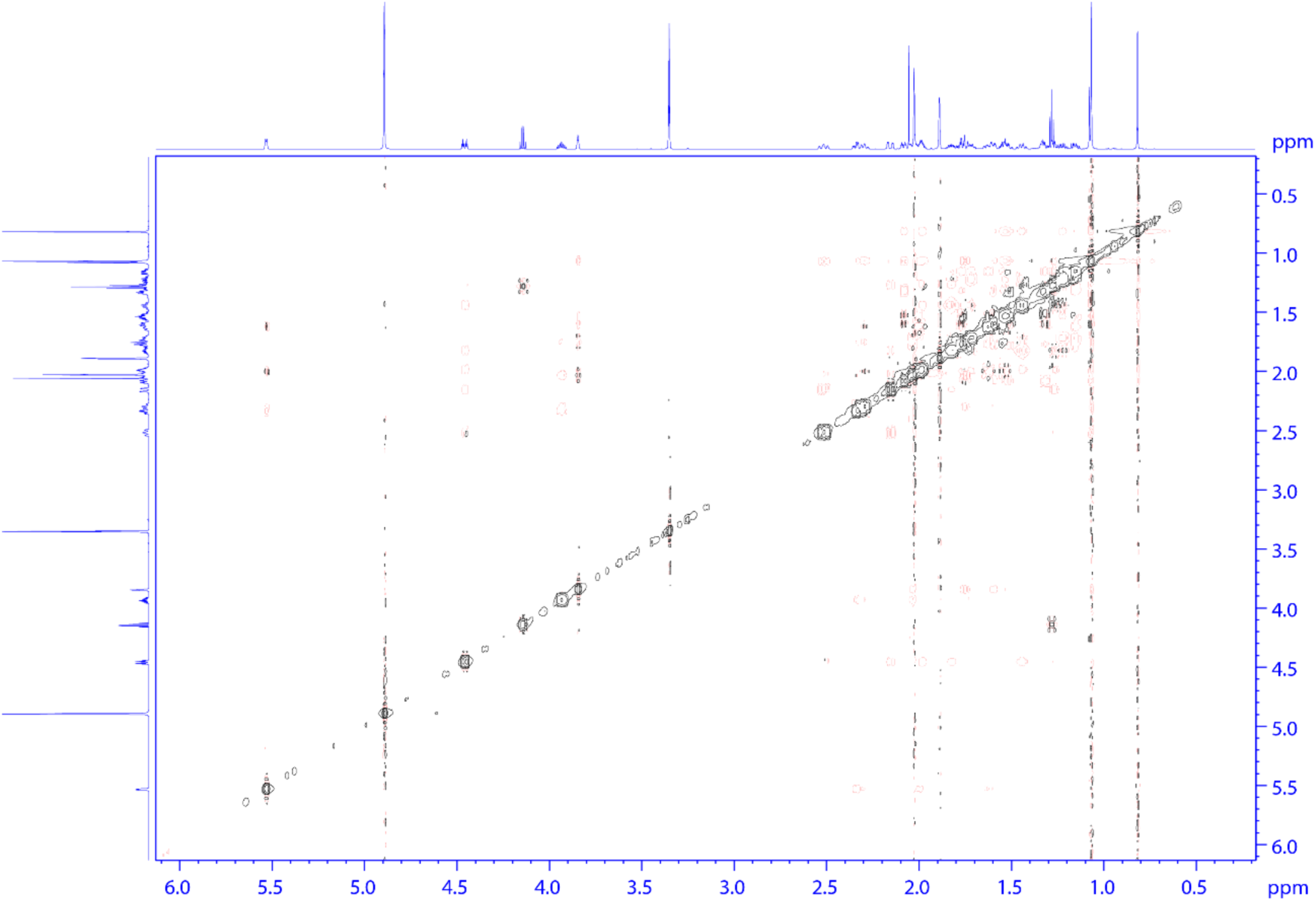
**NOESY spectrum of 7.**

**Supplementary Fig. 25.**
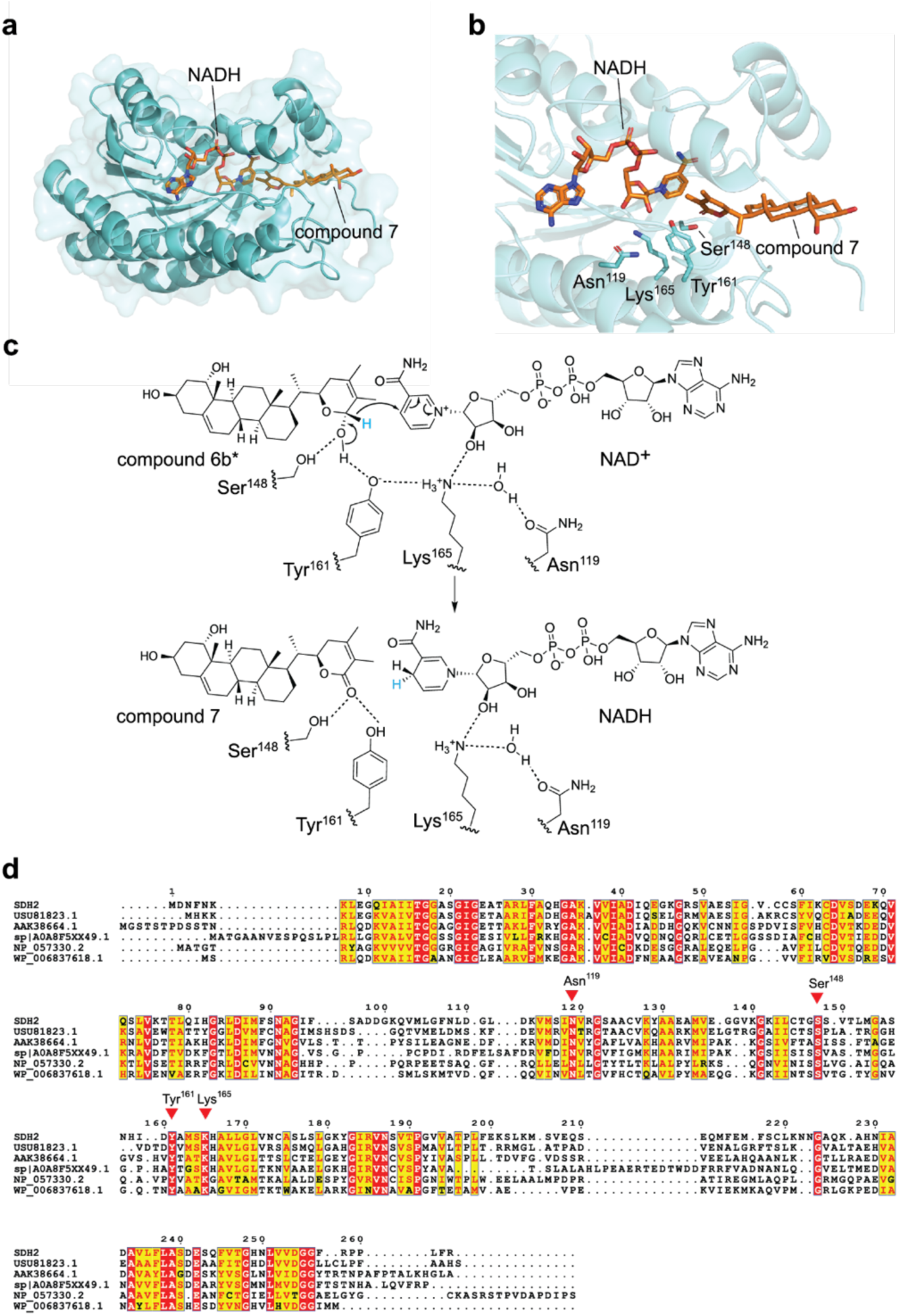
Computational modeling, ligand docking, and sequence analysis of SDH2. **a,** The overall structural model of SDH2 docked with **7** and the cofactor NADH. **b,** A close-up view of the SDH2 active site, showing the top docking pose of **7**, NADH, and several active-site residues (Asn^119^, Ser^148^, Tyr^161^, and Lys^165^) predicted to be involved in catalysis. **c**, The proposed catalytic mechanism for the oxidation of **6b\*** (the predicted favored stereoisomer of **6b** as a substrate for SDH2) to **7** by SDH2 using NAD^+^ as the electron acceptor. Based on the docking results, we predict that the hydride from **6b\*** (highlighted in blue) is transferred to the “pro-S” position on NAD⁺. **d**, Sequence alignment of SDH2 with other short-chain dehydrogenase homologs, with conserved active site residues indicated by the red triangles.

**Supplementary Fig. 26.**
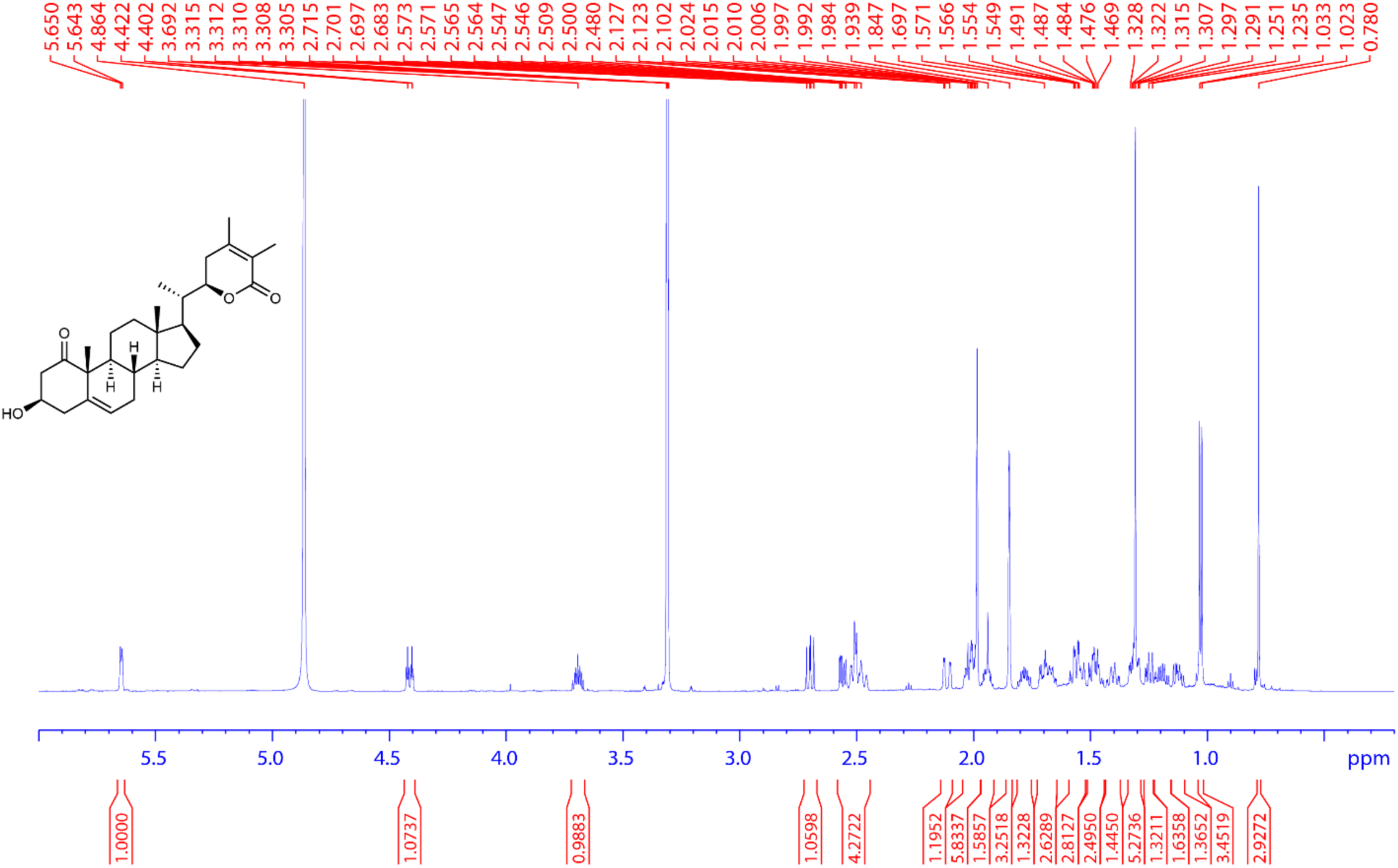
**^1^H spectrum of 8. (700 MHz, CD_3_OD, 298 K).**

**Supplementary Fig. 27.**
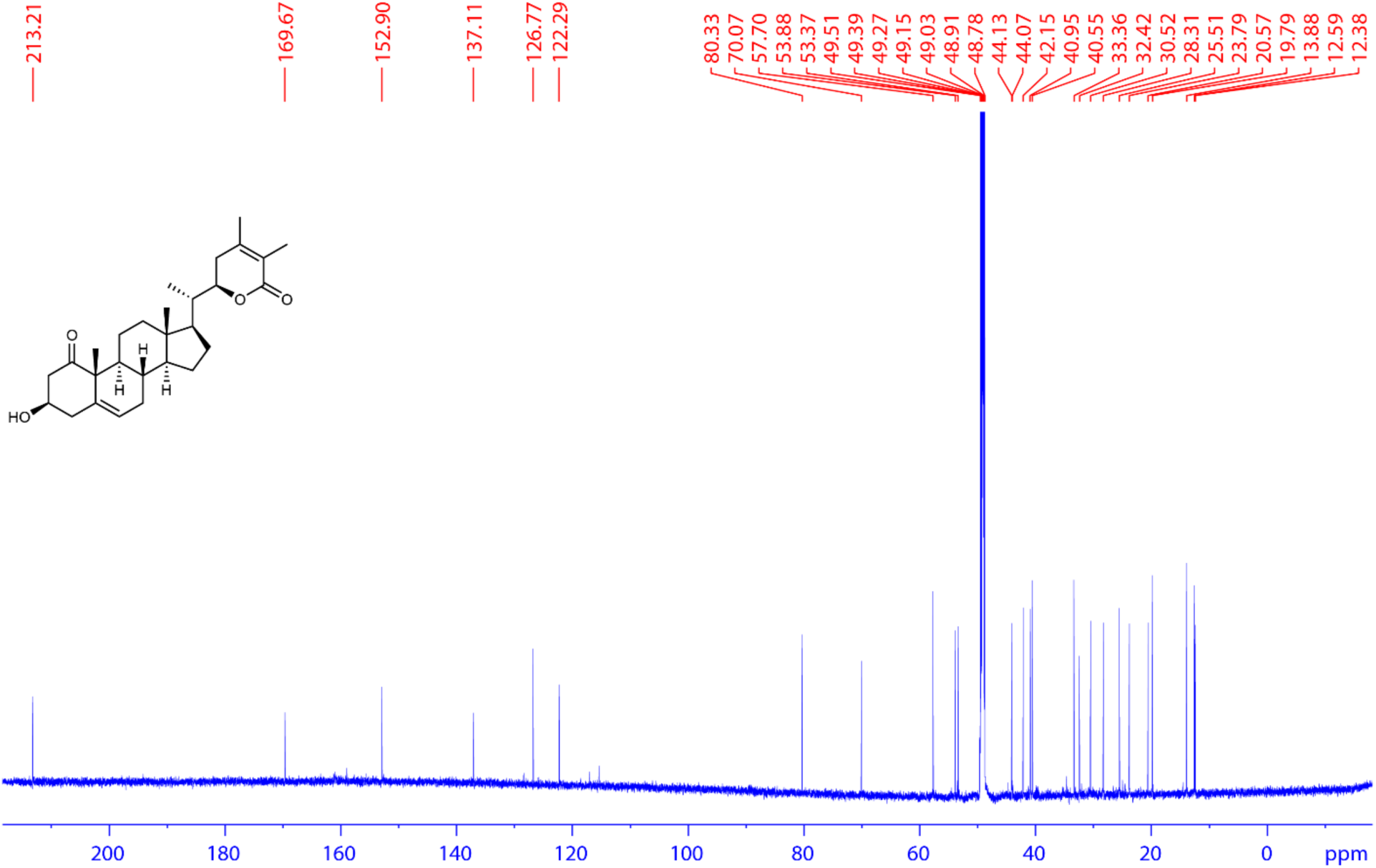
**^13^C spectrum of 8. (175 MHz, CD_3_OD, 298 K).**

**Supplementary Fig. 28.**
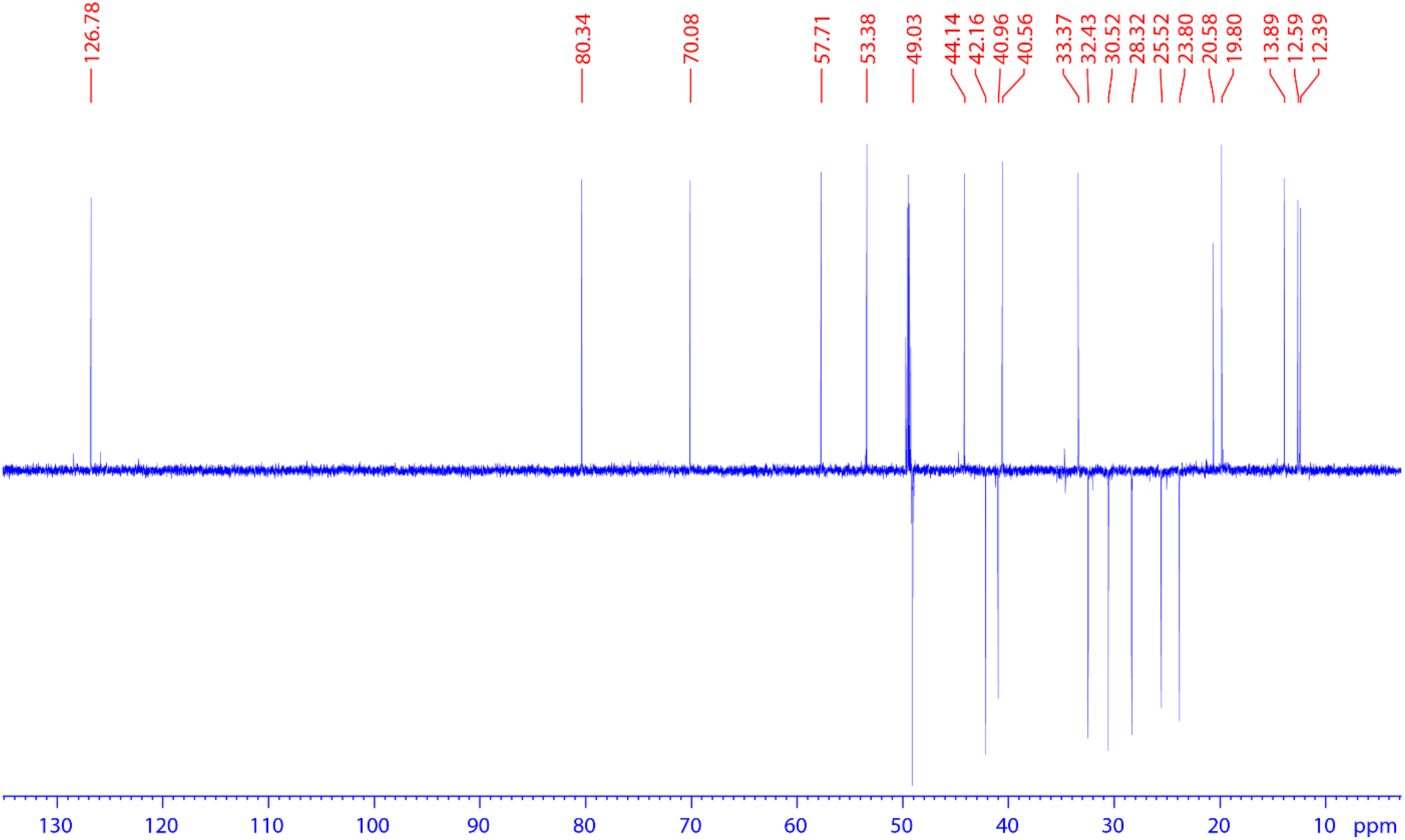
**DEPT spectrum of 8.**

**Supplementary Fig. 29.**
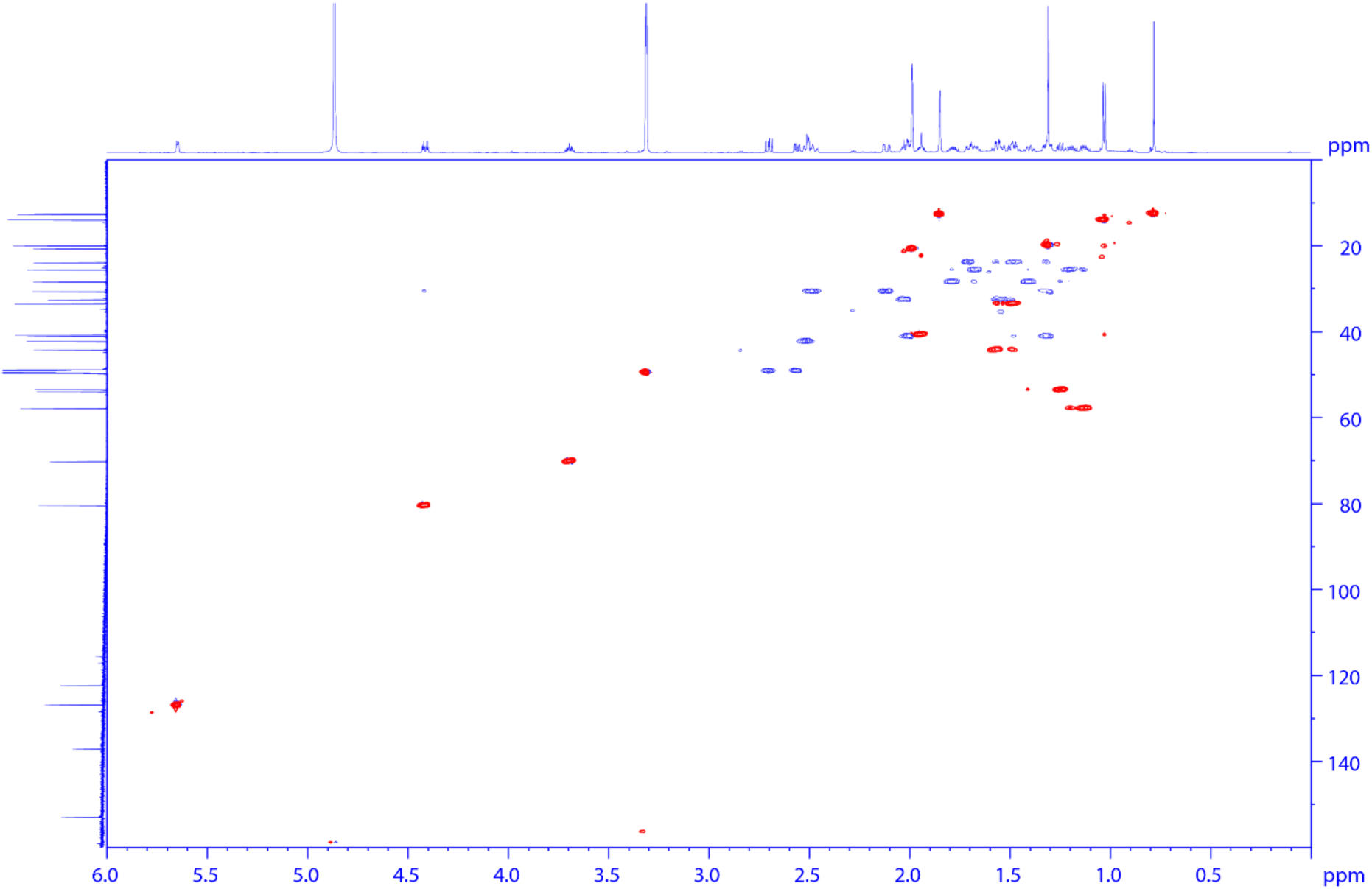
**HSQC spectrum of 8.**

**Supplementary Fig. 30.**
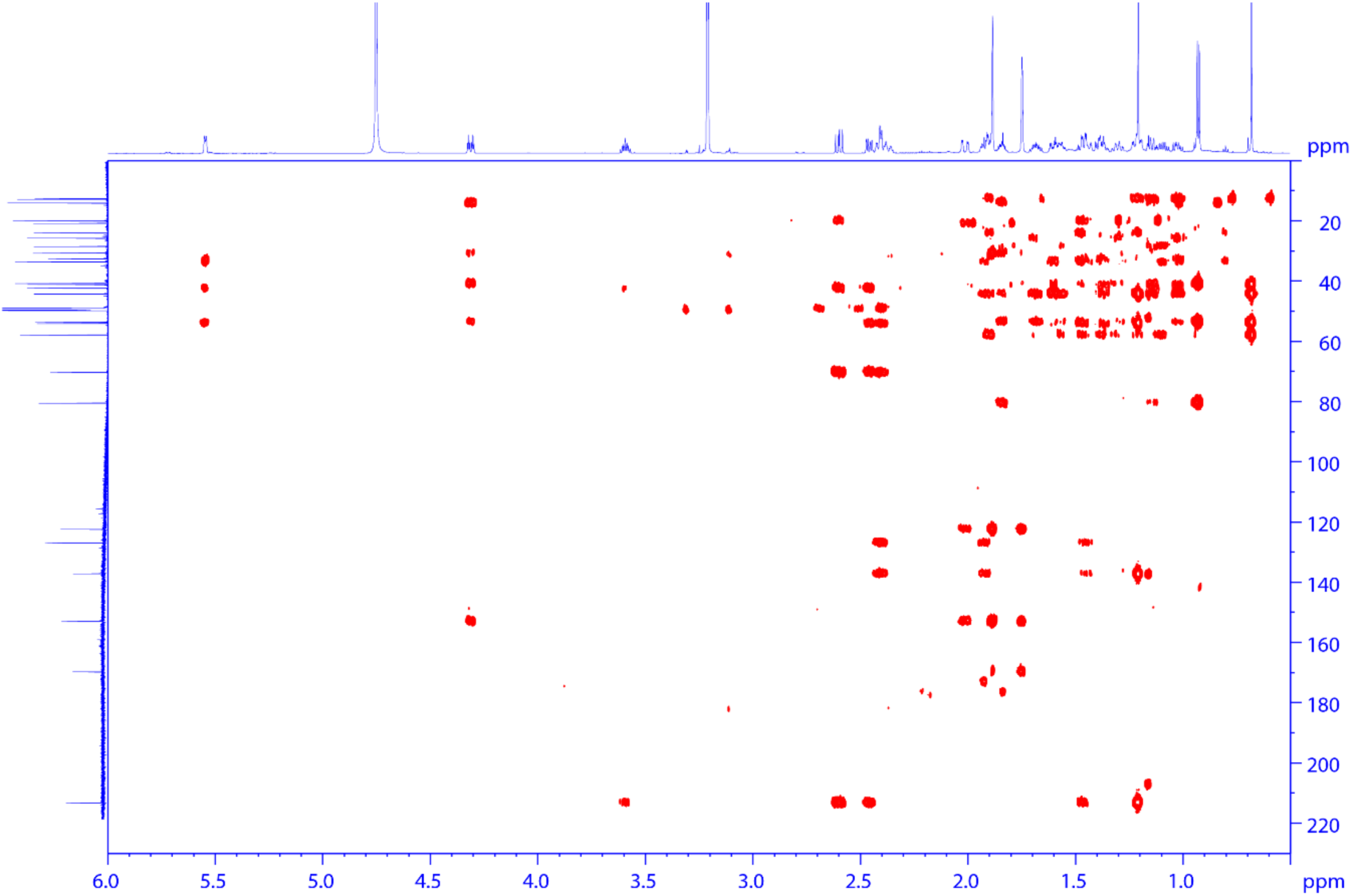
**HMBC spectrum of 8.**

**Supplementary Fig. 31.**
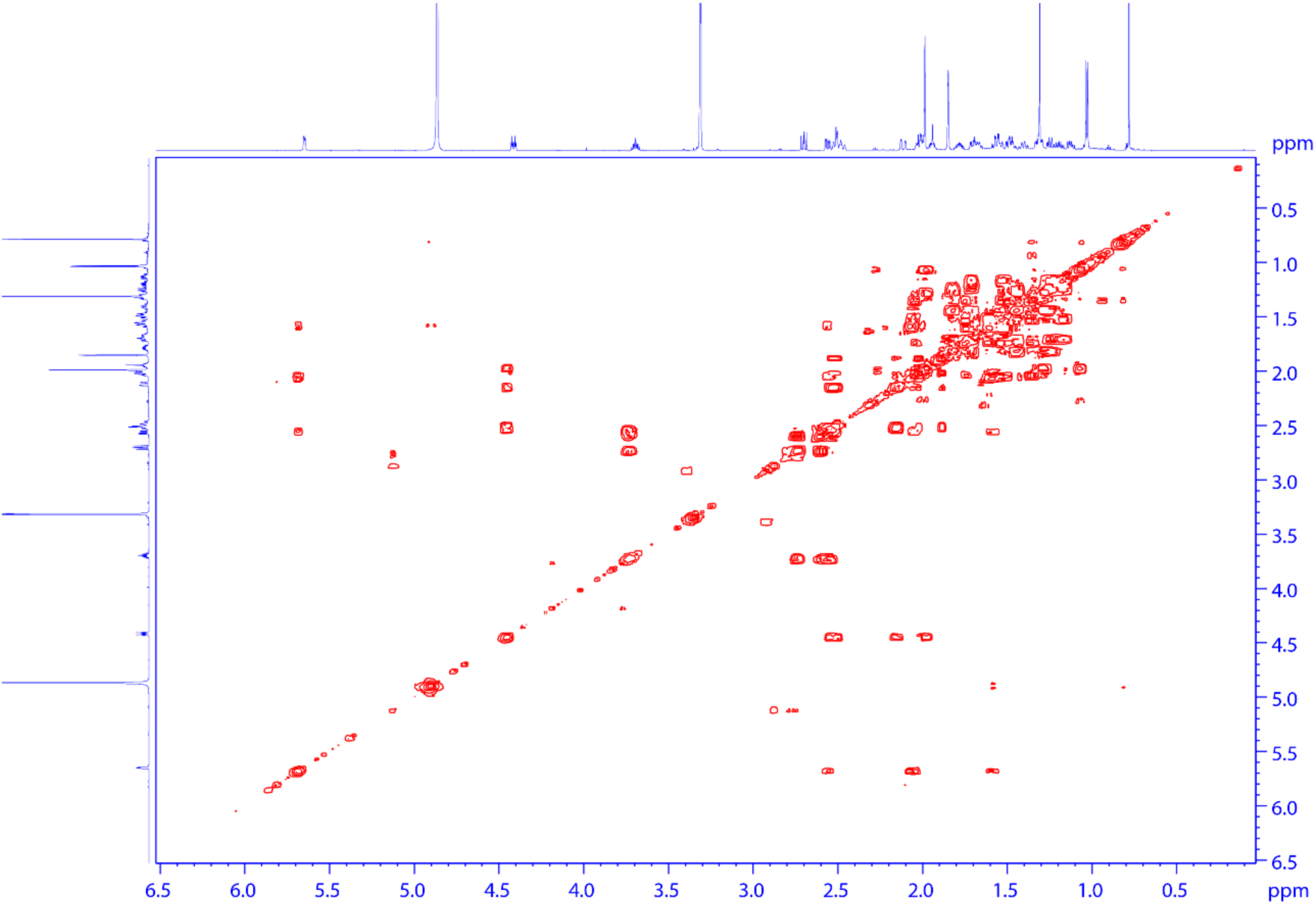
**HHCOSY spectrum of 8.**

**Supplementary Fig. 32.**
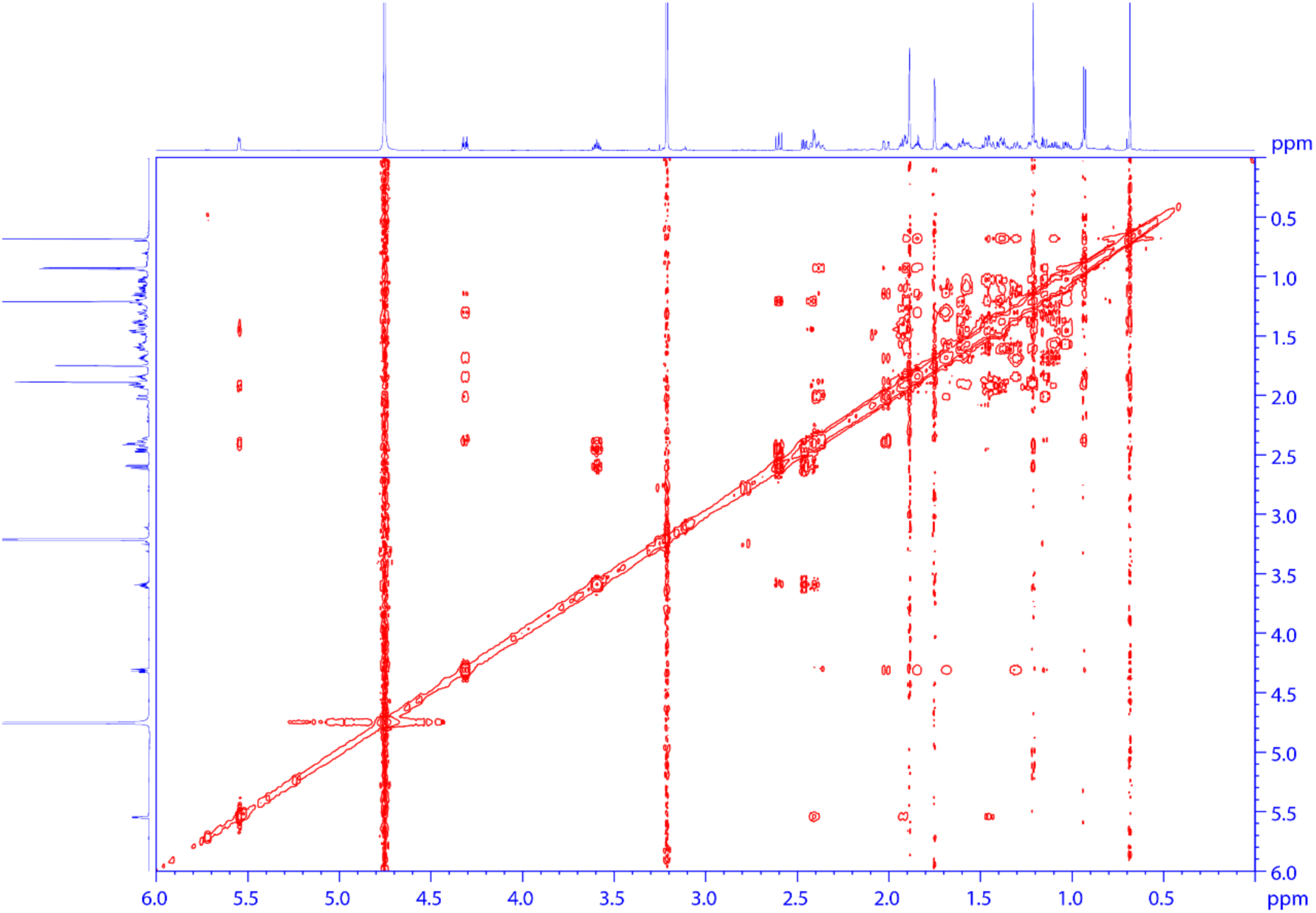
**NOESY spectrum of 8.**

**Supplementary Fig. 33.**
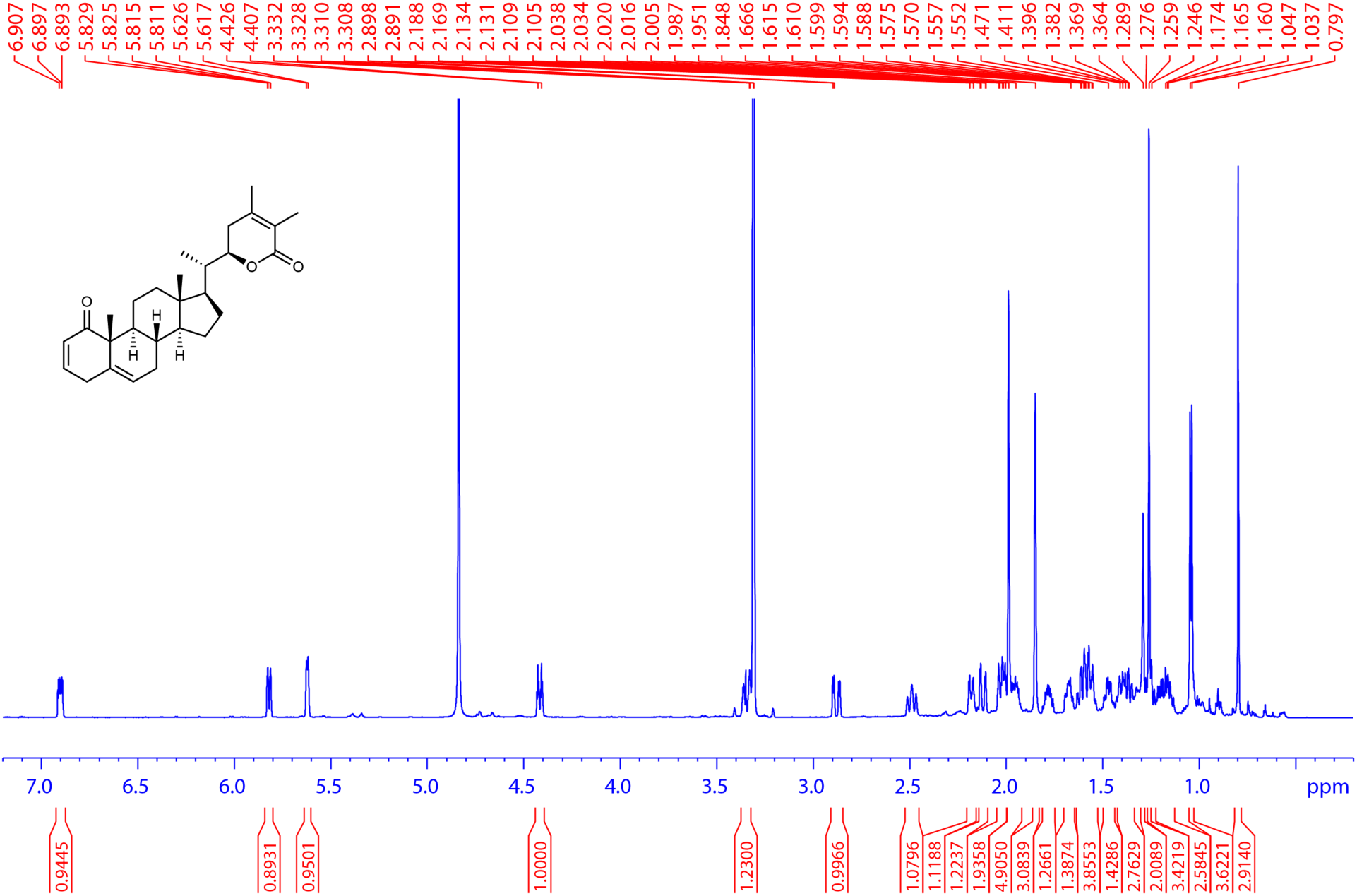
**^1^H spectrum of 10. (700 MHz, CD_3_OD, 298 K).**

**Supplementary Fig. 34.**
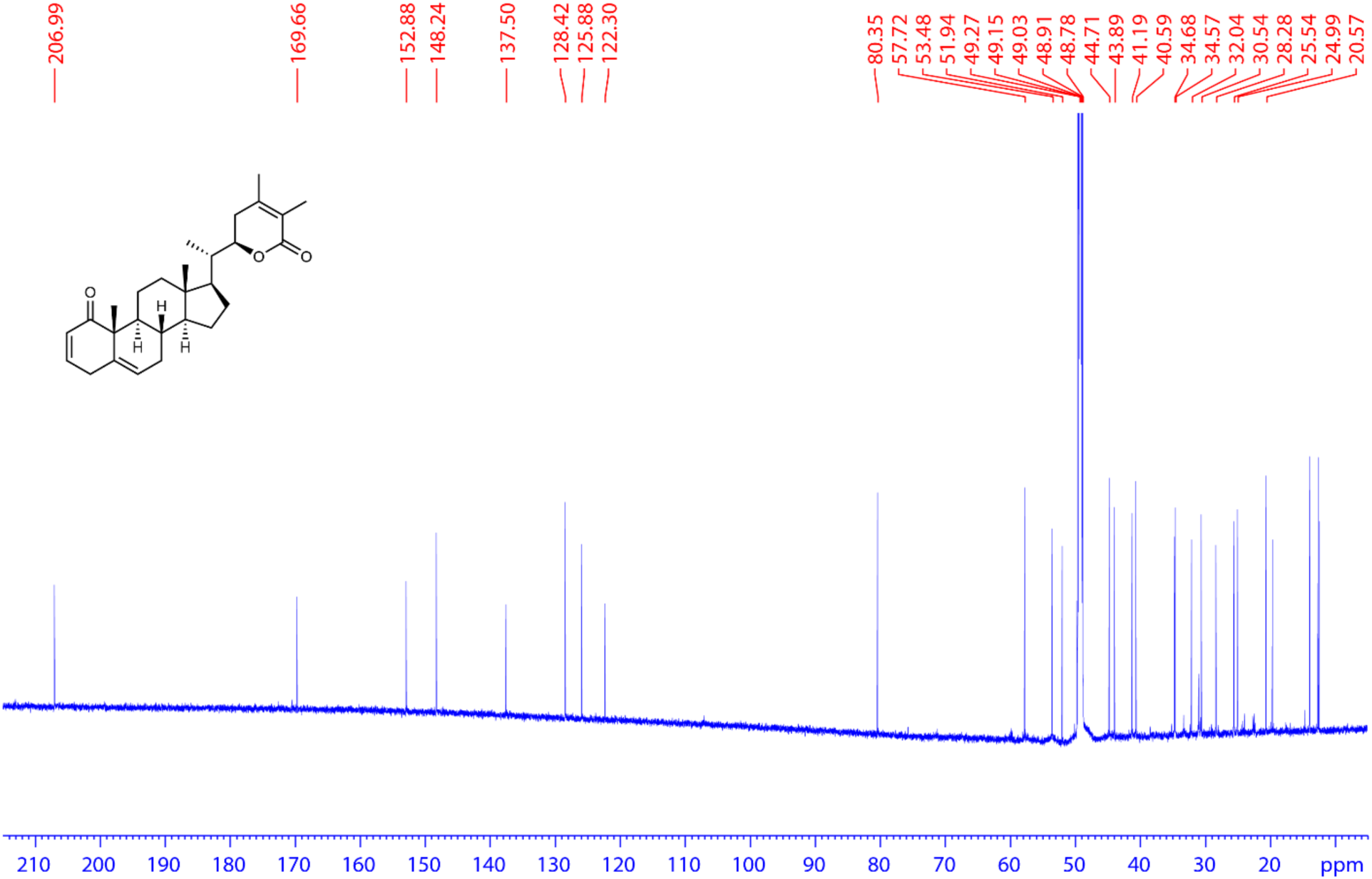
**^13^C spectrum of 10. (175 MHz, CD_3_OD, 298 K).**

**Supplementary Fig. 35.**
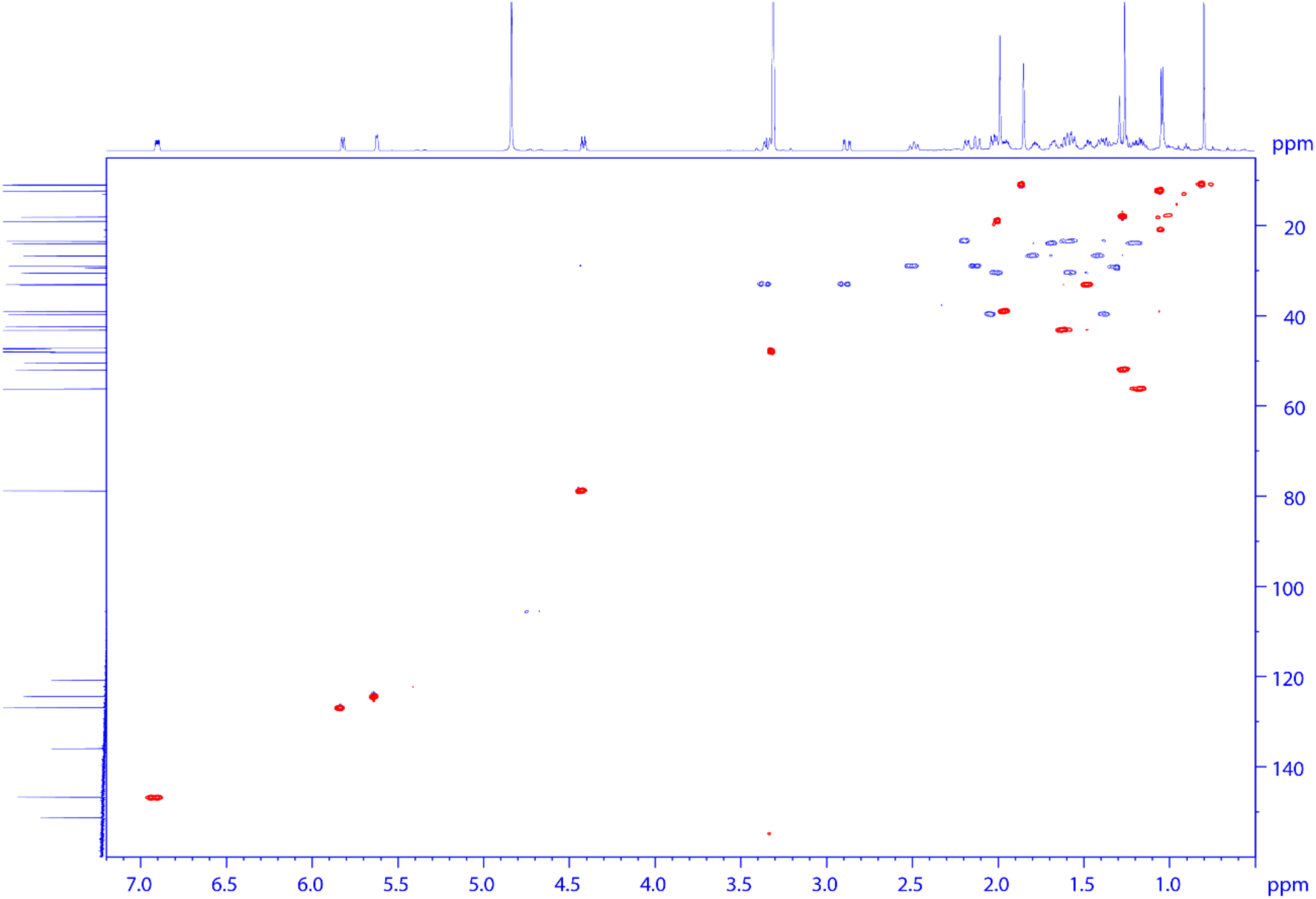
**HSQC spectrum of 10.**

**Supplementary Fig. 36.**
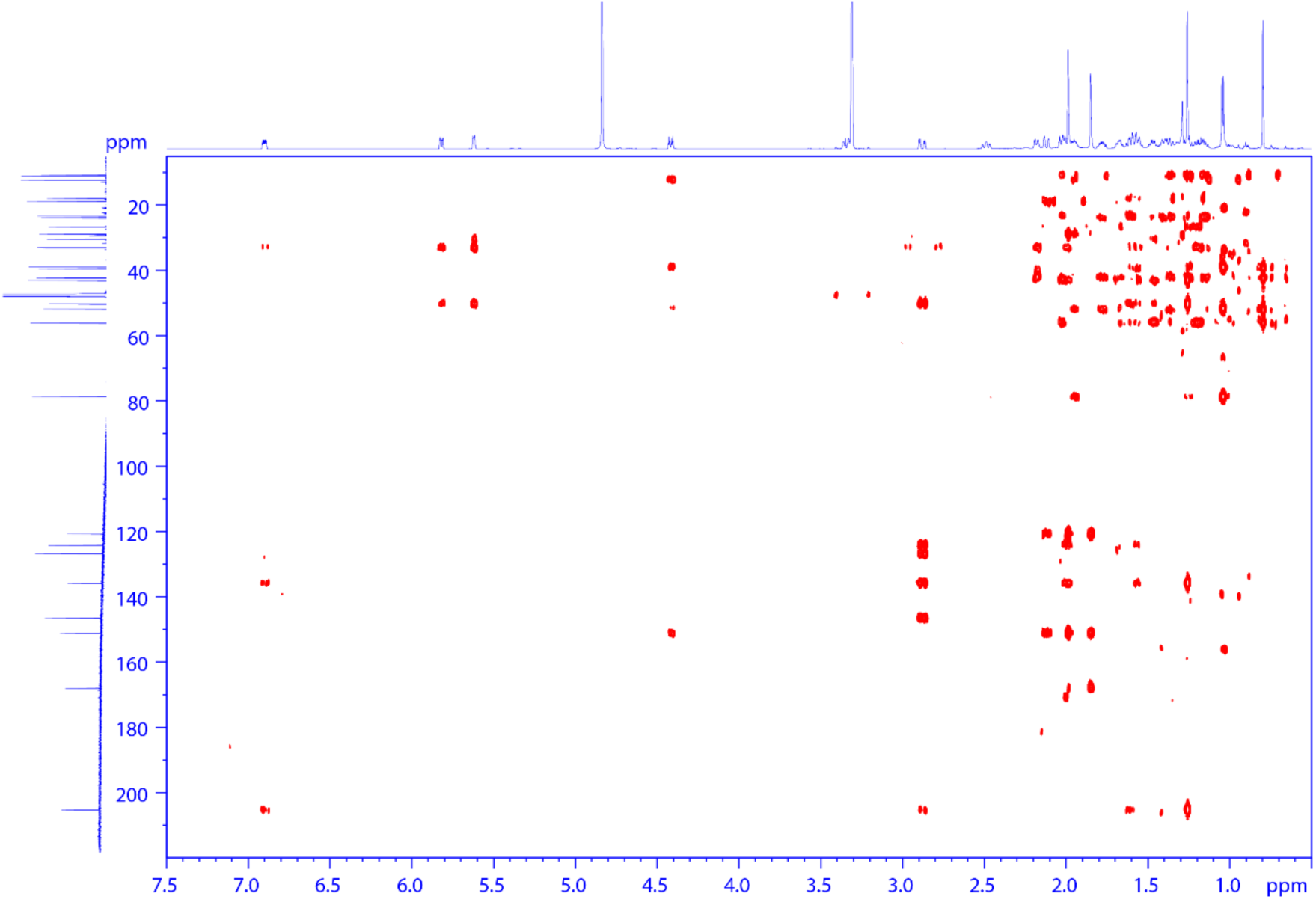
**HMBC spectrum of 10.**

**Supplementary Fig. 37.**
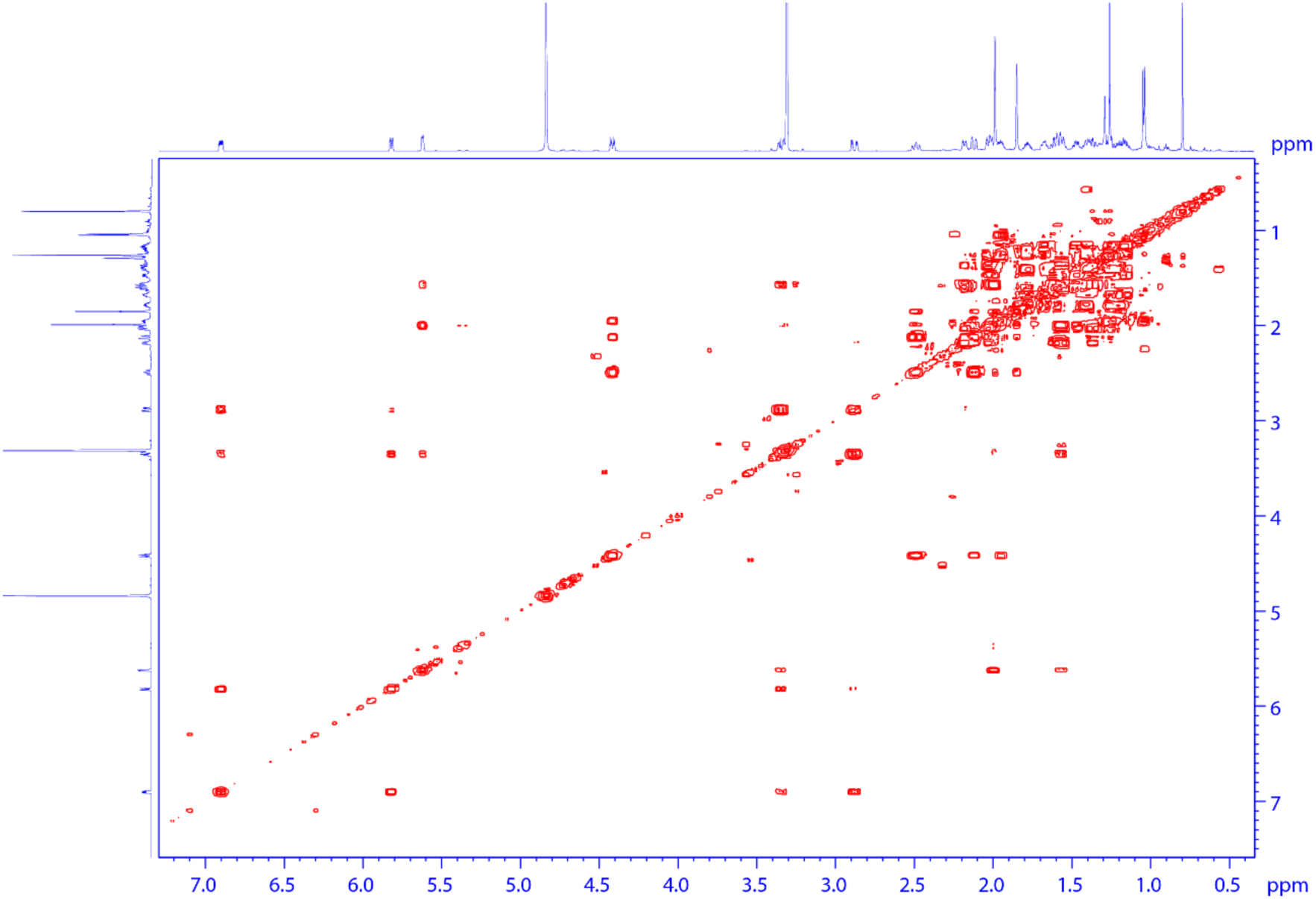
**HHCOSY spectrum of 10.**

**Supplementary Fig. 38.**
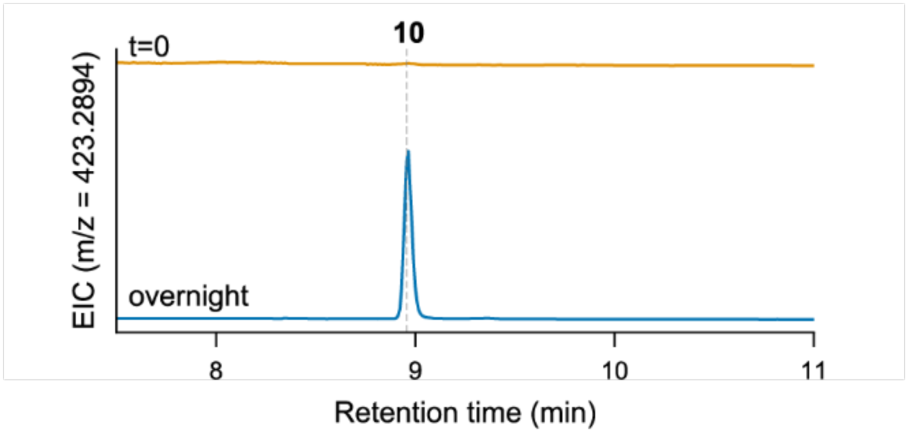
Spontaneous conversion of 10. EIC for [M+H]^+^ ion for **10** (*m/z* = 423.2894). Semi-pure **9** at timepoint zero (t=0) contained no detectable **10**. After incubation at 30 °C in potassium phosphate assay buffer overnight, production of **10** was observed. Representative traces are shown of n=3 replicates.

**Supplementary Fig. 39.**
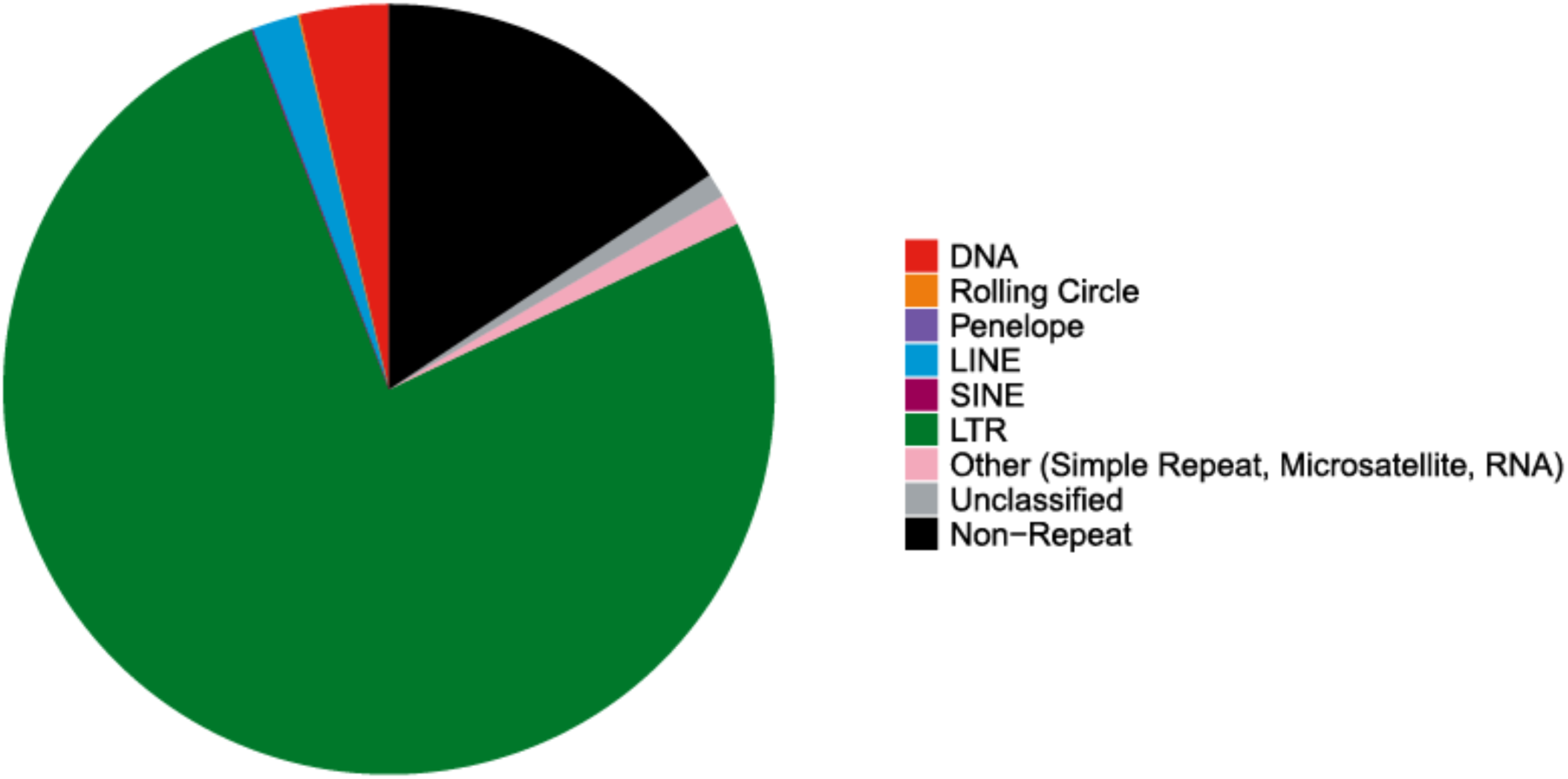
Transposable elements in the *W. somnifera* genome. Transposable element annotation was performed with Earl Grey. LINE (long interspersed nuclear element), SINE (short interspersed nuclear element), LTR (long terminal repeat). 81% of the *W. somnifera* genome consists of repetitive elements and 19% consists of non-repetitive elements.

**Supplementary Fig. 40.**
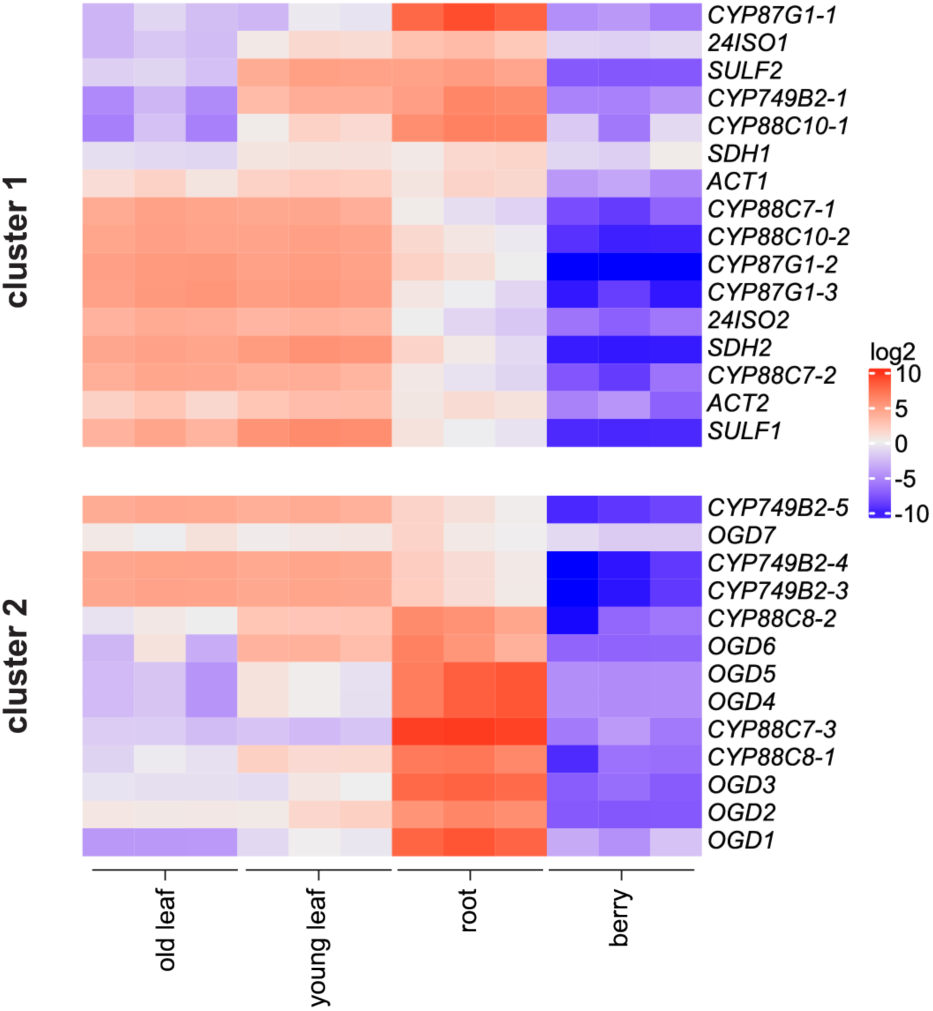
Unadjusted heatmap of RNA sequencing read counts. Heatmap displaying normalized log2-transformed RNA sequencing read counts for the biosynthetic cluster genes across four tissue types (old leaf, young leaf, root, and berry), n=3 biological replicates. Reads are centered around zero for each gene.

## Supplemental Tables

**Supplementary Table 1.** Summary of *W. somnifera* genome assembly and annotation.

|  | Number | Size (Mb) |
| --- | --- | --- |
| Total contigs | 532 | 2,792.40 |
| Contig N50 |  | 9 |
| Contig N75 |  | 5 |
| Total scaffolds | 419 | 2,792.68 |
| Scaffold N50 |  | 119.6 |
| Scaffold N75 |  | 105.1 |
| Total pseudochromosomes | 24 | 2,778.99 |

**Supplemental Table 2.** Comparison of amino acid sequence identity among leaf- and root-expressed paralogs in the biosynthetic clusters. Percent amino acid identity (%AA ID) between paralogs in the biosynthetic gene clusters (pseudogenes were excluded from the analysis). Whether the paralog was more highly expressed in leaf or root tissue is indicated. Yellow highlighted and bolded gene is the sequence selected for candidate gene testing. In some cases, the tested sequences differed slightly from the sequences obtained from the *W. somnifera* genome because the original sequences were obtained from an assembled transcriptome for *W. somnifera*. % AA ID is greater than 99% between transcriptome and genome versions of genes and the differences are most likely due to natural variation between individual plants or sequencing/assembly mistakes.

|  | 24ISO-1 (root) | 24ISO-2 (leaf) |
| --- | --- | --- |
| 24ISO-1 (root) | 100% | 93% |
| **24ISO-2 (leaf)** | 93% | 100% |

\*% AA ID of 24ISO-2 to tested version (from transcriptome): **100%**

|  | CYP87G1-1 (root) | CYP87G1-2 (leaf) | CYP87G1-3 (leaf) |
| --- | --- | --- | --- |
| CYP87G1-1 (root) | 100% | 95% | 94% |
| **CYP87G1-2 (leaf)** | 94% | 100% | 99% |
| CYP87G1-3 (leaf) | 95% | 99% | 100% |

\*% AA ID of CYP87G1-2 to tested version (from transcriptome): **99.4%**

|  | CYP88C7-1 (leaf) | CYP88C7-2 (leaf) | CYP88C7-3 (root) |
| --- | --- | --- | --- |
| **CYP88C7-1 (leaf)** | 100% | 99.8% | 90% |
| CYP88C7-2 (leaf) | 99.8% | 100% | 90% |
| CYP88C7-3 (root) | 90% | 90% | 100% |

\*% AA ID of CYP88C7-1 to tested version (from transcriptome): **100%**

|  | CYP749B2-1 (root) | CYP749B2-5 (leaf) | CYP749B2-4 (leaf) | CYP749B2-3 (leaf) |
| --- | --- | --- | --- | --- |
| CYP749B2-1 (root) | 100% | 88% | 89% | 89% |
| CYP749B2-5 (leaf) | 88% | 100% | 98% | 99% |
| **CYP749B2-4 (leaf)** | 89% | 98% | 100% | 99% |
| CYP749B2-3 (leaf) | 89% | 99% | 99% | 100% |

\*% AA ID of CYP749B2-3 to tested version (from transcriptome): **100%**

|  | CYP88C10-1 (root) | CYP88C10-2 (leaf) |
| --- | --- | --- |
| CYP88C10-1 (root) | 100% | 75% |
| **CYP88C10-2 (leaf)** | 75% | 100% |

\*% AA ID of CYP88C10-2 to tested version (from transcriptome): **99.6%**

|  |  |  |
| --- | --- | --- |
|  | SDH-1 (root) | SDH-2 (leaf) |
| SDH-1 (root) | 100% | 88% |
| **SDH-2 (leaf)** | 88% | 100% |

\*% AA ID of SDH-2 to tested version (amplified from *W. somnifera* cDNA): **100%**

|  |  |  |
| --- | --- | --- |
|  | SULF1 (leaf) | SULF2 (root) |
| **SULF1 (leaf)** | 100% | 88% |
| SULF2 (root) | 88% | 100% |

\*% AA ID of SULF1 to tested version (from transcriptome): **100%**

|  |  |  |
| --- | --- | --- |
|  | CYP88C8-2 (root) | CYP88C8-1 (root) |
| **CYP88C8-2 (root)** | 100% | 99% |
| CYP88C8-1 (root) | 99% | 100% |

\*% AA ID of CYP88C8-2 to tested version (from transcriptome): **99.8%**

**Supplemental Table 3.** Yeast strains used in this work.

| Strain | Genotype | Source |
| --- | --- | --- |
| Y4108 | *MATα his3Δ1 leu2Δ0 lys2Δ0 ura3Δ0 erg4::KanMX4 erg5::ble<sup>r</sup>* | 31 |
| Y6798 | Y4108 + ura3:: <T<sub>ADH1</sub>-ATR1-P<sub>GAL1/10</sub>-CYP87G1-T<sub>PGK1</sub>> | This work |
| Y6852 | Y6798 + leu2:: <T<sub>ENO2</sub>-7RED-P<sub>GAL1/10</sub>-24ISO-1-T<sub>ADH1</sub>> | This work |
| Y7037 | Y6852 + pcfB2312-Hyg (cas9 plasmid) | This work |
| Y7106 | Y7037 + XII-I:: <P<sub>GAL1</sub>-CYP88C7-1-T<sub>CYC1</sub>> | This work |
| Y7126 | Y4108 + leu2:: <T<sub>ENO2</sub>-7RED-P<sub>GAL1/10</sub>-24ISO-2-T<sub>ADH1</sub>> | This work |
| Y7162 | Y7037 + XII-4:: <T<sub>ADH1</sub>-CYP88C7-P<sub>GAL1/10</sub>-CYP749B2-T<sub>CYC1</sub>> | This work |
| Y7193 | Y4108 + X-3:: <P<sub>GAL1</sub>-ATR1-T<sub>CYC1</sub>> | This work |
| Y7215 | Y4108 + XII-4:: <P<sub>GAL1</sub>-ATR1-T<sub>CYC1</sub>> + X-3:: <P<sub>GAL1</sub>-CYP88C10-T<sub>CYC1</sub>> | This work |
| Y7509 | Y7162 + XI-2:: <P<sub>GAL1</sub>-SDH1-T<sub>CYC1</sub>> | This work |
| Y7514 | Y7126 + ura3:: <T<sub>ADH1</sub>-ATR1-P<sub>GAL1/10</sub>-CYP87G1-1-T<sub>PGK1</sub>> | This work |
| Y7515 | Y7126 + ura3:: <T<sub>ADH1</sub>-ATR1-P<sub>GAL1/10</sub>-CYP87G1-2-T<sub>PGK1</sub>> | This work |
| Y7516 | Y7126 + ura3:: <T<sub>ADH1</sub>-ATR1-P<sub>GAL1/10</sub>-CYP87G1-3-T<sub>PGK1</sub>> | This work |
| Y7517 | Y4108 + leu2:: <T<sub>ENO2</sub>-7RED-P<sub>GAL1/10</sub>-24ISO-1-T<sub>ADH1</sub>> | This work |
| Y7541 | Y7037 + X-3:: <P<sub>GAL1</sub>-CYP88C7-2-T<sub>CYC1</sub>> | This work |
| Y7542 | Y7037 + X-3:: <P<sub>GAL1</sub>-CYP88C7-3-T<sub>CYC1</sub>> | This work |
| Y7543 | Y7037 + X-3:: <P<sub>GAL1</sub>-CYP88C7-1-T<sub>CYC1</sub>> | This work |
| Y7567 | Y7037 + his3:: <P<sub>GAL1</sub>-CYP87G1-T<sub>CYC1</sub>> + XII-4:: <T<sub>ADH1</sub>-CYP88C7-P<sub>GAL1/10</sub>-CYP749B2-T<sub>CYC1</sub>> + X-3:: <T<sub>ADH1</sub>-CYP88C7-P<sub>GAL1/10</sub>-CYP749B2-T<sub>CYC1</sub>> | This work |
| Y7587 | Y7567 + XI-2:: <P<sub>GAL1</sub>-SDH1-T<sub>CYC1</sub>> | This work |
| Y7610 | Y7162 + XI-2:: <P<sub>GAL1</sub>-SDH2-T<sub>CYC1</sub>> | This work |
| Y7658 | Y7610 + XI-3:: <P<sub>GAL1</sub>-CYP88C10-T<sub>CYC1</sub>> | This work |
| Y7666 | Y7509 + XI-5:: <P<sub>GAL1</sub>-SULF1-T<sub>CYC1</sub>> | This work |
| Y7665 | Y7509 + XI-5:: <P<sub>GAL1</sub>-SULF2-T<sub>CYC1</sub>> | This work |
| Y7748 | Y7610 + XI-5:: <P<sub>GAL1</sub>-SULF1-T<sub>CYC1</sub>> | This work |
| Y7752 | Y7658 + XI-5:: <P<sub>GAL1</sub>-SULF1-T<sub>CYC1</sub>> | This work |
| Y7771 | Y7567 + XI-2:: <T<sub>ADH1</sub>-SDH2-P<sub>GAL1/10</sub>-SDH2-T<sub>CYC1</sub>> + XII-5:: <T<sub>ADH1</sub>-CYP88C10-P<sub>GAL1/10</sub>-CYP88C10-T<sub>CYC1</sub>> | This work |
| Y7841 | Y7771 + XI-5:: <T<sub>ADH1</sub>-SULF1-P<sub>GAL1/10</sub>-SULF2-T<sub>CYC1</sub>> | This work |

**Supplementary Table 4.** ^13^C NMR data of isolated compounds (175 MHz, 298 K). ^13^C-NMR (175MHz) Data for Compound **3, 4, 7, 8 and 10** (ppm)

| Position | Cmpd3<sup>a</sup> | Cmpd4<sup>a</sup> | Cmpd7<sup>b</sup> | Cmpd8<sup>b</sup> | Cmpd10<sup>b</sup> |
| --- | --- | --- | --- | --- | --- |
| 1 | 37.4 | 73.2 | 73.9 | 213.2 | 207.0 |
| 2 | 31.8 | 38.0 | 39.5 | 49.0 | 128.4 |
| 3 | 72.0 | 66.9 | 67.3 | 70.1 | 148.2 |
| 4 | 42.5 | 41.2 | 42.8 | 42.1 | 34.6 |
| 5 | 141.0 | 136.9 | 139.7 | 137.1 | 137.5 |
| 6 | 121.8 | 126.0 | 125.4 | 126.8 | 125.9 |
| 7 | 32.1 | 31.8 | 33.2 | 32.4 | 32.0 |
| 8 | 32.1 | 31.9 | 33.6 | 33.4 | 34.7 |
| 9 | 50.3 | 41.7 | 42.9 | 44.1 | 44.7 |
| 10 | 36.7 | 41.7 | 42.7 | 53.9 | 51.9 |
| 11 | 21.2 | 24.5 | 25.8 | 25.5 | 25.5 |
| 12 | 39.9 | 39.5 | 41.2 | 40.9 | 41.2 |
| 13 | 42.9 | 42.7 | 44.2 | 44.1 | 43.9 |
| 14 | 56.6 | 56.3 | 58.0 | 57.7 | 57.7 |
| 15 | 24.6 | 20.3 | 21.6 | 23.8 | 25.0 |
| 16 | 27.6 | 27.4 | 28.6 | 28.3 | 28.3 |
| 17 | 53.2 | 52.9 | 53.6 | 53.4 | 53.5 |
| 18 | 12.0 | 11.8 | 12.4 | 12.4 | 12.4 |
| 19 | 19.5 | 19.5 | 20.3 | 19.8 | 19.6 |
| 20 | 40.7 | 40.3 | 40.7 | 40.6 | 40.6 |
| 21 | 12.6 | 12.5 | 14.1 | 13.9 | 13.9 |
| 22 | 70.8 | 71.2 | 80.5 | 80.3 | 80.4 |
| 23 | 34.6 | 34.2 | 30.7 | 30.5 | 30.5 |
| 24 | 128.6 | 129.1 | 153.0 | 152.9 | 152.9 |
| 25 | 124.6 | 124.0 | 122.4 | 122.3 | 122.3 |
| 26 | 20.8 | 20.6 | 12.7 | 12.6 | 12.6 |
| 27 | 21.1 | 21.0 | 169.8 | 169.7 | 169.7 |
| 28 | 18.5 | 18.3 | 20.7 | 20.6 | 20.6 |

a: $\text{CDCl}_3$ , b: $\text{CD}_3\text{OD}$

**Supplementary Table 5.** 1H NMR data of isolated compounds (700 MHz, 298 K) ^1^H-NMR (700MHz) Data for Compound **3, 4, 7, 8 and 10** (ppm)

| Position | Cmpd3<sup>a</sup> | Cmpd4<sup>a</sup> | Cmpd7<sup>b</sup> | Cmpd8<sup>b</sup> | Cmpd10<sup>b</sup> |
| --- | --- | --- | --- | --- | --- |
| 1 | 1.08, 1.85 (2H, m) | 3.89 (1H, t, *J* = 2.5) | 3.80(1H, t, *J* = 2.7) |  |  |
| 2 | 1.52, 1.84 (2H, m) | 1.77, 2.12 (2H, m) | 1.71, 1.99 (2H, m) | 2.56 (1H, dd, *J* = 12.5, 9.7), 2.70<br>(1H, ddd, *J* = 12.5, 5.6, 1.4), | 5.82 (1H, dd, *J* = 10.0, 2.7) |
| 3 | 3.52 (1H, m) | 4.02 (1H, m) | 3.89 (1H, m) | 3.69 (1H, m) | 6.90 (1H, m) |
| 4 | 2.25, 2.30 (2H, m) | 2.33 (1H, t, *J* = 11.4),<br>2.40 (1H, dd, *J* = 3.4,<br>13.4) | 2.28 (2H, m) | 2.51 (2H, m) | 2.88 (1H, dd, *J* = 21.5, 4.8),<br>3.35 (1H, m) |
| 5 |  |  |  |  |  |
| 6 | 5.35 (1H, t, *J* = 3.2) | 5.61(1H, d, *J* = 5.3) | 5.49 (1H, d, *J* = 5.5) | 5.65 (1H, d, *J* = 5.5) | 5.62 (1H, d, *J* = 6.0) |
| 7 | 1.55, 1.99 (2H, m) | 1.60, 2.01 (2H, m) | 1.58, 1.96 (2H, m) | 1.54, 2.02 (2H, m) | 1.56, 2.00 (2H, m) |
| 8 | 1.47 (1H, m) | 1.50 (1H, m) | 1.50 (1H, m) | 1.52 (1H, m) | 1.47 (1H, m) |
| 9 | 0.94 (1H, m) | 1.57 (1H, m) | 1.73 (1H, m) | 1.57 (1H, m) | 1.61 (1H, m) |
| 10 |  |  |  |  |  |
| 11 | 1.50 (1H, m) | 1.14, 1.64 (2H, m) | 1.18, 1.68 (2H, m) | 1.19, 1.67 (2H, m) | 1.19, 1.67 (2H, m) |
| 12 | 1.18, 2.03 (2H, m) | 1.23, 2.04 (2H, m) | 1.28, 2.04 (2H, m) | 1.32, 2.00 (2H, m) | 1.37, 2.03 (2H, m) |
| 13 |  |  |  |  |  |
| 14 | 1.00 (1H, m) | 1.06 (1H, m) | 1.11 (1H, m) | 1.12 (1H, m) | 1.16 (1H, m) |
| 15 | 1.12, 1.62 (2H, m) | 1.49 (2H, m) | 1.49, 1.55 (2H, m) | 1.48, 1.70 (2H, m) | 1.57, 2.18 (2H, m) |
| 16 | 1.40, 1.78 (2H, m) | 1.42, 1.78 (2H, m) | 1.40, 1.79 (2H, m) | 1.40, 1.78 (2H, m) | 1.41, 1.79 (2H, m) |
| 17 | 1.15 (1H, m) | 1.16 (1H, m) | 1.23 (1H, m) | 1.25 (1H, m) | 1.26 (1H, m) |
| 18 | 0.72 (3H, s) | 0.72 (3H, s) | 0.77 (3H, s) | 0.78 (3H, s) | 0.80 (3H, s) |
| 19 | 1.01 (3H, s) | 1.04 (3H, s) | 1.02 (3H, s) | 1.31 (3H, s) | 1.26 (3H, s) |
| 20 | 1.85 (1H, m) | 1.87 (1H, m) | 1.94 (1H, m) | 1.94 (1H, m) | 1.95 (1H, m) |
| 21 | 0.99 (3H, d, *J* = 7.9) | 0.99 (3H, d, *J* = 6.7) | 1.03 (3H, d, *J* = 6.7) | 1.03 (3H, d, *J* = 6.7) | 1.04 (3H, d, *J* = 6.7) |
| 22 | 3.76 (1H, dt, *J* = 13.3, | 3.83 (1H, dt, *J* = 2.5, | 4.42 (1H, dt, *J* = 3.6, | 4.41 (1H, dt, *J* = 13.4, 3.6) | 4.42 (1H, dt, *J* = 13.4, 3.5) |

|  | 3.2) | 11.2) | 13.4) |  |  |
| --- | --- | --- | --- | --- | --- |
| 23 | 1.79 (1H, m), 2.40 (1H, dd, $J = 13.1, 15.6$) | 1.80, 2.44 (2H, m) | 2.11, 2.47 (2H, m) | 2.12, 2.48 (2H, m) | 2.12 (1H, dd, $J = 18.0, 2.5$),<br>2.49 (1H, t, $J = 16.2$) |
| 24 |  |  |  |  |  |
| 25 |  |  |  |  |  |
| 26 | 1.71 (3H, s) | 1.71 (3H, s) | 1.85 (3H, s) | 1.85 (3H, s) | 1.85 (3H, s) |
| 27 | 1.71 (3H, s) | 1.71 (3H, s) |  |  |  |
| 28 | 1.66 (3H, s) | 1.66 (3H, s) | 1.98 (3H, s) | 1.98 (3H, s) | 1.99 (3H, s) |

a: CDCl<sub>3</sub>, b: CD<sub>3</sub>OD

